# Sex differences in neural representations of social and nonsocial reward in the medial prefrontal cortex

**DOI:** 10.1101/2023.03.09.531947

**Authors:** Jennifer Isaac, Sonia Karkare, Hymavathy Balasubramanian, Nicholas Schappaugh, Jarildy Javier, Maha Rashid, Malavika Murugan

## Abstract

The reinforcing nature of social interactions is necessary for the maintenance of appropriate social behavior. However, the neural substrates underlying social reward processing and how they might differ based on the sex and internal state of the animal remains unknown. It is also unclear whether these neural substrates are shared with those involved in nonsocial rewarding processing. We developed a fully automated, novel two choice (social-sucrose) operant assay in which mice choose between social and nonsocial rewards to directly compare the reward-related behaviors associated with two competing stimuli. We performed cellular resolution calcium imaging of medial prefrontal cortex (mPFC) neurons in male and female mice across varying states of water restriction and social isolation. We found that mPFC neurons maintain largely non-overlapping, flexible representations of social and nonsocial reward that vary with internal state in a sex-dependent manner. Additionally, optogenetic manipulation of mPFC activity during the reward period of the assay disrupted reward-seeking behavior across male and female mice. Thus, using a novel operant assay, we have identified sex-dependent, non-overlapping neural representations of social and nonsocial reward in the mPFC that vary with internal state and that are essential for appropriate reward-seeking behavior.

## Introduction

Animals, including humans, are capable of flexibly choosing between various rewarding stimuli such as food, water, and social interactions. Although considerable progress has been made toward understanding the neural substrates underlying nonsocial reward-related behaviors, less is known about social reward-related behaviors due to the inherent complexity of social interactions^1^. Across species, affiliative social interactions appear to engage the same neural circuits, including the canonical reward-related mesolimbic system, as nonsocial rewards^2–5^. Additionally, social interactions, like nonsocial rewards, can act as positive reinforcers, with rodents returning to the context of a social interaction^6,7^ and engaging in operant tasks for access to a conspecific^8–12^. Despite these commonalities, it is unclear if social and nonsocial reward-related information is encoded by the same or distinct populations of neurons in various brain regions. Some lines of evidence support the common currency view^13,14^, which proposes that the same reward circuitry is used to process both social and nonsocial rewards. For example, neurons in the primate amygdala respond similarly to both juice and pictures of monkeys^15^. In contrast, other evidence supports social specificity, the idea that social stimuli are processed by distinct populations of neurons in the brain^16–18^. Thus, it remains unresolved whether social and nonsocial reward-related behaviors are mediated by shared or separate neuronal populations, specifically in the medial prefrontal cortex (mPFC).

The mPFC is a central node in an interconnected network of brain regions that has been implicated in both nonsocial reward processing^19–24^ and a wide range of social behavior^25–30^. While the role of the mPFC has been extensively studied in the context of nonsocial reward-related behaviors^19–24^, less is known regarding its contribution to social reward-seeking. In rodents, recent work has shown that mPFC neurons are active in the proximity of conspecifics and show increased synchrony when mice engage in affiliative social interactions^25,29,31^. Furthermore, mPFC dysfunction is thought to underlie the social behavioral deficits associated with Autism Spectrum Disorders (ASD)^32–34^. These social behavioral deficits are often accompanied by deficits in the evaluation and processing of nonsocial reward-related stimuli^35–37^. Consequently, determining whether social and nonsocial rewards share neural substrates within the mPFC has important implications for improving therapeutic options for individuals with ASD.

The internal state of an animal also plays a large role in determining its motivation to engage in certain rewarding behaviors and the perceived value of the chosen reward^38–40^. While there is emerging literature to suggest that brain wide changes in neural dynamics result from changes in internal state through different types of deprivation such as thirst^41^ and social isolation^42^, how the internal state of an animal modulates reward representations in the mPFC is poorly understood. Despite there being evidence that sex differences may contribute to how animals engage with and process rewarding stimuli, sex is an often overlooked component when determining how an animal’s internal state modulates reward-related behaviors. In fact, recent studies have shown that male and female mice exhibit divergent learning and decision-making strategies in reward-guided behaviors^22,43,44^. Moreover, sex differences in mPFC reward representations were shown to underlie some of the behavioral differences observed in nonsocial reward-seeking^45^. However, it is unknown if these sex differences extend to social reward-related behaviors. Additionally, even though studies have shown differential impacts of thirst and social isolation on reward-related behaviors in male versus female mice, it is unclear if there are sex differences in how reward representations are affected by varying internal state.

To address these questions, we developed a novel two choice (social-sucrose) operant assay in which mice can freely choose between social and sucrose (nonsocial) rewards in order to directly compare these two types of reward-related behavior. We found that, under control conditions (no social, food or water deprivation), both male and female mice showed a comparable preference for social and sucrose rewards on the two choice assay, with female mice preferring social reward slightly more than sucrose reward. We then sought to determine if distinct ensembles of neurons in the mPFC encode social versus nonsocial rewards. Using cellular resolution calcium imaging, we identified largely non-overlapping ensembles of mPFC neurons that responded to social versus sucrose reward. While the neurons that responded to social reward displayed mostly excitatory responses, the neurons modulated by sucrose reward exhibited both inhibitory and excitatory responses. Interestingly, the neurons that displayed an inhibitory response to sucrose reward were also excited in response to social reward. The social and sucrose reward ensembles were less overlapping in female mice compared to male mice. After characterizing these distinct social and nonsocial reward representations in the mPFC, we optogenetically manipulated mPFC neurons during the reward period of the two choice operant assay and found that it disrupted reward-seeking behavior in both male and female mice. We then examined how varying the internal state of the animal affected reward representations in the mPFC by either water depriving or socially isolating mice prior to imaging them on the two choice operant assay. Water-deprived mice of both sexes showed increased sucrose reward-seeking behavior and a greater proportion of mPFC neurons that were modulated by sucrose reward relative to control conditions. By tracking the same neurons across imaging sessions, we found that the newly recruited sucrose reward-responsive neurons were largely derived from previously latent, reward-unresponsive neurons. We also found that acute social isolation differentially affected reward-seeking behavior in male and female mice. However, it did not affect the proportion of mPFC neurons that were modulated by social reward. Instead, social isolation altered the amplitude of the neural responses to social but not sucrose reward in a sex-dependent manner, with increased amplitude in socially-isolated female mice and decreased amplitude in socially-isolated male mice relative to control conditions. Overall, these findings suggest that although male and female mice behave similarly while flexibly choosing between social and nonsocial rewards on a novel two choice operant assay, there are sex differences in how social and nonsocial rewards are represented and in how changing internal state affects these representations in the mPFC.

## Results

### Male and female mice demonstrate positive reinforcement for social and sucrose reward stimuli

In order to directly compare nonsocial and social reward-related behavior and the neural circuits underlying these behaviors, we developed a self-paced, automated two choice (social-sucrose) operant assay in which mice were trained to associate two separate choice ports with access to either a sucrose reward or a same sex conspecific (social reward) over one hour daily sessions (Figure 1a-c).

**Figure 1.**
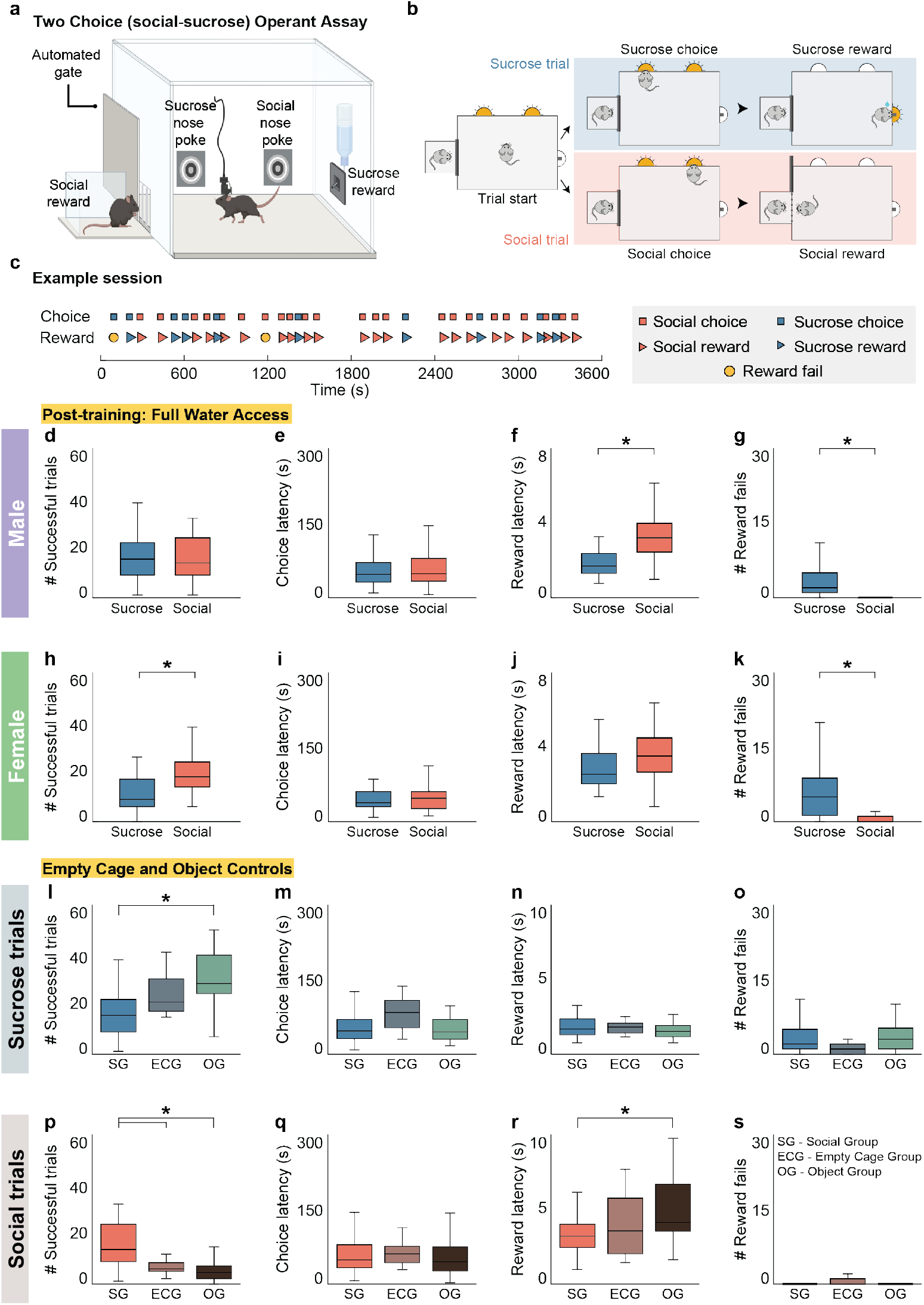
Male and female mice find both social and sucrose stimuli positively reinforcing on a two choice (social-sucrose) operant assay. **a,b**) Schematic of a novel automated two choice (social-sucrose) operant assay in which a mouse can freely choose (nose-poke) to obtain either a sucrose (right, top blue panel) or social (right, bottom red panel) reward. c) Timeline of an example behavioral session showing social (red squares) and sucrose (blue squares) choices followed by the respective reward consumption (red or blue arrowheads) over an hour. Reward fails (yellow circles) are defined as failure of a mouse to enter the chosen reward zone while the reward is available for consumption. d,h) Fully trained male mice (**d**) complete an equivalent number of successful sucrose and social trials, while female mice (**h**) complete significantly more successful social than sucrose trials. Paired two-tailed t test (male: number of trials: p = 0.73; female: p = 4.91*10-4). e,i) Male and female mice (**e**,**i**) show a similar choice latency for sucrose and social choices. Paired two-tailed t test (male: p = 0.12; female: p = 0.93). f,j) Male mice (**f**) were slightly faster to consume sucrose reward than social reward, while female mice (**j**) had equivalent latencies to consume social and sucrose reward. Paired two-tailed t test (male: p = 2.04*10-8; female: p = 0.093). g,k) Male (**g**) and female (**k**) mice made fewer social reward fails than sucrose reward fails. Paired two-tailed t test (male: p = 1.00*10-7; female: p = 4.10*10-6). N = 21 male mice, 12 female mice, 3 behavioral sessions per mouse. l) Under full water access conditions, mice run on the two choice operant assay with a social target (SG) completed significantly fewer successful sucrose trials when compared to mice from the object group (OG) and a similar number of successful sucrose trials when compared to mice from the empty cage group (ECG). One-way ANOVA (p = 1.34*10-8) with post-hoc t tests (SG vs OG: p = 8.44*10-9; SG vs ECG: p = 0.052; OG vs ECG: p = 0.10). m-o) There was no difference in sucrose choice latency (**m**), sucrose reward latency (**n**) or number of sucrose reward fails (**o**) between groups. One-way ANOVA (**m**, p = 0.080; **n**, p = 0.19; **o**, p = 0.078). p) Under full water access conditions, mice run on the two choice assay with a social target (SG) completed more social trials than mice run with an empty cage (ECG) or an object (OG). One-way ANOVA (p = 3.30*10-10) with post-hoc t tests (SG vs OG: p = 2.63*10-9; SG vs ECG: p =1.17*10-4; OG vs ECG: p = 0.87). q,s) Across all conditions, mice showed similar social choice latency (**q**) and number of social reward fails (**s**). One-way ANOVA (**q**, p = 0.92; **s**, p = 0.44). r) Mice run with a social target (SG) showed similar social reward latency to mice run with an empty cage (ECG) and decreased social reward latency compared to mice run with an object (OG). One-way ANOVA (p = 2.00*10-4) with post-hoc t tests (SG vs OG: p = 1.04*10-4; SG vs ECG: p = 0.46; OG vs ECG: p = 0.19). SG: n = 21 mice, ECG: n = 4 mice, OG: n = 10 mice, 3 sessions per mouse. Box plots: center line denotes median, box edges indicate the 25th and 75th percentiles and whiskers extend to ± 2.7σ. *p<0.05.

Initially, mice on restricted water are trained to associate nose-poking a choice port with sucrose reward delivery at a reward port located on the adjacent wall of the operant chamber through a series of increasingly difficult assays (Supplementary Figure 1 and see Methods). Mice showed a consistent decrease in the time from nose-poking the choice port to accessing the reward port (sucrose reward latency) with training (Supplementary Figure 1h: one-way ANOVA with post-hoc t tests, p = 4.01*10-8). During the final training stage, a social component was added to the assay to allow mice to choose between a social and sucrose reward (Supplementary Figure 1a, Supplementary Figure 2). At the start of each trial on the two choice operant assay, two choice ports on one wall of the operant chamber illuminate (Figure 1b, trial start). Nose-poking one of the illuminated choice ports gives mice access to 10μl of 10% sucrose at the sucrose reward port for up to 8 seconds (Figure 1b, sucrose trial, blue panel), while nose-poking the other illuminated choice port opens an automatic gate to give mice access to a novel, same sex conspecific (Figure 1b, social trial, red panel). Both male and female mice showed a decrease in their social reward latency across five training sessions on the two choice operant assay, as mice learned to associate the social choice port with a social reward (Supplementary Figure 2g,o: one-way ANOVA with post-hoc t tests, male: p = 8.19*10-6, female: p = 0.016). Male mice also showed a decrease in social reward fails, defined as failure to enter the social zone when the gate is open, with training (Supplementary Figure 2h: one-way ANOVA with post-hoc t tests, p = 0.0011). Additionally, both sexes showed a stable number of successful social trials across five training sessions (Supplementary Figure 2e,m: one-way ANOVA with post-hoc t tests, male: p = 0.11, female: p = 0.98).

After five training sessions on the two choice operant assay, mice were removed from restricted water access and run for an additional five sessions (post-training, full water access). Post-training, we found that male mice in control conditions (not water, food or socially deprived) completed an equivalent number of successful sucrose and social trials (Figure 1d: paired t test, p = 0.73), while female mice completed slightly more successful social than sucrose trials (Figure 1h: paired t test, p = 4.91*10-4) at comparable choice latencies (Figure 1e,i: paired t test, male: p = 0.12, female: p = 0.93). Male mice were slightly slower to engage with social compared to sucrose reward (Figure 1f, paired t test, p = 2.04*10-8), likely reflecting their longer exposure to sucrose reward across all training assays. In contrast, female mice demonstrated equivalent reward latencies for both trial types (Figure 1j: paired t test, p = 0.093). Male and female mice made more sucrose reward fails than social reward fails (Figure 1g,k: paired t test, male: p = 1.0*10-7, female: p = 4.10*10-6).

Although female mice completed slightly more successful social trials and slightly fewer successful sucrose trials than male mice (Supplementary Figure 3i: two-factor ANOVA with sex (male/female) and trial type (sucrose/social) as factors, interaction: p = 8.00*10-4, sex: p = 0.78, trial type: p = 0.0035, with post-hoc unpaired t tests comparing sex within trial type, sucrose: p = 0.011, social: p = 0.029), both sexes completed significantly more social and significantly fewer sucrose trials when compared to partial water access conditions during training (Supplementary Figure 3a,e: two-factor ANOVA with water condition (PW/FW) and trial type (sucrose/social) as factors; male - interaction: p = 2.63*10-26, water condition: p = 9.54*10-13, trial type: p = 1.22*10-27, with post-hoc unpaired t tests comparing water condition within trial type, sucrose: p = 1.04*10-20, social: p = 2.32*10-5; female - interaction: p = 4.93*10-20, water condition: p = 5.76*10-6, trial type: p = 3.02*10-9, with post-hoc unpaired t tests comparing water condition within trial type, sucrose: p = 4.49*10-13, social: p = 4.14*10-8). Additionally, male mice displayed shorter reward latencies to consume sucrose reward and made fewer sucrose reward fails compared to female mice (Supplementary Figure 3k,l: two-factor ANOVA with sex (male/female) and trial type (sucrose/social) as factors, reward latency - interaction: p = 0.032, sex: p = 4.00*10-4, trial type: p = 1.64*10-7, with post-hoc unpaired t tests comparing sex within trial type, sucrose: p = 1.29*10-5, social: p = 0.35; reward fails - interaction: p = 0.010, sex: p = 0.0048, trial type: p = 2.79*10-15) with post-hoc unpaired t tests comparing sex within trial type, sucrose: p = 0.0073, social: p = 0.28), male and female mice otherwise performed similarly when seeking social rewards on the two choice operant assay (Supplementary Figure 3i-l). Furthermore, female reward-seeking behavior on the two choice operant assay was not affected by estrous cycle (Supplementary Figure 6a,b) or mouse strain (Supplementary Figure 6c-f). These findings suggest that when male and female mice have ad libitum access to water, they are motivated to seek both social and nonsocial rewards (Figure 1, Supplementary Figure 3).

Importantly, mice trained on the two choice operant assay without a social target present (empty cage group, ECG) or with a novel object instead of a social target (object group, OG) significantly decreased the number of successful empty cage/object trials, while increasing the number of sucrose trials over five consecutive sessions (Supplementary Figure 4b,f: one-way ANOVA with post-hoc t tests, number of sucrose trials - ECG: p = 0.01, OG: p = 1.25*10-7; number of social trials - ECG: p = 5.51*10-6, OG: p = 3.0*10-4). When these mice had ad libitum water access, they continued to complete significantly fewer social, but not sucrose, trials when there was no social target (empty cage or object) compared to mice run on the two choice operant assay with a social target (Figure 1l,p: one-way ANOVA with post-hoc t tests, number of sucrose trials: p = 1.34*10-8; number of social trials: p = 3.30*10-10), which indicates that social reward-seeking behavior is positively reinforced by the presence of a social target and not solely by the gate opening or novelty-seeking behavior (Figure 1l-s, Supplementary Figure 4). To further confirm the goal-directed nature of the social reward-seeking behavior observed on the novel two choice operant assay, we developed an additional multi-choice assay in which a third choice port was introduced that was not associated with any reward (Supplementary Figure 5a,b). Mice run on this assay preferentially increased the number of successful social but not null trials completed over time and were significantly faster to enter the corresponding reward zone on sucrose and social trials but not null trials (Supplementary Figure 5c: two-factor ANOVA with training day (day 1/day 9) and trial type (sucrose/social/null) as factors, interaction: p = 0.0059, training day: p = 0.68, trial type: p = 5.02*10-13, with post-hoc unpaired t tests comparing trial type within training day, day 1 - suc v soc: p = 1.02*10-4, suc v null: p = 1.45*10-4, soc v null: p = 0.48, day 9 - suc v soc: p 0.078, suc v null: p = 5.21*10-4, soc v null: p = 0.013; Supplementary Figure 5d: two-factor ANOVA with choice type (sucrose/social/null) and reward zone (sucrose/social) as factors, interaction: p = 2.44*10-13, choice type: p = 0.41, reward zone: p = 0.10, with post-hoc unpaired t tests comparing reward zone within choice type, sucrose: p = 2.19*10-8, social: p = 1.82*10-5, null: p = 0.41). These findings are among the first to demonstrate that social interaction can promote positive reinforcement of reward-seeking behavior similarly to sucrose consumption in an operant assay.

### Cellular resolution imaging of mPFC neurons in male and female mice during the two choice operant assay

To determine what information mPFC neurons encode during social and nonsocial reward-related behavior, we performed cellular resolution calcium imaging of mPFC neurons during the two choice operant assay. We injected an AAV expressing the calcium indicator GCaMP6f and implanted a gradient-index (GRIN) lens into the mPFC of 9 male and 6 female mice (Figure 2a). All lens-implanted mice were then successfully trained on the two choice operant assay. After three weeks of training, the lens was coupled to a head-mounted miniscope to record activity of GCaMP6f-expressing mPFC neurons while mice completed the two choice operant assay. CNMFe was used to detect the activity of individual neurons^46^. Post-training, in full water access conditions, lens-implanted imaging mice were run on a minimum of 3 daily sessions prior to imaging mPFC neural activity in the two choice operant assay. We imaged a total of 459 neurons from 9 male mice and 570 neurons from 6 female mice in control conditions (Figure 2b,d). GRIN lens placement and viral expression were confirmed on post-hoc histology (Figure 2c,e).

**Figure 2.**
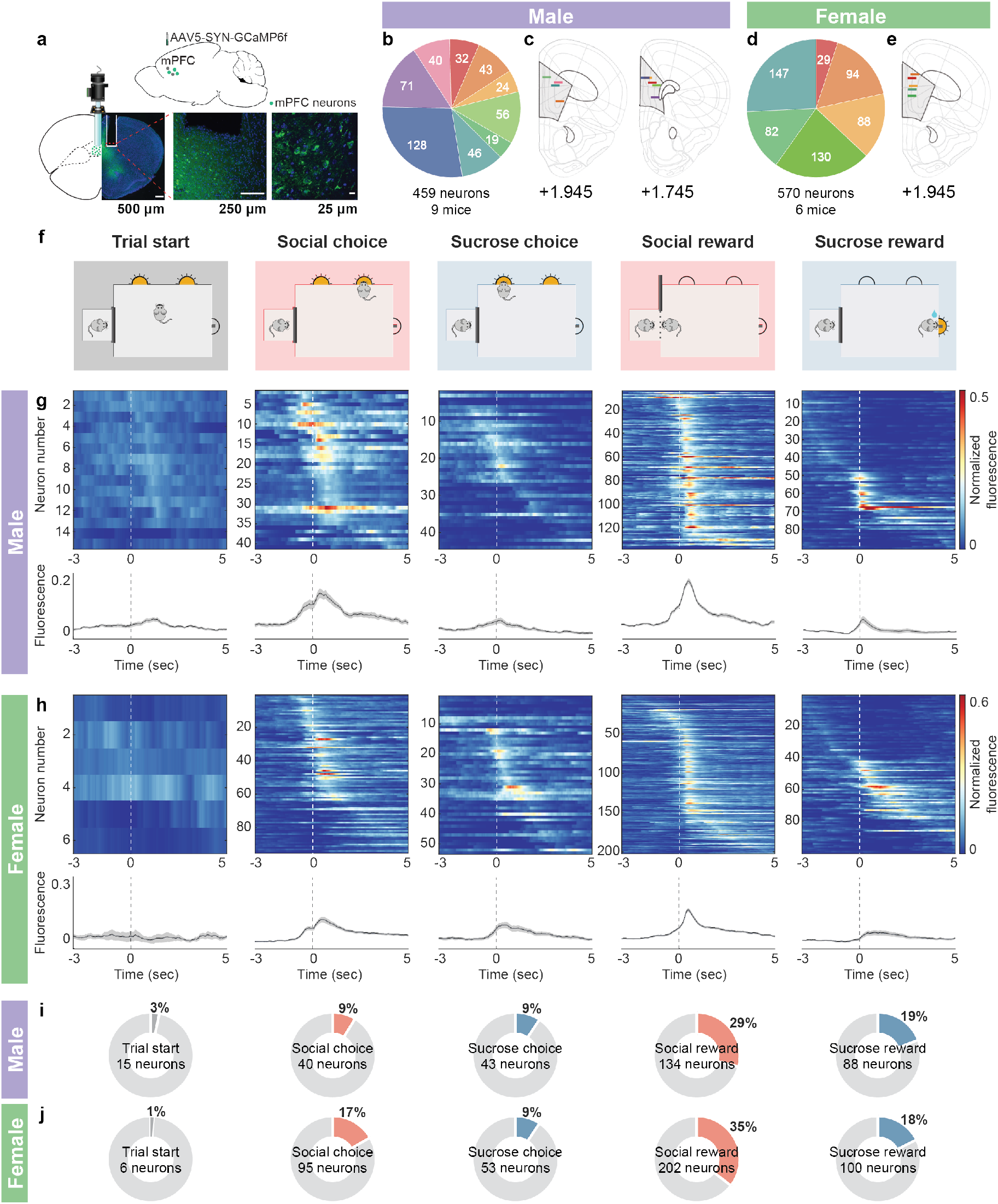
Cellular resolution calcium imaging of mPFC neurons during the two choice operant assay. **a**) Top row: Schematic of the viral strategy used to label mPFC neurons with GCaMP6f. Bottom row: Imaging setup of GRIN lens placement in the mPFC, including representative images showing GRIN lens placement (left), GCaMP6f expression (middle, GCaMP6f in green; DAPI in blue) in the mPFC at (from left to right) 4x, 20x and 60x magnification. Right panel shows nuclear exclusion of GCaMP6f. Scale bars: 500 µm, 250 µm and 25 µm. b,d) A pie chart showing the number of neurons recorded from each male (**b**, n = 459 neurons, 9 mice) and female mouse (**d**, n = 570 neurons, 6 mice). c,e) Reconstruction of GRIN lens placement in the nine male (**c**) and six female (**e**) mice using WholeBrain. Each colored line corresponds to the same colored slice in the pie chart and shows the position of the lens in the Allen Mouse Brain Common Coordinate Framework. Coordinates are relative to bregma. f) A schematic of the two choice operant assay events to which mPFC neuronal activity was aligned, from left to right, trial start, social choice, sucrose choice, social reward and sucrose reward. g,h) Top row: Heatmaps of average normalized fluorescence of all mPFC neurons that are significantly modulated (excited or inhibited) by each task event in male (**g**) and female (**h**) mice. Neurons are sorted by the time of maximum fluorescence across each task event. Bottom row: Average normalized fluorescence traces of the neurons from the corresponding heatmap that are significantly modulated by each task event. Shaded error regions indicate ± SEM. i,j) Proportions of total recorded neurons that are modulated by the various task events in male (**i**) and female (**j**) mice (male: n = 459 neurons, 9 mice; female: n = 570 neurons, 6 mice).

Neurons were classified as task-modulated if the maximum or minimum of their average activity differed from a null distribution generated by randomly shuffling neural activity (see Methods, example task-modulated neurons Supplementary Figure 7a) within a three second window around the occurrence of a task event (trial start, social or sucrose choice, social or sucrose reward, Figure 2f)^20^. We found that many mPFC neurons showed significantly modulated time-locked responses to specific behavioral events in the two choice operant assay (Figure 2g,h, Supplementary Figure 7b-e). In fact, across all mPFC neurons recorded in male mice during the two choice operant task, 3.27% (n = 15/459) were significantly modulated by trial start, 8.71% (n = 40/459) were significantly modulated by social choice, 9.37% (n = 43/459) were significantly modulated by sucrose choice, 29.19% (n = 134/459) were significantly modulated by social reward and 19.17% (n = 88/459) were significantly modulated by sucrose reward (Figure 2i). Across all mPFC neurons recorded from female mice during the two choice operant task, 1.05% (n = 6/570) were significantly modulated by trial start, 16.67% (n = 95/570) were significantly modulated by social choice, 9.30% (n = 53/570) were significantly modulated by sucrose choice, 35.44% (n = 202/570) were significantly modulated by social reward and 17.54% (n = 100/570) were significantly modulated by sucrose reward (Figure 2j). We found similar fractions of task-modulated neurons when using an encoding model to temporally separate calcium activity kernels associated with various behavioral events (Supplementary Figure 8; proportion z test, male - choice: p = 0.86, reward: p = 0.39; female - choice: p = 0.12, reward: p = 0.15). Previous literature has shown that mPFC neurons respond more to action (i.e., nose-poke, reward consumption) than stimulus (i.e., trial start) events^21^. Consistent with these findings, we found that a greater proportion of mPFC neurons responded to choice and reward consumption than to trial start (Supplementary Figure 9a: male - stimulus events: 3.27%,15/459, action events: 66.45%, 305/459; female - stimulus events: 1.05%, 6/570, action events: 78.95%, 450/570; proportion z test, male and female: p<0.00001). Additionally, we found that more mPFC neurons were modulated by reward than choice (Supplementary Figure 9b,c: male - choice: 18.08%, 83/459, reward: 48.37%, 222/459; female - choice: 25.96%, 148/570, reward: 52.98%, 302/570; proportion z test, male and female: p<0.00001).

### Sex differences in selectivity of social and nonsocial reward-related neural representations

Given the well-defined role of the mPFC in decision-making^47,48^, we next examined the responses of choice-modulated neurons during the two choice assay. We found that choice-modulated neurons reliably showed peak fluorescence around the time of choice port entry and that the timing of peak fluorescence did not vary with reward latency (Figure 3a,c). In contrast, the timing of the peak fluorescence of reward-modulated neurons varied with reward latency when aligned to choice port entry (Figure 3b,d). Furthermore, we found that a similar proportion of neurons responded to social (n = 8.71%, 40/459) and sucrose (n = 9.37%, 43/459) choice in male mice (Figure 2i, proportion z test, p = 0.73), while a significantly greater proportion of neurons responded to social choice (n = 16.67%, 95/570) than sucrose choice (n = 9.30%, 53/570) in female mice (Figure 2j, proportion z test, p = 2.15*10-4). The neurons that were excited by sucrose choice were more selective for sucrose choice in female mice (Supplementary Figure 9h: paired t test, p = 3.61*10-6) relative to male mice (Supplementary Figure 9e: paired t test, p = 0.45), while neurons that were excited by social choice (Supplementary Figure 9d,g) as well as those that were inhibited by sucrose choice (Supplementary Figure 9f,i) showed similar responses across both sexes. These findings suggest that mPFC neurons differentially represent social and sucrose choice in male versus female mice.

**Figure 3.**
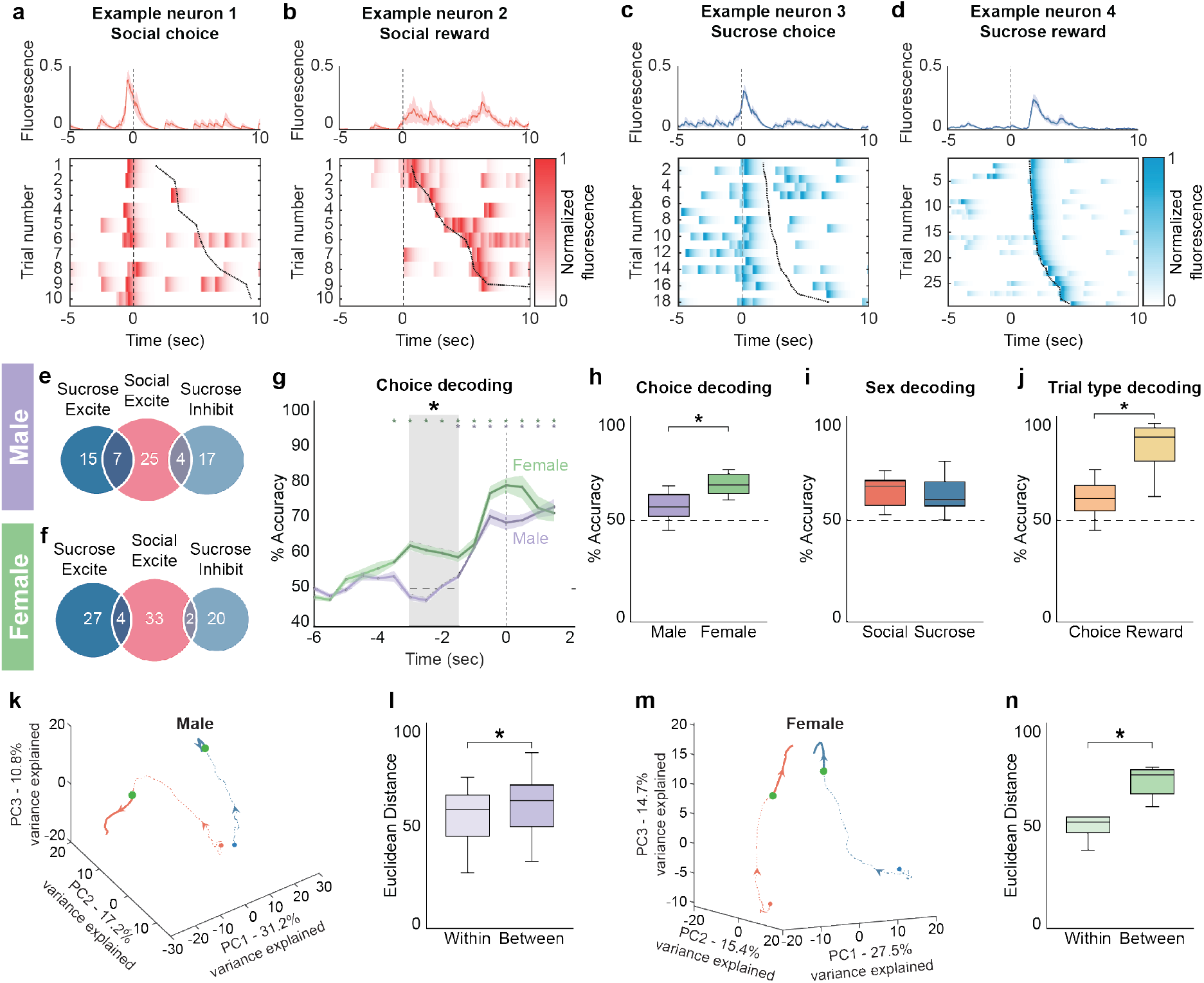
Sex differences in selectivity of social and nonsocial neural representations of choice. **a,b**) Normalized fluorescence of an example neuron significantly modulated by social choice aligned to social choice (**a**) compared to the normalized fluorescence of an example neuron significantly modulated by social reward aligned to social choice (**b**). c,d) Normalized fluorescence of an example neuron significantly modulated by sucrose choice aligned to sucrose choice (**c**) compared to the normalized fluorescence of an example neuron significantly modulated by sucrose reward aligned to sucrose choice (**d**). Top row shows average fluorescence ± SEM (dashed line at zero indicates choice onset). Bottom row shows a heatmap of normalized fluorescence on each trial aligned to choice (dashed line at zero indicates choice onset) and sorted by reward latency (dark dotted line indicates sucrose reward start). e,f) Venn diagrams showing that mPFC choice neurons are largely non-overlapping in their responsiveness to social and sucrose choice in both male (**e**) and female (**f**) mice. Choice neurons are largely exclusive in their response to either social or sucrose choice (male: 83.82%, n = 57/68; female: 93.02%, n = 80/86). g) Average choice decoding accuracy as a function of time relative to choice. mPFC population activity accurately decoded the subsequent choice earlier in female (green) mice relative to male (purple) mice indicated by shaded gray region. Unpaired t test (−3.0 to − 2.5s: p = 0.0027, −2.5 to −2.0s: p = 0.0032, −2.0 to −1.5s: p = 0.019). Choice decoding accuracy in female and male mice becomes equivalent at 1.5s before choice port entry and both are greater than chance decoding accuracy (colored asterisks and bolded lines indicate where choice decoding is significantly greater than chance). Shaded error regions indicate ± SEM. Dashed line at zero indicates choice port entry. h) mPFC population activity in female mice (n = 5 mice) more accurately decoded the choice made on a particular trial compared to male mice (n = 9 mice). Decoding accuracy was calculated for each animal from all recorded neurons, with a trial-matched number of sucrose and social trials. Unpaired t test (p = 0.0090). i) mPFC population activity in a 3s window around social and sucrose choice was equivalently sufficient to decode the sex of the animal and decoded with significantly higher accuracy than shuffled data (dotted line) on both trial types. Unpaired t test (social versus sucrose: p = 0.19; social versus shuffled: p = 2.08*10-19; sucrose versus shuffled: p = 6.84*10-12). j) Across all mice, mPFC neural representations resulted in greater decoding accuracy for reward compared to choice. Paired t test (p = 7.90*10-8). k,m) Trial-averaged population neural activity traces of sucrose (blue) and social (red) trials in male (**k**, n = 459 neurons, 9 mice) and female (**m**, n = 423 neurons, 5 mice) mice plotted on the first 3 PCs in state space. Arrowhead indicates direction of time. Filled green circle indicates choice onset. l,n) Euclidean distance separating PC-projected population vectors in a 3s window around choice is significantly greater between social and sucrose trials than within each trial type in both male (**l**) and female (**n**) mice. Paired t test (male: p = 5.76*10-4; female: p = 0.0048). All decoding was significantly greater than shuffled data, indicated by a dashed line at 50%. Box plot: center line denotes median, box edges indicate the 25th and 75th percentiles and whiskers extend to ± 2.7σ. *p<0.05.

Interestingly, although the neural representations of social and sucrose choice in the mPFC neurons were largely non-overlapping in both male and female mice (Figure 3e,f: % non-overlapping, male: n = 83.82%, 57/68; female: n = 93.02%, 80/86, proportion z test, p = 0.07), we were able to decode subsequent choice at an earlier time point in female mice compared to male mice (Figure 3g: female: 3.5s, male: 1.5s prior to choice). We were also able to decode the choice made by the mouse from mPFC population activity with higher accuracy in female mice compared to male mice (Figure 3h: average decoding accuracy, male: 56.09 ± 2.23, female: 67.03 ± 2.43; unpaired t test, p = 0.0090). Additionally, we were able to decode the sex of the animal from mPFC population activity associated with choice on both trial types. (Figure 3i: average decoding accuracy, social choice: 63.70 ± 1.026, sucrose choice: 61.68 ± 1.13; unpaired t test, p = 0.19). The largely non-overlapping nature of the choice responses was corroborated by plotting the neural trajectories of mPFC population activity on the first 3 principal components (PCs) in a 3 second window around choice port entry in both male and female mice (Figure 3k,m: total variance explained, male: 59.2%, female: 57.6%) and calculating pairwise Euclidean distances of population vectors within trial type versus between trial type (Figure 3l,n). In both sexes, we found that the Euclidean distance within trial type was significantly less than the distance between trial type across social and sucrose trials (Figure 3l,n: male - average distance within: 54.71 ± 4.76, between: 60.87 ± 5.13; paired t test, p = 5.76*10-4; female - average distance within: 49.89 ± 2.55, between: 71.93 ± 2.99; paired t test, p = 0.0048). Thus, population activity even at a single-trial level showed separable subspace representation for social and nonsocial trials. These findings suggest that mPFC neural activity associated with choice port entry is distinct for social and nonsocial trials despite the mice engaging in similar behaviors (nose-poking a choice port). Consequently, neural representations of social and nonsocial reward-related behaviors are highly selective and sex-dependent even prior to reward consumption.

Since we found that the largest proportion of mPFC neurons were modulated by reward (Supplementary Figure 9b,c) and that trial type decoding was greater for reward than choice in male and female mice (Figure 3j: average decoding accuracy, choice: 60.0 ± 2.18, reward: 86.38 ± 2.78; paired t test, p = 7.90*10-8), we further characterized the mPFC reward responses. In both sexes, we found that the majority of neurons that were responsive to social reward were positively modulated or excited by social reward (Figure 4a, male: 96.27%, n = 129/134; female: 87.62%, n = 177/202), compared to those that were negatively modulated or inhibited by social reward (Figure 4a, male: 3.73%, n = 5/134; female: 12.38%, n = 25/202; proportion z test, male and female: p<0.00001). In contrast, in all mice, more neurons were inhibited by sucrose reward (Figure 4b, male: 64.77%, n = 57/88; female: 60.0%, n = 60/100) than excited by sucrose reward (Figure 4b, male: 35.23%, n = 31/88; female: 40.0%, n = 40/100; proportion z test, male: p = 8.87*10-5; female: p = 0.0047). Thus, it appears that mPFC neurons differ in their response to social and nonsocial rewards, with a largely excitatory response to social reward and a mixed inhibitory and excitatory response to sucrose reward.

**Figure 4.**
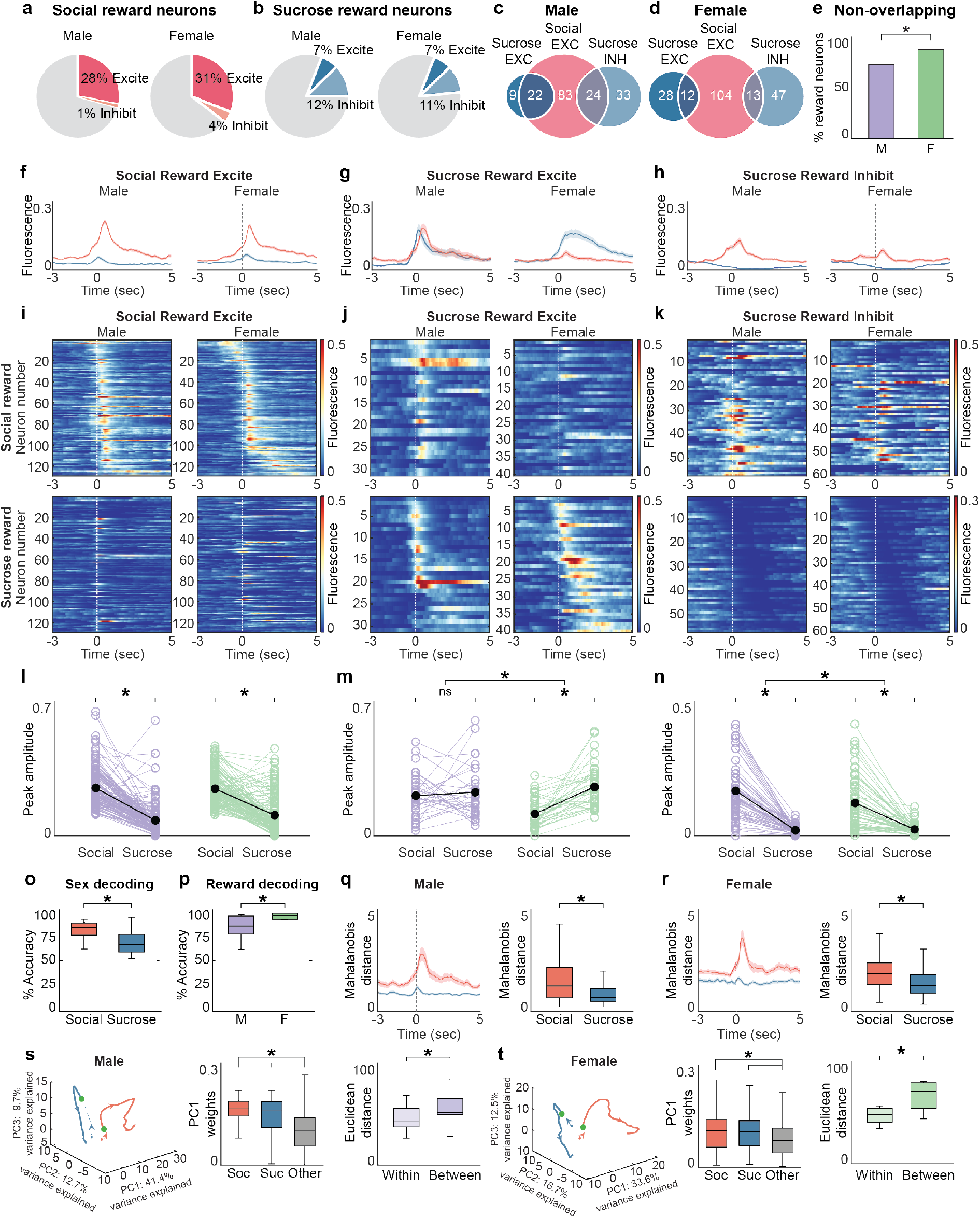
mPFC neurons differ in social and nonsocial reward response selectivity in a sex-dependent manner. **a**) Pie charts showing the distribution of social reward responses in male (left) and female (right) mice. In both male and female mice, a larger proportion of mPFC neurons are positively modulated (excited) in response to social reward (male: 28.11%, n = 129/459; female: 31.10%, n = 177/570) compared to those that are negatively modulated (inhibited) by social reward (male: 1.09%, n = 5/459; female: 4.39%, n = 25/570). Proportion z test (male: p<0.00001; female: p<0.00001). b) Pie charts showing the distribution of sucrose reward responses in male (left) and female (right) mice. In contrast to social reward responses, mPFC neurons in male and female mice are more likely to be inhibited (male: 12.42%, n = 57/459; female: 10.53%, n = 60/570) rather than excited (male: 6.75%, n = 31/459; female: 7.02%, n = 40/570) in response to sucrose reward. Proportion z test (male: p = 0.0036; female: p = 0.036). c) A venn diagram showing that mPFC reward neurons differ in the selectivity of their reward responsiveness in male mice. Social excite neurons are largely exclusive in their response to social reward (64.34%, n = 83/129), with a significantly smaller subset of social excite neurons also responding to sucrose reward (35.66%, 46/129). Proportion z test (p = 4.08*10-6). In contrast, most sucrose excite neurons also responded to social reward (70.97%, n = 22/31). Around half of sucrose inhibit neurons also responded to social reward (42.11%, n = 24/57). Proportion z test (sucrose excite: p = 9.60*10-4, sucrose inhibit: p = 0.092). d) A venn diagram showing that mPFC reward neurons are largely non-overlapping in their reward responsiveness in female mice. Social excite neurons are largely exclusive in their response to social reward (80.62%, n = 104/129), with a significantly smaller subset of social excite neurons also responding to sucrose reward (19.38%, n = 25/129). Proportion z test (p<0.00001). In addition, the majority of sucrose excite neurons did not respond to social reward (70.0%, n = 28/40) and the majority of sucrose inhibit neurons did not respond to social reward (78.3%, n = 47/60). Proportion z test (sucrose excite: p = 3.45*10-4, sucrose inhibit: p = 5.38*10-10). e) Largely non-overlapping populations of mPFC neurons respond to social and sucrose reward in both male (n = 73.10%, 125/171) and female (n = 87.75%, 179/204) mice. The populations are more distinct in female mice relative to male mice. Proportion z test (p = 3.11*10-4). f) Average responses of social reward excite neurons to social (red) and sucrose (blue) reward in male (left) and female (right) mice. g) Average responses of sucrose reward excite neurons to social (red) and sucrose (blue) reward in male (left) and female (right) mice. h) Average responses of sucrose reward inhibit neurons to social (red) and sucrose (blue) reward in male (left) and female (right) mice. i) Heatmaps of the average fluorescence of mPFC neurons in male (left, n = 129 neurons) and female (right, n = 129 neurons) mice that are significantly excited by social reward aligned to social reward (top row) and sucrose reward (bottom row). Neurons are sorted by the time of peak response to social reward. j) Heatmaps of the average fluorescence of mPFC neurons in male (left, n = 31 neurons) and female (right, n = 40 neurons) mice that are significantly excited by sucrose reward aligned to social reward (top row) and sucrose reward (bottom row). Neurons are sorted by the time of peak response to sucrose reward. k) Heatmaps of the average fluorescence of mPFC neurons in male (left, n = 57 neurons) and female (right, n = 60 neurons) mice that are significantly inhibited by sucrose reward aligned to social reward (top row) and sucrose reward (bottom row). Neurons are sorted by the time of minimum response to sucrose reward. A row on the top and bottom panels of each heatmap (**i**,**j**,**k**) corresponds to the same neuron. l) Comparison of the peak amplitude of responses of social reward excite neurons to social and sucrose reward in male (purple, n = 129 neurons, 9 mice) and female (green, n = 129 neurons, 5 mice) mice shows that these neurons on average have a robust excitatory response to social but not sucrose reward in both sexes. Two-factor ANOVA with sex and trial type as factors (interaction: p = 0.067, sex: p = 0.17, trial type: p = 4.11*10-58) with post-hoc t tests comparing trial type within sex (male: p = 1.20*10-33, female: p = 4.15*10-26). m) Comparison of the peak amplitude of responses of sucrose reward excite neurons to social and sucrose reward in male (purple, n = 31 neurons, 9 mice) and female (green, n = 40 neurons, 5 mice) mice shows that sucrose reward excite neurons are selective for sucrose reward in female but not male mice. Two-factor ANOVA with sex and trial type as factors (interaction: p = 0.0042, sex: p = 0.014, trial type: p = 1.87*10-4) with post-hoc t tests comparing trial type within sex (male: p = 0.58, female: p = 4.79*10-9). n) Comparison of the peak amplitude of responses of sucrose reward inhibit neurons to social and sucrose reward in male (purple, n = 57 neurons, 9 mice) and female (green, n = 60 neurons, 5 mice) mice shows that these neurons on average have a higher amplitude response to social reward compared to sucrose reward across both sexes. Two-factor ANOVA with sex and trial type as factors (interaction: p = 0.011, sex: p<0.00001, trial type: p = 0.057) with post-hoc t tests comparing trial type within sex (male: p = 5.48*10-18, female: p = 7.60*10-12). o) mPFC population activity in a 3s window around reward was able to decode the sex of the animal with higher accuracy on social than on sucrose trials. Unpaired t test (p = 1.70*10-6). p) Decoders trained on female mPFC neural reward responses decoded the reward type with higher accuracy than decoders trained on male reward responses. Unpaired t test (p = 0.022). Decoding accuracy was calculated for each animal from all recorded neurons, with a trial-matched number of sucrose and social trials. All decoding was significantly greater than shuffled data, indicated by a dashed line at 50%. q,r) Mahalanobis distance was greater for social than sucrose reward in male (**q**) and female (**r**) mice. Unpaired t test (male: p = 1.71*10-8; female: p = 2.07*10-5). s,t) Trial-averaged population neural activity traces of sucrose (blue) and social (red) reward trials in male (**s**, left panel, n = 459 neurons, 9 mice) and female (**t**, left panel, n = 423 neurons, 5 mice) mice plotted on the first 3 PCs in state space. Arrowhead indicates direction of time. Filled green circle indicates reward onset. PC1 weights are higher for both social and sucrose reward neurons compared to other neurons in both male (**s**, middle panel) and female (**t**, middle panel) mice. One-way ANOVA (male: p = 9.22*10-17; female: p = 2.93*10-8) with post-hoc t tests (male - soc v other: p = 2.46*10-17, suc v other: p = 3.14*10-6, soc v suc: p = 0.076; female - soc v other: p = 6.21*10-7, suc v other: p = 2.11*10-5, soc v suc: p = 0.96). Euclidean distance separating PC-projected population vectors in a 3s window around reward is significantly greater between social and sucrose trials than within each trial type in both male (**s**, right panel) and female (**t**, right panel) mice. Paired t test (male: p = 5.81*10-4; female: p = 0.0020). *p<0.05. Shaded error regions indicate ± SEM. Dashed line at zero indicates reward onset. Box plots: center line denotes median, box edges indicate the 25th and 75th percentiles and whiskers extend to ± 2.7σ.

Next, we examined the selectivity of mPFC reward responses by determining the overlap between these populations. In both male (Figure 4c) and female (Figure 4d) mice, largely non-overlapping populations of mPFC neurons responded to social and sucrose reward (% non-overlapping: male: 73.10%, n = 125/171; female: 87.75%, n = 179/204; unpaired t test comparing neural data to shuffled data, male and female: p<0.00001, see Methods). The social and nonsocial reward ensembles in the mPFC were even more distinct in female mice relative to male mice (Figure 4e: proportion z test, p = 3.11*10-4). Thus, both male and female mice showed largely non-overlapping social and nonsocial reward responses in the mPFC.

We then evaluated how social and sucrose reward-responsive neurons were modulated by the alternative reward stimulus. We compared the social and sucrose reward responses of neurons that were excited by social reward (Figure 4f,i,l), excited by sucrose reward (Figure 4g,j,m) and inhibited by sucrose reward (Figure 4h,k,n). Both social reward excite and sucrose reward inhibit neurons showed significantly higher responses to social reward compared to sucrose reward in male and female mice (Figure 4l,n: two-factor ANOVA with sex (male/female) and trial type (sucrose/social) as factors, social excite - interaction: p = 0.067, sex: p = 0.17, trial type: p = 4.11*10-58, with post-hoc unpaired t tests comparing trial type within sex, male: p = 1.20*10-33, female: p = 4.15*10-26; sucrose inhibit - interaction: p = 0.011, sex: p<0.00001, trial type: p = 0.057, with post-hoc unpaired t tests comparing trial type within sex, male: p = 5.48*10-18, female: p = 7.60*10-12). In contrast, sucrose reward excite neurons showed significantly higher responses to sucrose reward relative to social reward in female mice but not male mice (Figure 4m: two-factor ANOVA with sex (male/female) and trial type (sucrose/social) as factors, interaction: p = 0.0042, sex: p = 0.014, trial type: p = 1.87*10-4 with post-hoc unpaired t tests comparing trial type within sex, male: p = 0.58, female: p = 4.76*10-9). These sex differences in reward selectivity were further supported by evidence that reward decoding accuracy is greater in female mice than in male mice (Figure 4p: average decoding accuracy, male: 82.54 ± 3.71, female: 93.29 ± 1.47; unpaired t test, p = 0.022). Additionally, mPFC activity during the reward period could decode animal sex with greater accuracy on social trials compared to sucrose trials (Figure 4o: average decoding accuracy, social: 79.45 ± 1.34, sucrose: 68.10 ± 1.74; unpaired t test, p = 1.70*10-6). We used Mahalanobis distance to determine if social and nonsocial reward responses were different at a population level (Figure q,r). We found that there was a significantly larger population response to social rewards compared to sucrose rewards in male and female mice (Figure 4q,r: unpaired t test, male: p = 1.71*10-8; female: p = 2.07*10-5). Sucrose and social reward trials also occupy distinct neural subspaces (Figure 4s,t, left panel: total variance explained, male: 63.8%; female: 62.8%) in both sexes as measured by Euclidean distance between and within trial type population vectors (Figure 4s,t, right panel: male - average distance within: 45.63 ± 4.21, between: 55.14 ± 5.19; paired t test, p = 5.81*10-4; female - average distance within: 47.16 ± 3.22, between: 67.17 ± 5.80; paired t test, p = 0.0020). Interestingly, across sexes, PC1 weights are higher for neurons classified as either social or sucrose reward-responsive compared to those that are reward unresponsive, further supporting the encoding of distinct social and nonsocial reward information at the population level in the mPFC (Figure 4s,t, middle panel: one-way ANOVA with post hoc t tests, male: p = 9.22*10-17, female: p = 2.93*10-8). Non-overlapping social and nonsocial reward representations in the mPFC were also seen in mice that were passively exposed to either a social target or sucrose solution (Supplementary Figure 10m-o) and when the nonsocial reward stimulus was water instead of sucrose (Supplementary Figure 10a-l).

These data suggest that in male mice, reward-responsive neurons fall into three broad categories: one that is exclusively excited by social reward (Figure 4f,i,l: left column), a second that is excited by both social and sucrose reward (Figure 4g,j,m: left column) and a third that is excited by social reward and inhibited by sucrose reward (Figure 4h,k,n: left column). However, in female mice, there is increased selectivity amongst the categories of responses such that the non-specific reward excite category seen in male mice is exclusively excited by sucrose reward in female mice (Figure 4g,j,m: right column). The sex-dependent differences in social and nonsocial reward representations in the mPFC are also observed at the population level.

### Optogenetic manipulation of mPFC neurons disrupts reward-seeking behavior

Since we found that the largest proportion of mPFC neurons were reward-responsive during the two choice operant assay, we then wanted to determine if optogenetic activation of mPFC activity during the reward period could alter reward-seeking behavior. We injected male and female mice with channelrhodopsin (ChR2), implanted ferrules bilaterally in the mPFC (Figure 5b,c) and trained those mice on the two choice assay (Figure 5a). Once mice were trained, we used blue light to activate mPFC neurons during the reward period on a random 50% of trials over the course of 7 sessions (Supplementary Figure 11a). We found that activation of mPFC neurons during the reward period increased both reward latency and number of reward fails on both trial types when compared to controls across sexes (Figure 5d,e,h,i: two-factor ANOVA with virus (ChR2/GFP) and trial type (sucrose/social) as factors, male: reward latency - interaction: p = 0.21, virus: p = 7.52*10-9, trial type: p = 5.62*10-13 with post-hoc unpaired t tests comparing virus within trial type, sucrose: p = 1.95*10-6, social: p = 5.51*10-5; reward fails - interaction: p = 6.57*10-6, virus: p = 2.38*10-10, trial type: p = 3.21*10-8 with post-hoc unpaired t tests comparing virus within trial type, sucrose: p = 2.83*10-8, social: p = 0.0042; female: reward latency - interaction: p = 0.064, virus: p = 4.09*10-13, trial type: p = 2.05*10-19, with post-hoc unpaired t tests comparing virus within trial type, sucrose: p = 7.92*10-10, social: p = 3.81*10-7; reward fails - interaction: p = 0.0026, virus: p = 1.14*10-9, trial type: p = 0.0002, with post-hoc unpaired t tests comparing virus within trial type, sucrose: p = 9.98*10-7, social: p = 1.63*10-4). We also found that optogenetic activation of mPFC neurons increased the average distance traveled by male and female mice in the operant arena during the reward period (Figure 5f,j: two-factor ANOVA with virus (ChR2/GFP) and trial type (sucrose/social) as factors, male - interaction: p = 0.19, virus: p = 1.41*10-18, trial type: p = 5.50*10-98, with post-hoc unpaired t tests comparing virus within trial types, sucrose: p = 3.67*10-15, social: p = 3.64*10-8; female - interaction: p = 0.97, virus: p = 2.00*10-11, trial type: p = 9.23*10-114, with post-hoc unpaired t tests comparing virus within trial types, sucrose: p = 2.71*10-6, social: p = 2.75*10-6). This effect on distance traveled was not seen on trials without laser stimulation in male mice or on sucrose trials without laser stimulation in female mice (Supplementary Figure 11d,i: two-factor ANOVA with virus (ChR2/GFP) and trial type (sucrose/social) as factors, male - interaction: p = 0.15, virus: p = 0.037, trial type: p = 4.71*10-89; female - interaction: p = 0.084, virus: p = 0.0052, trial type: p = 5.49*10-120, with post-hoc unpaired t tests comparing virus within trial types, sucrose: p = 0.86, social: p = 0.0076). Importantly, activating mPFC neurons outside of the reward period had no effect on the average distance traveled by mice in the operant arena (Supplementary Figure 11f,k: two-factor ANOVA with virus (ChR2/GFP) and stimulation (on/off) as factors, male - interaction: p = 0.55, virus: p = 0.28, stimulation: p = 0.74; female - interaction: p = 0.36, virus: p = 7.46*10-4, stimulation: p = 0.10), which suggests that the effects on distance traveled observed during the reward period are not solely the result of increased motor behavior due to mPFC activation. Additionally, we found that mPFC activation caused ChR2-expressing mice to spend less time in the social zone during stimulated social trials compared to controls (Figure 5g,k: unpaired t test, male: p = 0.015; female: p = 2.69*10-6). This effect was also seen in female, but not male mice, in trials without laser stimulation (Supplementary Figure 11 e,j: unpaired t test, male: p = 0.29; female: p = 8.02*10-4). These findings are consistent with decreased social investigation observed with optogenetic activation of mPFC neurons in previous studies^25,26^.

**Figure 5.**
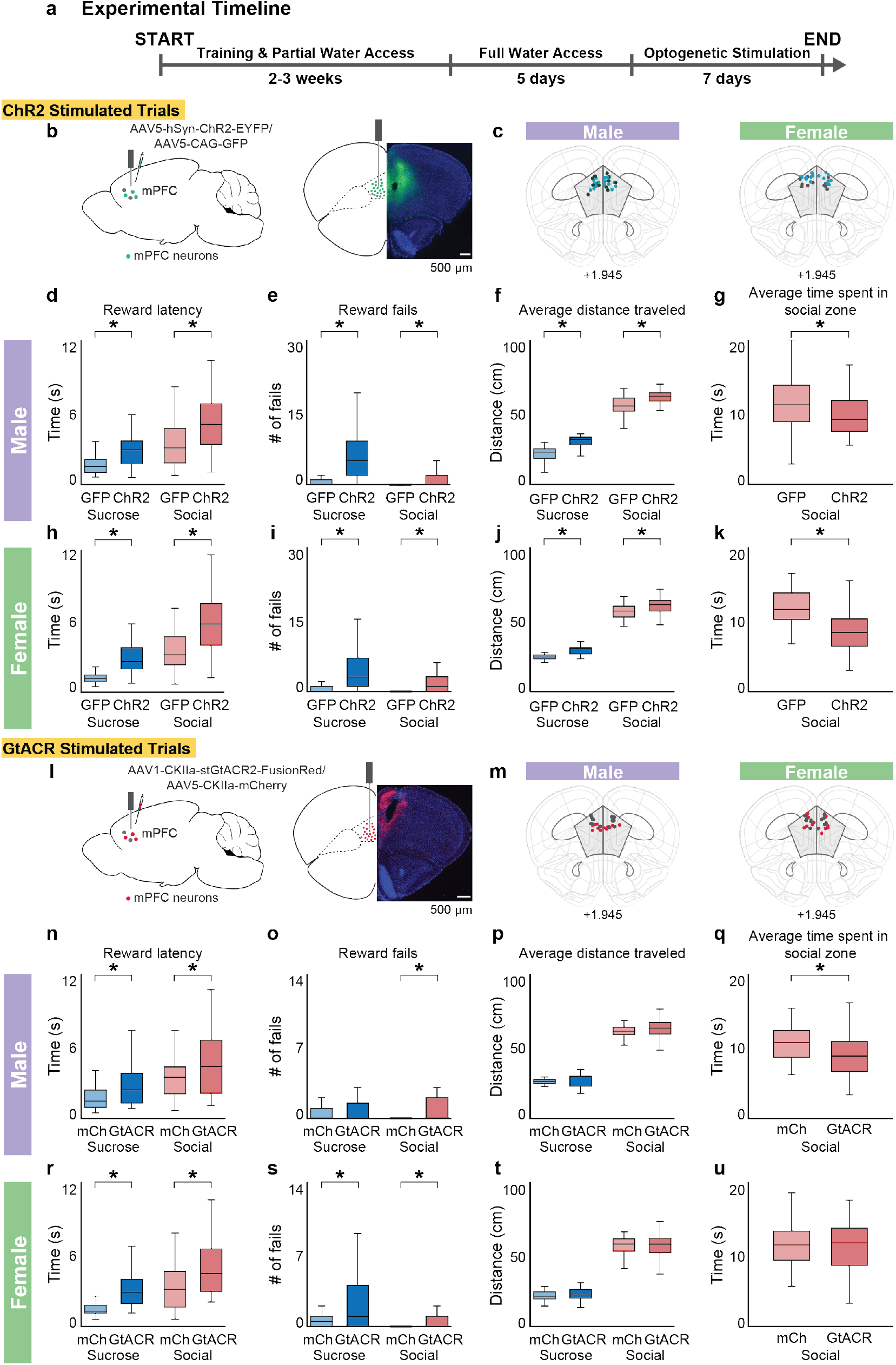
Activation and inhibition of mPFC neurons during the reward period disrupts reward-seeking behavior. **a**) Experimental timeline showing training and optogenetic stimulation schedule. b) Schematic of the viral strategy (left panel) used to label mPFC neurons with either channelrhodopsin (ChR2) or GFP (control). Example histology showing ChR2 expression (ChR2 in green, DAPI labeling of cell nuclei in blue) and ferrule placement in the mPFC at 4x magnification (right panel). Scale bar: 500 µm. c) Reconstruction of optic ferrule placement in male (left, n = 14 mice) and female (right, n = 14 mice) mice using WholeBrain. Each colored dot shows the position of the optic ferrule in the Allen Mouse Brain Common Coordinate Framework. Blue dots indicate ChR2 mice (male: n = 8 mice; female: n = 8 mice), black dots indicate GFP mice (male: n = 6 mice; female: n = 6 mice). Coronal slice is 1.945 mm anterior to bregma. d-f,h-j) Optogenetic activation of mPFC neurons during the reward period increases reward latency (**d**,**h**), reward fails (**e**,**i**) and average distance traveled (**f**,**j**) on sucrose and social trials compared to GFP controls in both male (**d**,**e**,**f**) and female (**h**,**i**,**j**) mice. Two-factor ANOVA with virus (ChR2/GFP) and trial type (sucrose/social) as factors (**d**, interaction: p = 0.21, virus: p = 7.52*10-9, trial type: p = 5.62*10-13; **e**, interaction: p = 6.57*10-6, virus: p = 2.38*10-10, trial type: p = 3.21*10-8; **f**, interaction: p = 0.12, virus: p = 1.41*10-18, trial type: p = 5.50*10-98; **h**, interaction: p = 0.064, virus: p = 4.09*10-13, trial type: p = 2.05*10-19; **i**, interaction: p = 0.0026, virus: p = 1.14*10-9, trial type: p = 0.0002; **j**, interaction: p = 0.97, virus: p = 2.00*10-11, trial type: p = 9.23*10-114) with post-hoc unpaired t tests comparing virus within trial type (**d**, sucrose: p = 1.95*10-6, social: p = 5.51*10-5; **e**, sucrose: p = 2.83*10-8, social: p = 0.0042; **f**, sucrose: p = 3.67*10-15, social: p = 3.64*10-8; **h**, sucrose: p = 7.92*10-10, social: p = 3.81*10-7; **i**, sucrose: p = 9.98*10-7, social: p = 1.63*10-4; **j**, sucrose: p = 2.71*10-6, social: p = 2.75*10-6). g,k) Optogenetic activation of mPFC neurons also caused a decrease in the time spent in the social zone during the reward period when compared to GFP controls in male (**g**) and female (**k**) mice. Unpaired t test (male: p = 0.015; female: p = 2.69*10-6). l) Schematic of the viral strategy (left panel) used to label mPFC neurons with either GtACR inhibitory opsin (GtACR) or mCherry (control). Example histology showing GtACR expression (GtACR in red, DAPI labeling of cell nuclei in blue) and ferrule placement in the mPFC at 4x magnification (right panel). Scale bar: 500 µm. m) Reconstruction of optic ferrule placement in male (left, n = 13 mice) and female (right, n = 13 mice) mice using WholeBrain. Each colored dot shows the position of the optic ferrule in the Allen Mouse Brain Common Coordinate Framework. Red dots indicate GtACR mice (male: n = 7 mice; female: n = 7 mice), black dots indicate mCherry mice (male: n = 6 mice; female: n = 6 mice). Coronal slice is 1.945 mm anterior to bregma. n,r) Optogenetic inhibition of mPFC neurons during the reward period increases reward latency on both social and sucrose trials compared to mCherry controls in male (**n**) and female (**r**) mice. Two-factor ANOVA with virus (GtACR/mCherry) and trial type (sucrose/social) as factors (**n**, interaction: p = 0.67, virus: p = 0.0004, trial type: p<0.00001; **r**, interaction: p = 0.83, virus: p = 2.46*10-8, trial type: p = 2.29*10-12) with post-hoc unpaired t tests comparing virus within trial type (**n**, sucrose: p = 0.0014, social: p = 0.021; **r**, sucrose: p = 5.46*10-10, social: p = 0.0014). o,s) Optogenetic inhibition also increases the number of social reward fails compared to mCherry controls in male mice (**o**) and both sucrose and social reward fails in female mice (**s**). Two-factor ANOVA with virus (GtACR/mCherry) and trial type (sucrose/social) as factors (**o**, interaction: p = 0.50, virus: p = 0.0039, trial type: p = 0.37; **s**, interaction: p = 0.055, virus: p = 0.0002, trial type: p = 0.0001) with post-hoc unpaired t tests comparing virus within trial type (**o**, sucrose: p = 0.13, social: p = 0.0084; **s**, sucrose: p = 0.002, social: p = 0.026). p,t) Optogenetic inhibition of mPFC neurons had no effect on average distance traveled during social or sucrose trials in both sexes. Two factor ANOVA (**p**, interaction: p = 0.18, virus: p = 0.16, trial type: p = 3.01*10-106; **t**, interaction: p = 0.76, virus: p = 0.42, trial type: p = 1.97*10-89) q,u) Optogenetic inhibition of mPFC neurons caused a decrease in the time spent in the social zone during the reward period when compared to mCherry controls in male (**q**) but not female (**u**) mice. Unpaired t test (male: p = 0.013; female: p = 0.31). *p<0.05. Box plots: center line denotes median, box edges indicate the 25th and 75th percentiles and whiskers extend to ± 2.7σ.

We next sought to determine if optogenetic inhibition of mPFC activity during the reward period of the two choice operant assay could alter reward-seeking behavior. We injected male and female mice with an inhibitory opsin (GtACR), implanted ferrules bilaterally in the mPFC (Figure 5l,m) and trained those mice on the two choice assay (Figure 5a). Once mice were trained, we used blue light to inhibit mPFC neurons during the reward period on a random 50% of trials over the course of 7 sessions (Supplementary Figure 11a). We found that inhibition of mPFC neurons during the reward period increased reward latency on both trial types when compared to controls across sexes (Figure 5n,r: two-factor ANOVA with virus (GtACR/mCherry) and trial type (sucrose/social) as factors, male - interaction: p = 0.67, virus: p = 0.0004, trial type: p<0.00001, with post-hoc unpaired t tests comparing virus within trial type, sucrose: p = 0.0014, social: p = 0.021; female - interaction: p = 0.83, virus: p = 2.46*10-8, trial type: p = 2.29*10-12, with post-hoc unpaired t tests comparing virus within trial type, sucrose: p = 5.46*10-10, social: p = 0.0014). Additionally, optogenetic inhibition caused an increase in the number of sucrose reward fails in male mice and in both sucrose and social reward fails in female mice compared to controls (Figure 5o,s: two-factor ANOVA with virus (GtACR/mCherry) and trial type (sucrose/social) as factors, male - interaction: p = 0.50, virus: p = 0.0039, trial type: p = 0.37, with post-hoc unpaired t tests comparing virus within trial type, sucrose: p = 0.13, social: p = 0.0084; female - interaction: p = 0.055, virus: p = 0.0002, trial type: p = 0.0001, with post-hoc unpaired t tests comparing virus within trial type, sucrose: p = 0.002, social: p = 0.026). Unlike optogenetic activation, we found that despite disrupting reward-seeking behavior, optogenetic inhibition of mPFC neurons did not affect the average distance traveled by male or female mice in the operant arena during or outside of the reward period across all trials with and without laser stimulation (Figure 5p,t: two-factor ANOVA with virus (GtACR/mCherry) and trial type (sucrose/social) as factors, male - interaction: p = 0.18, virus: p = 0.16, trial type: p = 3.01*10-106, female - interaction: p = 0.75, virus: p = 0.42, trial type: p = 1.97*10-89; Supplementary Figure 11n,s: two-factor ANOVA with virus (GtACR/mCherry) and trial type (sucrose/social) as factors, male - interaction: p = 0.69, virus: p = 0.57, trial type: p = 2.00*10-67, female - interaction: p = 0.67, virus: p = 0.20, trial type: p = 2.94*10-84; Supplementary Figure 11p,u: two-factor ANOVA with virus (GtACR/mCherry) and stimulation (on/off) as factors, male - interaction: p = 0.38, virus: p = 0.85, stimulation: p = 0.25; female - interaction: p = 0.22, virus: p = 0.45, stimulation: p = 0.22). In male mice, optogenetic inhibition of mPFC neurons during the reward period resulted in decreased time spent in the social reward zone (Figure 5q: unpaired t test, p = 0.013), an effect that was also seen in trials without laser stimulation (Supplementary Figure 11o: unpaired t test, p = 0.020). This effect was not observed in female mice on trials with or without laser stimulation (Figure 5u: unpaired t test, p = 0.31; Supplementary Figure 11t: unpaired t test, p = 0.12). Overall, these data support a causal role for the mPFC in modulating social and nonsocial reward-seeking behavior and demonstrate that intact mPFC activity during the reward period is crucial for animals to associate choice with reward during the two choice operant assay.

Optogenetic manipulation of mPFC neurons seemed to have an effect on reward-seeking behavior outside of trials in which there was laser stimulation, which resulted in increased reward latency and number of reward fails compared to controls on non-stimulated social trials in all male mice (Supplementary Figure 11b,c,l,m: two-factor ANOVA with virus (ChR2 or GtACR/GFP or mCherry) and trial type (sucrose/social) as factors, ChR2: reward latency - interaction: p = 0.016, virus: p = 8.00*10-5, trial type: p = 4.62*10-16 with post-hoc unpaired t tests comparing virus within trial type, sucrose: p = 0.074, social: p = 5.31*10-4; reward fails - interaction: p = 0.96, virus: p = 6.82*10-5, trial type: p = 0.0015 with post-hoc unpaired t tests comparing virus within trial type, sucrose: p = 0.025, social: p = 2.50*10-5; GtACR: reward latency - interaction: p = 0.0004, virus: p = 0.0001, trial type: p<0.00001, with post-hoc unpaired t tests comparing virus within trial type, sucrose: p = 0.60, social: p = 2.14*10-5; reward fails - interaction: p = 0.22, virus: p<0.00001, trial type: p = 0.38, with post-hoc unpaired t tests comparing virus within trial type, sucrose: p = 0.027, social: p = 3.04*10-5). Female mice across optogenetic manipulations showed increased reward latency and reward fails on both sucrose and social trials without laser stimulation compared to controls. (Supplementary Figure 11g,h,q,r: two-factor ANOVA with virus (ChR2 or GtACR/GFP or mCherry) and trial type (sucrose/social) as factors, ChR2: reward latency - interaction: p = 0.11, virus: p = 2.61*10-9, trial type: p = 5.12*10-23, with post-hoc unpaired t tests comparing virus within trial type, sucrose: p = 1.61*10-8, social: p = 6.04*10-5; reward fails - interaction: p = 0.024, virus: p = 2.23*10-6, trial type: p = 0.83, with post-hoc unpaired t tests comparing virus within trial type, sucrose: p = 0.044, social: p = 1.07*10-5; GtACR: reward latency - interaction: p = 1.78*10-12, virus: p = 2.51*10-4, trial type: p = 2.24*10-14, with post-hoc unpaired t tests comparing virus within trial type, sucrose: p = 1.94*10-4, social: p = 4.25*10-9; reward fails - interaction: p = 0.74, virus: p<0.00001, trial type: p = 0.31, with post-hoc unpaired t tests comparing virus within trial type, sucrose: p = 0.0076, social: p = 3.12*10-5). These data suggest that transiently altering mPFC activity has persistent effects on social and nonsocial reward-seeking behavior during the two choice operant assay.

### Reward-seeking behavior varies with the internal state of mice

We next asked if motivation to seek social versus nonsocial reward could be modulated by changing the internal state of the animal. We changed the level of thirst that animals experienced by restricting their access to water while also monitoring the activity of mPFC neurons. We first determined if thirsty mice (on restricted water access, RW) would change their behavior on the two choice operant assay (Figure 6a). We found that all mice completed significantly more sucrose trials than social trials when on restricted water access, compared to full water access (Figure 6b,f: two-factor ANOVA with water condition (FW/RW) and trial type (sucrose/social) as factors, male - interaction: p = 1.99*10-55, water condition: p = 5.14*10-32, trial type: p = 1.22*10-56 with post-hoc unpaired t tests comparing water conditions within trial types, sucrose: p = 1.16*10-38, social: p = 8.97*10-13; female - interaction: p = 1.81*10-23, water condition: p = 2.16*10-11, trial type: p = 8.73*10-15, with post-hoc unpaired t tests comparing water conditions within trial types, sucrose: p = 6.51*10-6, social: p = 6.76*10-17). They also performed the task more quickly with significantly decreased choice latency across all trials (Figure 6c,g: two-factor ANOVA with water condition and trial type as factors, male - interaction: p = 0.88, water condition: p = 7.37*10-18, trial type: p = 0.071 with post-hoc unpaired t tests comparing water conditions within trial types, sucrose: p = 1.11*10-15, social: p = 6.99*10-7; female - interaction: p = 0.29, water condition: p = 3.97*10-7, trial type: p = 0.18, with post-hoc unpaired t tests comparing water conditions within trial types, sucrose: p = 1.00*10-5, social: p = 0.0054) and decreased sucrose reward latency (Figure 6d,h: two-factor ANOVA with water condition and trial type as factors, male - interaction: p = 0.49, water condition: p = 3.48*10-9, trial type: p = 1.48*10-19 with post-hoc unpaired t tests comparing water conditions within trial types, sucrose: p = 7.61*10-13, social: p = 0.004; female - interaction: p = 0.46, water condition: p = 3.10*10-10, trial type: p = 2.89*10-6, with post-hoc unpaired t tests comparing water conditions within trial types, sucrose: p = 5.77*10-6, social: p = 0.014) when compared to full water access. Male mice also showed decreased social reward latency on restricted water access compared to full water access (Figure 6d), while female mice showed increased social reward latencies with water restriction (Figure 6h). All mice on restricted water access made significantly fewer sucrose reward fails compared to mice on full water access and made very few social reward fails overall (Figure 6e,i: two-factor ANOVA with water condition and trial type as factors, male - interaction: p = 1.24*10-6, water condition: p = 2.47*10-7, trial type: p = 5.31*10-9 with post-hoc unpaired t tests comparing water conditions within trial types, sucrose: p = 7.34*10-7, social: p = 0.36; female - interaction: p = 3.88*10-5, water condition: p = 0.0016, trial type: p = 0.0014 with post-hoc unpaired t tests comparing water conditions within trial types, sucrose: p = 3.01*10-4, social: p = 0.025). This may be the result of increased arousal with thirst^41,49^. Taken together, these findings demonstrate that thirst significantly alters the reward-seeking behavior of mice on the two choice operant assay with mice preferentially seeking sucrose over social reward.

**Figure 6.**
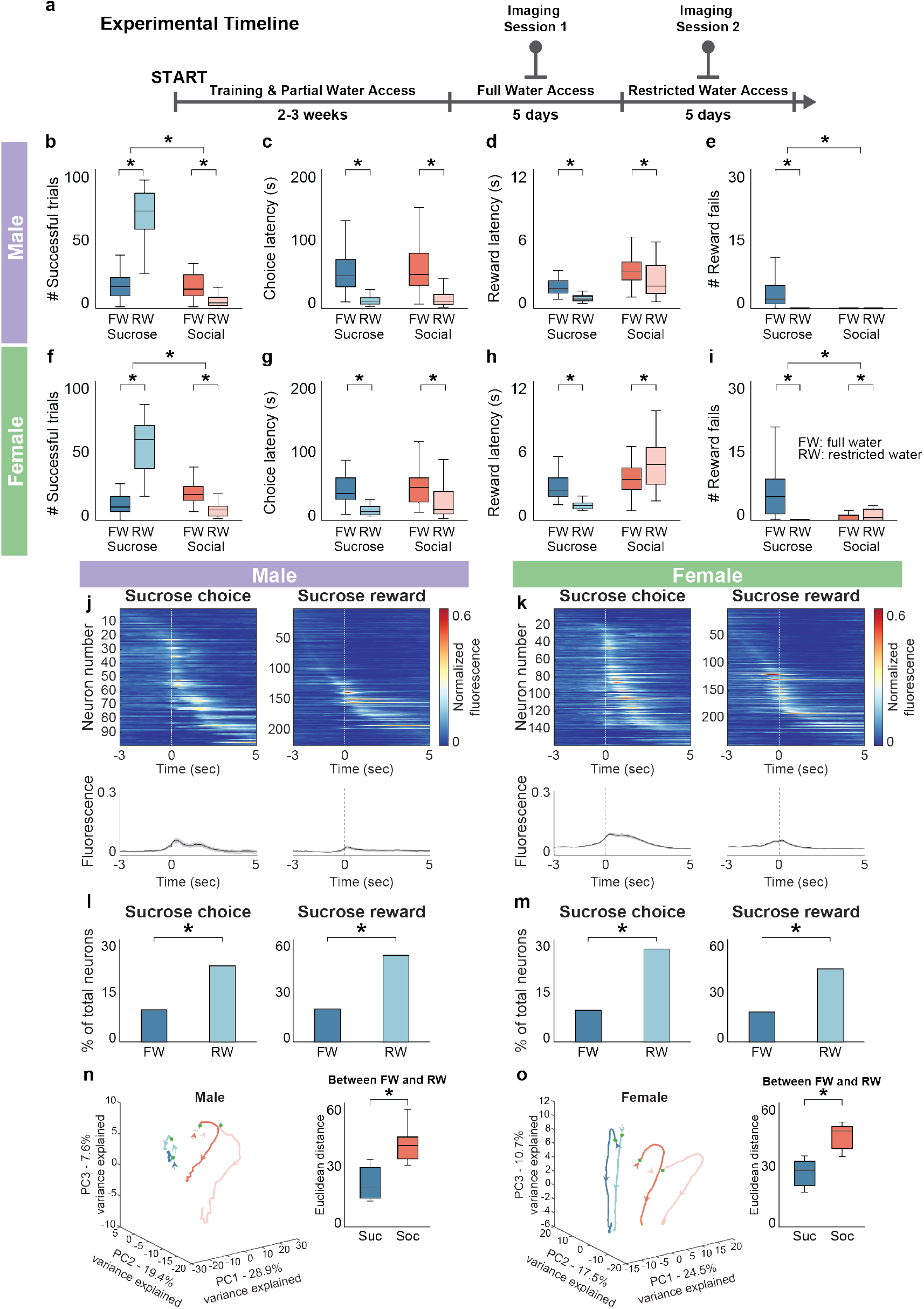
Sucrose reward-seeking behavior and mPFC neural representations of sucrose reward-seeking change with restricted water access in male and female mice. **a**) Experimental timeline showing water restriction and imaging schedule. b,f) During restricted water access conditions (RW), both male (**b**) and female (**f**) mice completed significantly more successful sucrose trials and significantly fewer successful social trials than during full water access conditions (FW). Two-factor ANOVA with water condition (FW/RW) and trial type (sucrose/social) as factors (**b**, interaction: p = 1.99*10-55, water condition: p = 5.14*10-32, trial type: p = 1.22*10-56; **f**, interaction: p = 1.81*10-23, water condition: p = 2.16*10-11, trial type: p = 8.73*10-15) with post-hoc unpaired t tests comparing water conditions within trial types (**b**, sucrose: p = 1.16*10-38, social: p = 8.97*10-13; **f**, sucrose: p = 6.76*10-17, social: p = 6.51*10-6). c,g) Male (**c**) and female (**g**) mice on RW demonstrated significantly decreased choice latency compared to FW. Two-factor ANOVA with water condition and trial type as factors (**c**, interaction: p = 0.88, water condition: p = 7.37*10-18, trial type: p = 0.071; **g**, interaction: p = 0.29, water condition: p = 3.97*10-7, trial type: p =0.18) with post-hoc unpaired t tests comparing water conditions within trial types (**c**, sucrose: p = 1.11*10-15, social: p = 6.99*10-7; **g**, sucrose: p = 1.00*10-5, social: p = 0.0054). d,h) Male (**d**) and female (**h**) mice on RW significantly decreased their sucrose reward latency compared to FW. Male mice also decreased their social reward latency on RW, while it increased in female mice. Two-factor ANOVA with water condition and trial type as factors (**d**, interaction: p = 0.49, water condition: p = 3.48*10-9, trial type: p = 1.48*10-19; **h**, interaction: p = 0.46, water condition: p = 3.10*10-10, trial type: p =2.89*10-6) with post-hoc unpaired t tests comparing water conditions within trial types (**d**, sucrose: p = 7.61*10-13, social: p = 0.004; **h**, sucrose: p = 5.77*10-6, social: p = 0.014). e,i) Male (**e**) and female (**i**) mice on RW made significantly fewer sucrose reward fails compared to FW, while female mice on RW also made significantly more social reward fails compared to FW. Two-factor ANOVA with water condition and trial type as factors (**e**, interaction: p = 1.24*10-6, water condition: p = 2.47*10-7, trial type: p = 5.31*10-9; **i**, interaction: p = 3.88*10-5, water condition: p = 0.0016, trial type: p = 0.0014) with post-hoc unpaired t tests comparing water conditions within trial types (**e**, sucrose: p = 7.34*10-7, social: p = 0.36; **i**, sucrose: p = 3.01*10-4, social: p = 0.025). j,k) Top row: Heatmaps of average normalized fluorescence of mPFC neurons in male (**j**) and female (**k**) mice that are significantly modulated by sucrose choice (left, male: n = 99 neurons; female: n = 159 neurons) and sucrose reward (right, male: n = 225 neurons; female: n = 250 neurons) during the two choice operant assay under restricted water access conditions. Rows are sorted by maximum fluorescence in each heatmap. Bottom row: Average normalized fluorescence traces of the neurons from the corresponding heatmaps in male (**j**) and female (**k**) mice that are significantly modulated by sucrose choice (left) and sucrose reward (right). l,m) During restricted water access conditions (RW), a greater proportion of mPFC neurons in both male (**l**) and female (**m**) mice are significantly modulated by sucrose choice (male: 22.30%, n = 99/444; female: 27.18%, n = 159/585) and sucrose reward (male: 50.68%, n = 225/444; female: 42.74%, n = 250/585) when compared to full water access conditions. Proportion z test with correction for multiple comparisons (male: sucrose choice: p = 9.53*10-8, sucrose reward: p<0.00001; female: sucrose choice: p = 4.22*10-15, sucrose reward: p<0.00001). n,o) Trial-averaged population neural activity traces of sucrose and social reward trials in male (**n**, left panel) and female (**o**, left panel) mice during FW (sucrose trials: dark blue, social trials: red) and RW (sucrose trials: light blue, social trials: pink) plotted on the first 3 PCs in state space. Arrowhead indicates direction of time. Filled green circle indicates reward onset. Euclidean distance separating PC-projected population vectors in a 3s window around reward is significantly greater between water restriction conditions for social than for sucrose trials in both male (**n**, right panel) and female (**o**, right panel) mice. Paired t test (male: p = 7.93*10-5; female: p = 0.0030). *p<0.05 Shaded error regions indicate ± SEM. Dashed line at zero indicates reward onset. Box plots: center line denotes median, box edges indicate the 25th and 75th percentiles and whiskers extend to ± 2.7σ.

### mPFC sucrose reward responses change with thirst through the recruitment of previously latent neurons

To determine if changes in behavior corresponded with changes in mPFC neural representations on the two choice operant assay, we performed cellular resolution calcium imaging of mPFC neurons when mice were on restricted water access (male: n = 444 neurons, 8 mice; female: n = 585 neurons, 6 mice) and compared that to full water access conditions.

We compared the neural representations of operant assay events (Figure 6j-m; Supplementary Figure 12j-l,n-p), specifically those related to sucrose reward-seeking behavior, across water access conditions. We found that across both sexes, a greater proportion of mPFC neurons were modulated by sucrose choice (Figure 6l,m: male - RW: 22.30%, 99/444, FW: 9.37%, 43/459; female - RW: 27.18%, 159/585, FW: 9.30%, 53/570; proportion z test, male: p = 9.53*10-8, female: p = 4.22*10-15) and sucrose reward (Figure 6l,m: male - RW: 50.68%, 225/444, FW: 19.17%, 88/459; female - RW: 42.74%, 250/585, FW: 17.54%, 100/570; proportion z test, male and female: p<0.00001) in restricted water access conditions compared to full water access conditions. However, the patterns of selectivity for sucrose and social reward seen in male and female mice during full water access were maintained during restricted water access (Supplementary Figure 12a-i). We also observed a decrease in the number of social reward-responsive neurons in male and female mice following water deprivation (Supplementary Figure 12l,p: male - RW: 4.28%, 19/444, FW: 29.19%, 134/459; female - RW: 25.60%, 150/585, FW: 35.44%, 202/570; proportion z test, male: p<0.00001, female: p = 2.99*10-4). At a population level, sucrose and social reward trials in both FW and RW conditions continued to occupy distinct trajectories in the first 3 PCs of state space (Figure 6n,o, left panel: total variance explained, male: 55.9%; female: 52.7%) in both sexes. However, as measured by Euclidean distance between and within trial type. Despite the changes seen in neural responsiveness, mPFC activity during the reward period decoded trial type with comparably high accuracy across water access conditions (Supplementary Figure 12m,q: average decoding accuracy, male - FW: 77.67 ± 4.55, RW: 69.84 ± 4.06; female - FW: 90.58 ± 1.26, RW: 90.11 ± 3.10; unpaired t test, male: p = 0.21; female: p = 0.91).

These findings raise the question of how the mPFC is able to maintain flexible neural representations of social and nonsocial rewards that are dependent on the internal state of the animal. One possibility is that individual mPFC neurons may change their identity to encode social or nonsocial reward information depending on the internal state of the animal. Alternatively, previously latent populations of mPFC neurons may be activated depending on the internal state of the animal. To distinguish between these possibilities, we utilized the high spatial resolution of calcium imaging to track individual mPFC neurons across water restriction conditions (Figure 7b)^50^. We compared the activity of neurons across three conditions, two conditions in which mice had access to full water and one condition of restricted water access (Figure 7a). Across male mice, we tracked 178 mPFC neurons between the first full water access imaging session and the restricted water access imaging session (Figure 7c, n = 8 mice), 158 mPFC neurons between the first full water access imaging session and the second full water access imaging session (Figure 7d, n = 7 mice) and 164 neurons between the restricted water access imaging session and the second full water access imaging session (Figure 7e, n = 7 mice). In female mice, we tracked 226 mPFC neurons between the first full water access imaging session and the restricted water access imaging session (Figure 7f, n = 5 mice), 256 mPFC neurons between the first full water access imaging session and the second full water access imaging session (Figure 7g, n = 5 mice) and 253 neurons between the restricted water access imaging session and the second full water access imaging session (Figure 7h, n = 5 mice).

**Figure 7.**
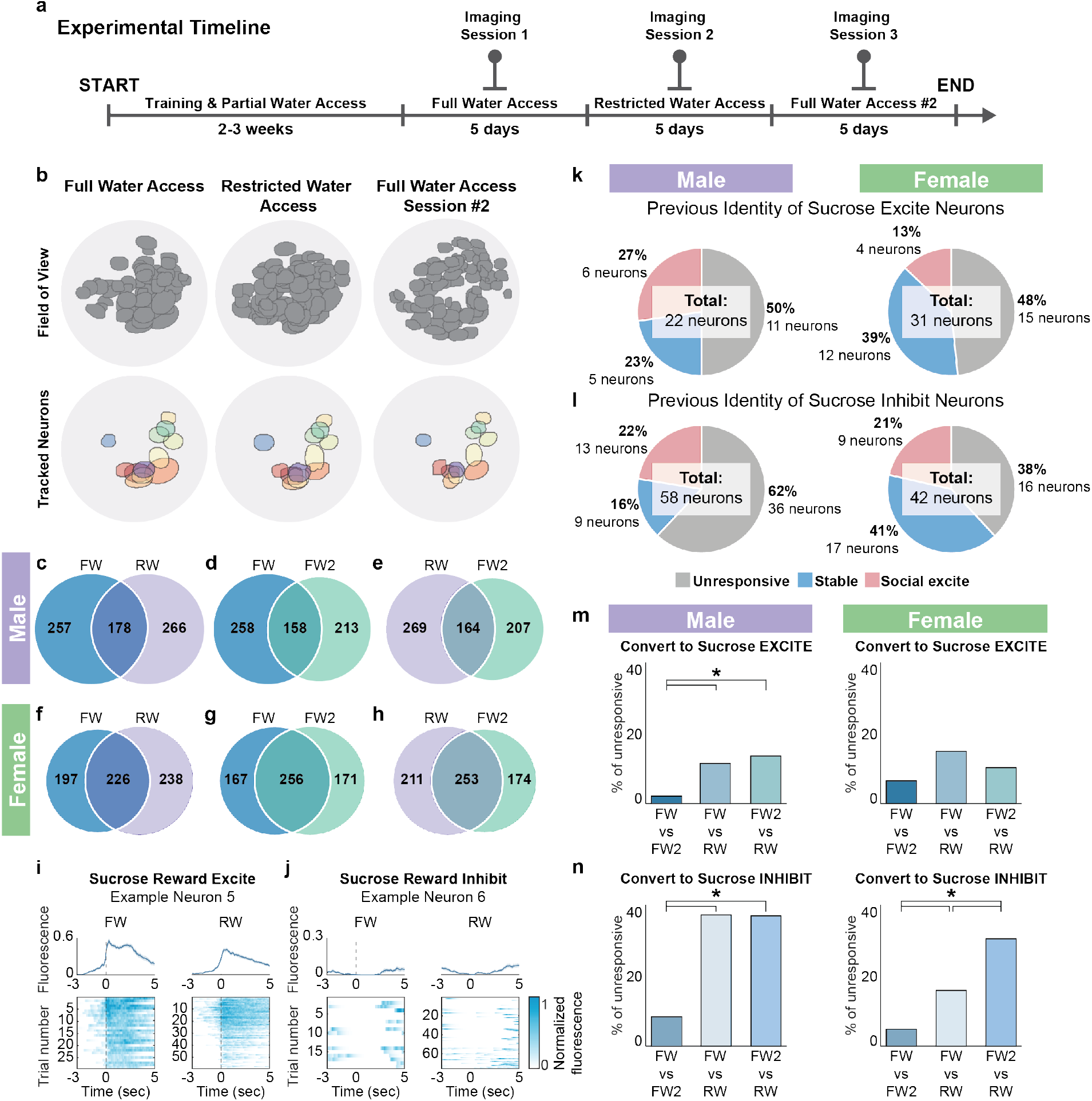
mPFC neural representations of sucrose reward change with water restriction through the recruitment of neurons that were previously reward-unresponsive. **a**) Experimental timeline showing water restriction and imaging schedule. b) mPFC neurons were tracked across three different imaging sessions using *CellReg,* a cell registration software. The top row depicts the CNMFe-generated field of view (FOV) of all identified neurons from each imaging session of a representative animal across various water access conditions. The bottom row depicts individual neurons that were successfully tracked across all three imaging sessions from the same animal. Individual neurons are represented by different colors. c-h) Venn diagrams showing the number of tracked neurons across various water restriction conditions in male (**c**-**e**) and female (**f**-**h**) mice. i,j) Time locked responses (top row: average fluorescence trace, bottom row: heatmap showing individual trials) of example neurons that showed stable sucrose reward excite (**i**) and sucrose reward inhibit (**j**) responses across both full water access (left panel) and restricted water access imaging sessions (right panel). k,l) Pie charts depicting the previous identity of sucrose reward excite (**k**) and sucrose reward inhibit neurons (**l**) in the restricted water access condition in male (left panel) and female (right panel) mice. m,n) Proportion of tracked reward-unresponsive neurons that convert to sucrose reward excite (**m**) or sucrose reward inhibit (**n**) neurons in the subsequent imaging session in male (left panel) and female (right panel) mice. Proportion z test with correction for multiple comparisons (**m**, male - FWvFW2 (1) compared to FWvRW (2): p = 0.011, FWvFW2 (1) compared to FW2vRW (3): p = 0.003, FWvRW (2) compared to FW2vRW (3): p = 0.64; female - 1 to 2: p = 0.036, 1 to 3: p = 0.27, 2 to 3: p = 0.26; **n**, male - 1 to 2: p = 2.25*10-6, 1 to 3: p = 1.65*10-6, 2 to 3: p = 0.98; female - 1 to 2: p = 0.0054, 1 to 3: p = 6.37*10-8, 2 to 3: p = 0.009). *p<0.05. Shaded error regions indicate ± SEM.

We then characterized the identity of neurons that we tracked based on their reward responses in both imaging sessions. In particular, we identified neurons as either social excite, sucrose excite, sucrose inhibit or reward-unresponsive. Neurons were considered stable if they maintained their response (excite or inhibit) to sucrose reward across imaging sessions (see example neurons Figure 7i,j). Excluding stable neurons, we found that a significantly greater proportion of mPFC neurons that were sucrose reward-responsive during the restricted water access imaging session were previously unresponsive as opposed to converting from social excite neurons during the first full water access imaging session in both male (n = 71.21%, 47/66 neurons, 8 mice) and female (n = 70.45%, 31/44 neurons, 5 mice) mice (Figure 7k,l: proportion z test, male: p = 1.09*10-6; female: p = 1.20*10-4). As a result, we next quantified the proportion of unresponsive neurons that converted to either sucrose reward excite (Figure 7m) or sucrose reward inhibit (Figure 7n) neurons across imaging sessions. We found that a significantly larger proportion of unresponsive neurons converted to either sucrose reward excite (Figure 7m: proportion z test with correction for multiple comparisons, male - FWvFW2 (1) compared to FWvRW (2): p = 0.011, FWvFW2 (1) compared to FW2vRW (3): p = 0.003, FWvRW (2) compared to FW2vRW (3): p = 0.64; female - 1 to 2: p = 0.036, 1 to 3: p = 0.27, 2 to 3: p = 0.26) or sucrose reward inhibit (Figure 7n: proportion z test with correction for multiple comparisons, male - 1 to 2: p = 2.25*10-6, 1 to 3: p = 1.65*10-6, 2 to 3: p = 0.98; female - 1 to 2: p = 0.0054, 1 to 3: p = 6.37*10-8, 2 to 3: p = 0.009) neurons across dissimilar water access conditions (full water access/full water access 2 and restricted water access) compared to similar water access conditions (full water access and full water access 2). These findings indicate that the increased proportion of sucrose reward-responsive neurons in the mPFC seen with water restriction were driven largely by the recruitment of previously reward-unresponsive neurons and less by social reward neurons changing identity or by turnover due to the passage of time.

### Neural representations of social reward change with social isolation in a sex-dependent manner

The two choice operant assay allows for the direct comparison of social and nonsocial reward-seeking behavior, which enables us to systematically change access to either type of reward outside the task and characterize how this affects motivation to seek either reward. Since water restriction preferentially drove reward-seeking behavior and mPFC neural representations toward sucrose reward, we then wanted to determine if social isolation would similarly drive preference for social reward. Consequently, we socially isolated mice for seven days after they had been trained on the two choice operant assay and imaged mPFC neurons before and after social isolation in male and female mice (Figure 8a). We found that social isolation had opposing effects on male and female reward-seeking behavior. Specifically, we found that social isolation caused an increase in the proportion of social trials completed by male mice relative to sucrose trials prior to social isolation (Figure 8b, two-factor ANOVA with isolation (pre/post) and trial type (sucrose/social) as factors; interaction: p = 0.020). In contrast, social isolation caused a relative decrease in the number of social trials completed by female mice compared to sucrose trials prior to social isolation (Figure 8i, two-factor ANOVA with isolation (pre/post) and trial type (sucrose/social) as factors; interaction: p = 0.0035). We did not observe effects of isolation on the other behavioral parameters in both sexes (Figure 8c-e, j-l).

**Figure 8.**
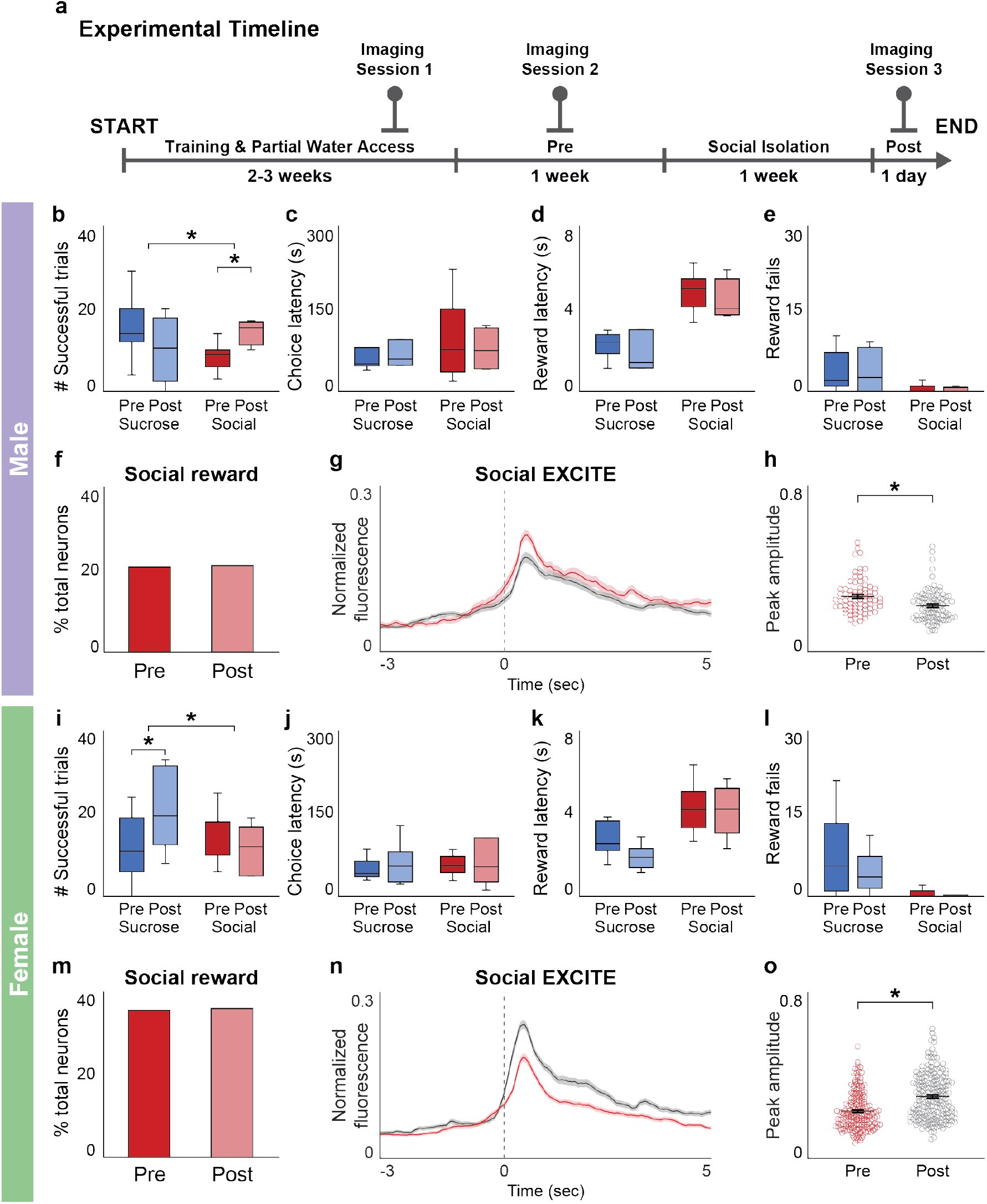
Social reward-seeking behavior and neural representations of social reward change in a sex-dependent manner following social isolation. **a**) Experimental timeline showing social isolation and imaging schedule. b) Male mice showed a relative increase in the number of successful social trials compared to sucrose trials completed after social isolation. Two-factor ANOVA with isolation (pre/post) and trial type (sucrose/social) as factors (interaction: p = 0.020, isolation: p = 0.97, trial type: p = 0.52) with post-hoc unpaired t tests comparing isolation within trial types (sucrose: p = 0.21, social: p = 0.0079). c-e) Male mice did not show any significant difference in choice latency (**c**), reward latency (**d**) and reward fails (**e**) of either trial type before and after social isolation. Two-factor ANOVA with isolation (pre/post) and trial type (sucrose/social) as factors (**c**, interaction: p = 0.69, isolation: p = 0.56, trial type: p = 0.40; **d**, interaction: p = 0.81, isolation: p = 0.21, trial type: p = 1.08*10-6; **e**, interaction: p = 0.91, isolation: p = 0.81, trial type: p = 0.0075) with post-hoc unpaired t tests comparing isolation within trial types (**d**, sucrose: p = 0.36, social: p = 0.34; **e**, sucrose: p = 0.95, social: p = 0.39). f) Male mice did not show a difference in the proportion of mPFC neurons significantly modulated by social reward before and after social isolation. Proportion z test (p = 0.91). g) Average normalized fluorescence traces of mPFC neurons significantly excited by social reward before (red) and after (gray) social isolation in male mice time-locked to social reward shown (left panel). h) Comparison of peak fluorescence before (n = 76 neurons) and after (n = 89 neurons) social isolation shows that neural responses to social reward decreased with social isolation in male mice (right panel). Unpaired t test (p = 4.24*10-4). i) Female mice showed a relative decrease in the number of successful social trials compared to sucrose trials completed after social isolation. Two-factor ANOVA with isolation and trial type as factors (interaction: p = 0.0035, isolation: p = 0.29, trial type: p = 0.30) with post-hoc unpaired t tests comparing isolation within trial types (sucrose: p = 0.017, social: p = 0.10). j-l) Female mice did not show any significant difference in choice latency (**j**), reward latency (**k**) and reward fails (**l**) of either trial type before and after social isolation. Two-factor ANOVA with isolation and trial type as factors (**j**, interaction: p = 0.61, isolation: p = 0.29, trial type: p = 0.41; **k**, interaction: p = 0.24, isolation: p = 0.13, trial type: p = 1.00*10-4; **l**, interaction: p = 0.36, isolation: p = 0.29, trial type: p = 0.0005) with post-hoc unpaired t tests comparing isolation within trial types (**k**, sucrose: p = 0.069, social: p = 0.81; **l**, sucrose: p = 0.42, social: p = 0.33). m) Female mice did not show a difference in proportion of mPFC neurons significantly modulated by social reward before and after social isolation. Proportion z test (p = 0.86). n) Average normalized fluorescence traces of mPFC neurons significantly excited by social reward before (red) and after (gray) social isolation in female mice time-locked to social reward (left panel). o) Comparison of peak fluorescence before (n = 177 neurons) and after (n = 193 neurons) social isolation shows that neural responses to social reward increased with social isolation in female mice (right panel). Unpaired t test (p = 7.58*10-11). *p<0.05. Shaded error regions indicate ± SEM. Dashed line at zero indicates reward onset. Box plots: center line denotes median, box edges indicate the 25th and 75th percentiles and whiskers extend to ± 2.7σ.

We next determined the effects of social isolation on mPFC neuronal activity. Across all mice, we found that the proportion of social reward neurons in the mPFC did not change with social isolation in male (pre: n = 20.59%, 84/408, post: n = 20.90%, 98/469, 4 mice) or female (pre: n = 35.44%, 202/570, post: n = 35.95%, 206/573, 6 mice) mice (Figure 8f,m: proportion z test, male: p = 0.91; female p = 0.86). However, when we compared the peak amplitude of social reward excite neuron responses, we found sex-dependent changes in peak amplitude following social isolation. Specifically, in male mice the peak amplitude of social reward excite neurons significantly decreased after social isolation (Figure 8g,h: average peak amplitude, pre: 0.27 ± 0.0094, post: 0.22 ± 0.0083; unpaired t test, p = 4.24*10-4), while in female mice the peak amplitude of social reward excite neurons significantly increased after social isolation (Figure 8n,o: average peak amplitude, pre: 0.23 ± 0.0069, post: 0.30 ± 0.0080; unpaired t test, p = 7.58*10-11). This sex-dependent change in the peak amplitude of the response of social reward excite neurons following social isolation was specific to social reward neurons. We compared the peak amplitude of sucrose reward excite (Supplementary Figure 14a,c: unpaired t test, male: p = 0.17; female: p = 0.66),and sucrose reward inhibit neurons (Supplementary Figure 14b,d: unpaired t test, male: p = 0.077; female: p = 0.30) before and after social isolation and found that there was no change in peak amplitude. To rule out the effects of change with time on neural responses, we compared the proportion of social reward neurons and the peak amplitude of their social reward responses across the partial water access (Supplementary Figure 13) and the full water access imaging sessions before social isolation. We also observed no difference in the proportion of social reward excite neurons between partial water access and full water access pre-isolation sessions (Supplementary Figure 14e,g: male - pre: 20.59%, n = 84/408, PW: 21.28%, n = 90/423; female - pre: 35.44%, n = 202/570, PW: 32.37%, n = 180/556, proportion z test, male: p = 0.81, female: p = 0.28). However, unlike with social isolation, there was no change in the amplitude of the responses of social reward excite neurons (Supplementary Figure 14f,h: unpaired t test, male: p = 0.74, female: p = 0.21). These data demonstrate that the sex-dependent changes in peak amplitude seen with social isolation are specific to social reward excite neurons as a result of social isolation.

## Discussion

Using a fully automated, novel two choice operant assay, we found that largely non-overlapping populations of neurons in the mPFC respond to social and sucrose reward and that these representations are more distinct in female mice compared to male mice. We additionally showed that manipulating mPFC activity through both optogenetic activation and inhibition during the reward period disrupts the ability of male and female mice to successfully engage with both types of reward, demonstrating that the mPFC plays a causal role in modulating social and nonsocial reward-seeking behavior. Finally, we found that changing internal state through either water deprivation or social isolation differentially modulates sucrose and social reward representations in a sexually dimorphic manner. Together, these findings provide exciting new insights into how social and nonsocial reward information is organized in the mPFC.

### Advantages offered by the two choice (social-sucrose) operant assay

The use of operant assays in rodent models has significantly contributed to our understanding of the neural circuitry underlying the seeking and processing of nonsocial reward-related behaviors, such as food rewards^51–53^. While we know that mice find social interactions rewarding, the complexity of social interactions has made understanding how the brain represents social interactions challenging^1^. Most studies determine whether a conspecific is perceived as rewarding by comparing the time mice spend in the proximity of a novel mouse relative to an object or an empty cage^3,4^. Although these assays are straightforward, ascertaining social reward-seeking behavior from these free ranging interactions is challenging. These assays also rarely compare social reward to other strongly appetitive stimuli, such as sucrose. As a result, similarities and differences in how social and nonsocial reward-seeking behaviors are encoded in the brain remain poorly understood. To overcome these challenges, we developed a low-cost, fully automated, novel two choice operant paradigm in which mice can freely choose between social and sucrose rewards. This assay allows for unsupervised high throughput data collection, permitting us to collect hundreds of social and sucrose trials per day across several animals.

Additionally, the operant nature of the assay allows us to constrain social behavior and monitor social reward-seeking with high temporal precision. Using this assay, we found that in control conditions (not social isolation or water deprivation), male mice showed an equal preference for social and nonsocial rewards and female mice slightly preferred social rewards over sucrose rewards (Figure 1d,h; Supplementary Figure 3i). Our results confirmed recent findings that adult mice find social stimuli reinforcing and will work to gain access to a social stimulus in operant assays^8–10,12^. Building off this work, we were able to directly compare the responses of mPFC neurons to social and nonsocial rewards while mice demonstrated similar motivation to seek both rewarding stimuli. This approach can be used to systematically map how social and nonsocial reward representations are transformed across the reward circuitry, by imaging or recording in regions such as the basolateral amygdala, ventral tegmental area, and the nucleus accumbens while mice engage in the two choice operant assay. In the future, this assay can also be used to vary dimensions of social reward, such as proximity of the social reward, valence, and reward duration independently and parametrically in freely behaving mice. Finally, although we did not observe strain differences in social and nonsocial reward-seeking behavior in female mice on our assay (Supplementary Figure 6), others have recently found strain differences in social reward-seeking behavior, with female CD1 mice showing a stronger preference for the social reward relative to C57BL/6j (used in this study)^9^. This raises interesting questions about how neural circuit differences across strains and species might give rise to varied behavioral outcomes.

### Social and nonsocial reward representations in the medial prefrontal cortex

There is little consensus regarding how social and nonsocial reward-related behaviors are mediated within the same brain region. One theory proposes that social and nonsocial behaviors are represented by discrete and mutually inhibitory neural populations^54^. Support for this theory comes from a recent study that found that neurons in the orbitofrontal cortex that are responsive to social interactions suppress feeding behavior when activated^16^. Another study also found that different populations within the medial amygdala represented distinct and antagonistic social and asocial behaviors^55^. Consistent with the idea that distinct and antagonistic populations of neurons mediate social and nonsocial reward, we observed largely non-overlapping populations of neurons within the mPFC that encoded social and nonsocial reward-related information (Figures 3 and 4). This mPFC activity was sufficient to decode the trial type during both the choice and reward periods (Figure 3g,h and Figure 4p), as well as, the sex of the animal (Figure 3i and Figure 4o). Additionally, we found evidence that these populations may regulate social and nonsocial reward information in an antagonistic manner. In particular, we identified a large subset of mPFC neurons that were inhibited in response to sucrose reward and excited in response to social reward (Figure 4h,k,n). In contrast, there were very few mPFC neurons that were inhibited by social reward (Figure 4a). These differences could arise as a result of non-mutual inhibition between social and sucrose ensembles driven by local inhibitory interneurons in the mPFC. Alternatively, they could result from differences in external inputs to the social and nonsocial reward-responsive neurons. Future cell-type specific imaging and optogenetic experiments are essential to begin to distinguish between these possibilities.

Furthermore, we showed that mPFC activity during the reward period plays a causal role in mediating reward-seeking behavior on the two choice operant assay. We found that altering mPFC activity through either optogenetic excitation or inhibition during the reward period increased reward latency and the number of reward fails across trial type in male and female mice (activation: Figure 5d,e,h,i and inhibition: Figure 5n,o,r,s). Although the role of the mPFC in nonsocial reward-seeking has been well established, these experiments are the first to suggest a causal role for the mPFC in shaping social reward-seeking behavior and not solely disrupting ongoing social interactions^25,26^. In the future, combining activity-dependent neuron tagging methods (e.g., TRAP-Cre^56^) with optogenetic tools could allow for the selective activation and silencing of social or nonsocial reward neurons. These experiments could help provide insight into how selective activity of social and nonsocial reward neurons shape reward-seeking and decision-making behavior.

### Effects of internal state and sex on neural representations of reward

There has been increasing evidence that male and female animals differ in their reward-seeking behavior, including differences in reward-associated learning strategies^43,57^ and sensitivity to reward outcomes^22,58^. Studies in rodents have also shown that there are sex differences in motivation to seek social reward, with female rodents demonstrating more robust social reward-seeking behavior than male rodents^59,60^. These findings raise the possibility that there might be differences in how social and nonsocial rewards are represented in male and female mice. In this study, we combined a novel operant assay with cellular resolution calcium imaging to identify the neural substrates that might underlie some of these sex-dependent behavioral differences.

Although male and female mice demonstrated similar levels of motivation to seek social and sucrose rewards under control conditions during the two choice operant assay, we observed sex differences in how social and nonsocial rewards were represented in the mPFC. Specifically, female mice had more distinct representations of social and nonsocial reward in the mPFC (Figure 4d,e). As a result, we were able to decode reward type with higher accuracy from the mPFC neural activity of female mice relative to that of male mice (Figure 4p). We were also able to decode the sex of the animal from the mPFC neural activity around both choice and reward (Figure 3i and Figure 4o). Interestingly, we were able to decode subsequent choice at earlier time points in female mice than in male mice (Figure 3g). Additionally, sucrose reward-responsive neurons in female mice were significantly less responsive to social reward, whereas sucrose reward-responsive neurons in male mice were equally responsive to social reward (Figure 4m). These findings raise the intriguing possibility that sex differences in mPFC activity could drive divergent reward-seeking behaviors when animals choose between competing social and nonsocial rewards.

We also found that social isolation differentially affected reward-seeking behavior in male and female mice. We observed a relative increase in the number of social trials completed by male mice and a relative decrease in the number of social trials completed by female mice compared to sucrose trials following acute social isolation (Figure 8b,i). These differences are consistent with previous findings that social isolation increases social approach behaviors in male mice while causing social withdrawal in female mice^61^. In addition, we observed that there were sex-dependent changes in how mPFC neurons responded to social reward but not sucrose reward following social isolation. In male mice, we found that social reward-responsive mPFC neurons showed a decreased peak amplitude in response to social reward following social isolation (Figure 8g,h). In contrast, in female mice, we observed an increase in the peak amplitude of social reward-responsive neurons in response to social reward following social isolation (Figure 8n,o). These changes in amplitude were socially specific, suggesting that the neural changes resulting from acute social isolation (∼1 week) act specifically on mPFC ensembles that are sensitive to social reward. These findings are consistent with a recent neuroimaging study in humans which found that acute social and food deprivation had distinct effects on social versus nonsocial craving responses in cortical regions^62^. Thus, acute social isolation may not affect nonsocial reward representations in the mPFC. However, it remains unclear if the changes observed in the neural representations of social reward in the mPFC causally contribute to the behavioral differences observed in male and female mice following social isolation. Furthermore, the cause of these sex-dependent changes in mPFC activity following social isolation remain unknown. Hormones^63^, neuromodulators^64^ and neuropeptides^65^, all of which have been shown to change in a sex-dependent manner following social isolation, may contribute to the observed changes in neural representations of social reward.

Although we found that acute social isolation did not alter mPFC responses to sucrose reward, there is substantial evidence that chronic social isolation affects nonsocial reward-seeking behaviors in rodents^66–68^ and humans^69^. Additionally, there is evidence that chronic social isolation affects mPFC activity^61,70,71^. Consequently, seven days of social isolation may not be sufficient to produce the neural and behavioral changes seen with more chronic forms of social isolation. In fact, the social homeostasis theory of social isolation postulates that social isolation activates homeostatic mechanisms that drive prosocial or antisocial behavior depending on the duration of isolation^72^. Thus, further work is needed to systematically vary the duration of social isolation to determine the precise timing of prosocial versus antisocial behaviors and the neural mechanisms underlying these changes. The two choice operant assay, as presented in this study, is uniquely suited to determine the impacts of social isolation on different reward-related behaviors, since it allows for the quantification and direct comparison of social and nonsocial reward-seeking behaviors following social isolation.

Unlike with social isolation, we did not find sex differences in changes of reward-related behavior or mPFC reward representations with water deprivation. Instead, all mice robustly changed their behavior in favor of sucrose reward-seeking. The neural representations of reward reflected these changes in behavior, with a larger proportion of mPFC neurons responding to sucrose choice and sucrose reward following water restriction (Figure 6j-m). By longitudinally tracking neurons across imaging sessions, we found that this increase in the number of sucrose reward-responsive neurons was driven by the recruitment of previously reward-unresponsive neurons (Figure 7k-n). These findings show that the reward selectivity of individual mPFC neurons rarely changes with internal state (i.e., a “social neuron” rarely converts to a “sucrose neuron” in a thirsty animal), demonstrating how thirst shapes reward-related neural activity in the mPFC. Similarly, thirst has been shown to cause widespread changes in brain activity^41,73^ and level of thirst is known to affect task performance on goal-directed behaviors^49^. A recent study also found that ventral tegmental area dopamine neurons track satiety and cause changes in dopamine dynamics in distributed brain networks^74^. In contrast, a separate dopaminergic system in the dorsal raphe nucleus has been shown to mediate behavioral changes following social isolation^75^.

These findings suggest that distinct neuromodulatory and neuropeptide systems may influence the changes seen in social and nonsocial reward representations following water deprivation and social isolation. Future studies are needed to determine how different neuromodulators and hormones act on neurons in the mPFC to modulate social and nonsocial reward representations with varying internal states in a sex-dependent manner.

### Broader implications

The ability to perceive social interactions as rewarding is necessary for appropriate social behavior. As such, a prevailing theory for the social dysfunction seen in Autism Spectrum Disorders (ASD), known as the social motivation theory, suggests that people with ASD find social interactions less rewarding^76^. However, it remains unresolved how social reward processing is disrupted in ASD. Human studies have shown that people with ASD have impairments in processing both social and nonsocial rewards^77,78^, a finding that has also been corroborated in rodent models of ASD^79,80,81^. In particular, one study found decreased differentiation between social and nonsocial odor cue encoding in the mPFC of mice with a genetic model of ASD^28^. Another study showed that social deficits in the same mouse model of ASD could be rescued by either increasing excitability of parvalbumin (PV) interneurons or decreasing excitability of pyramidal neurons in the mPFC^82^. We also found that optogenetically manipulating mPFC activity during the reward period disrupted reward-seeking behavior on the two choice operant assay (Figure 5). Thus, further interrogation of the neural mechanisms underlying social and nonsocial reward-related behavior, specifically within the mPFC, is imperative to further our understanding of diseases such as ASD. The novel two choice operant assay described in this study offers a means by which to study two competing reward-related behaviors and to provide further insight into how the brain processes social and nonsocial reward.

In summary, we developed a novel two choice (social-sucrose) operant assay that allowed us to characterize how social and nonsocial rewards are processed in the mPFC. Using this assay, we demonstrated that social and nonsocial reward representations are modulated by social isolation and thirst in a sex-dependent manner.

---

**Supplementary Figure 1.**
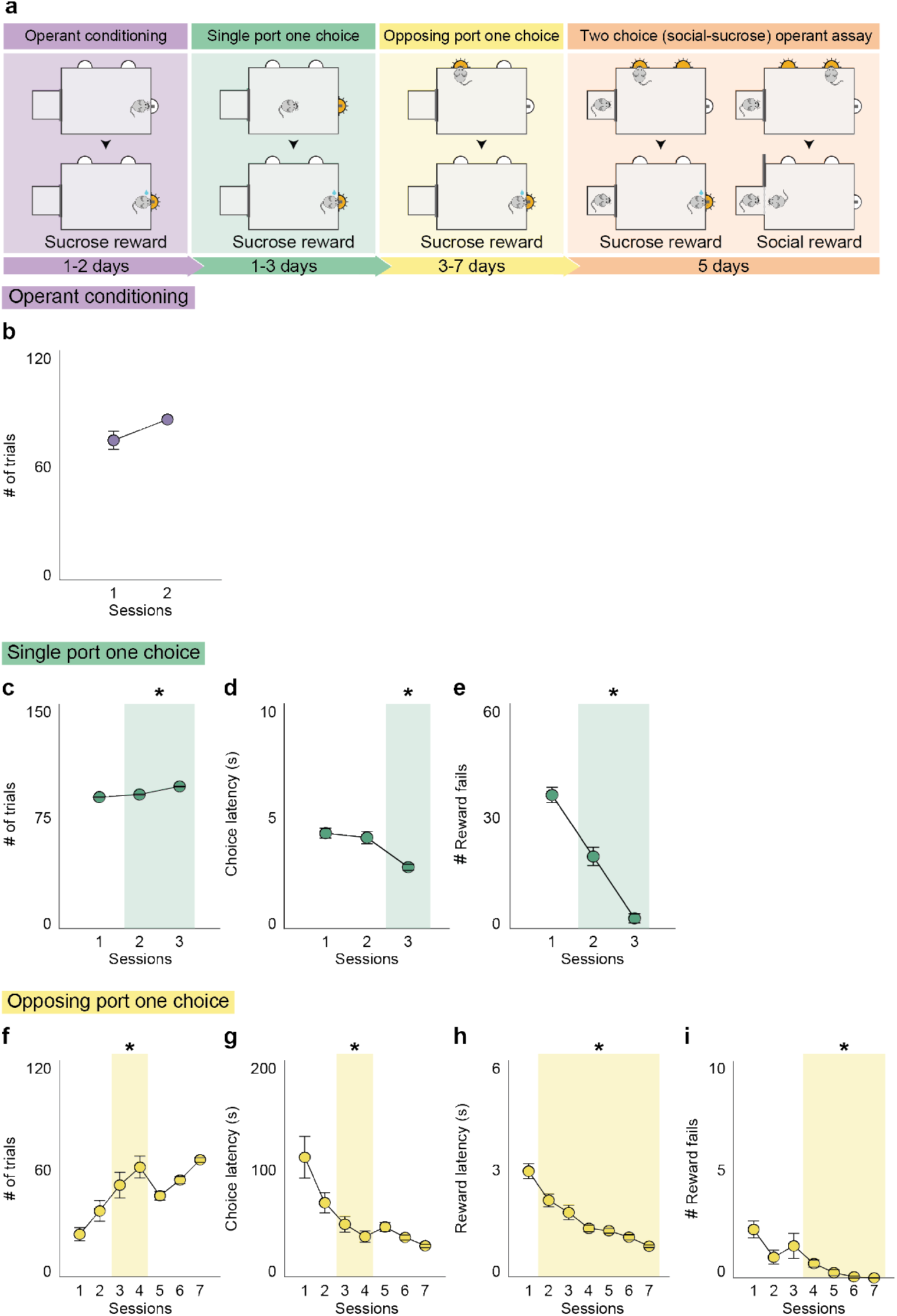
Mice are trained to engage with two choice (social-sucrose) operant assay over three training stages. **a**) Schematic of experimental training timeline. Mice were placed on restricted water access and run through a series of increasingly difficult training stages within the same operant chamber. b) All mice showed high engagement (number of pokes) with the operant conditioning paradigm within two sessions of training. c) Mice significantly increased their preference for the sucrose reward port (number of pokes) over the course of training. One-way ANOVA with post-hoc t tests (p = 6.1*10-19). d,e) Mice showed a significant decrease in sucrose reward latency (**d**) and in the number of sucrose reward fails (**e**) with training. One-way ANOVA with post-hoc t tests (reward latency: p = 0.017, reward fails: p = 1.9*10-16). f) Mice significantly increased engagement (number of pokes) with the opposing port one choice paradigm over the course of training. One-way ANOVA with post-hoc t tests (p = 6.0*10-4). g-i) In addition, they showed a significant decrease in choice latency (**g**), reward latency (**h**) and number of reward fails (**i**) with training. One-way ANOVA with post-hoc t tests (choice latency: p = 8.0*10-4, reward latency: p = 4.01*10-8, reward fails: p = 6.4*10-7). *p<0.05. N = 21 mice. Error bars indicate ± SEM.

**Supplementary Figure 2.**
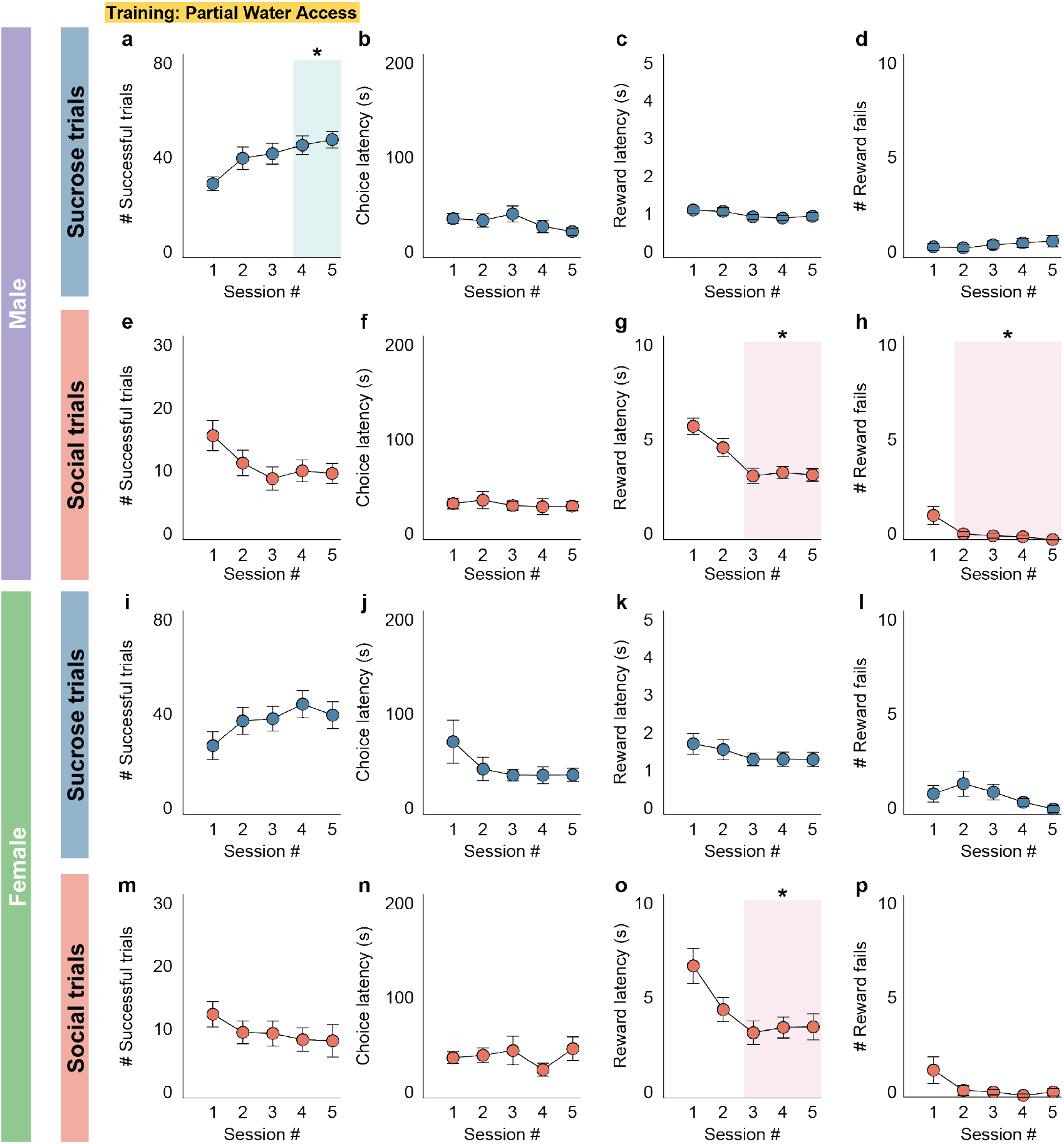
Male and female mice display positive reinforcement for social and nonsocial rewards during training on the two choice operant assay. **a-d**) Under partial water access conditions, male mice show an increased preference for the nonsocial (sucrose) reward with an increase in the number of successful sucrose trials (**a**) over five consecutive sessions on the two choice assay. Sucrose choice latency (**b**) and reward latency (**c**) remain stable across training. Mice make very few sucrose reward fails (**d**). One-way ANOVA with post-hoc t tests (number of trials: p = 0.013, choice latency: p = 0.27, reward latency: p = 0.28, reward fails: p = 0.80). e-h) Under partial water access conditions, male mice learn to associate the social choice port with social reward as demonstrated by a decrease in social reward latency (**g**) and a decrease in social reward fails (**h**) over five consecutive sessions. Social choice latency (**f**) and number of successful social trials (**e**) remain stable across training. One-way ANOVA with post-hoc t tests (number of trials: p = 0.11, choice latency: p = 0.96, reward latency: p = 8.19*10-6, reward fails: p = 0.0011). N = 21 male mice. i-l) Under partial water access conditions, female mice demonstrate stable nonsocial (sucrose) reward-seeking behavior with a consistent number of successful sucrose trials (**i**), sucrose choice latency (**j**), sucrose reward latency (**k**) and few reward fails (**l**) over five consecutive sessions on the two choice assay. One-way ANOVA (number of trials: p = 0.12, choice latency: p = 0.10, reward latency: p = 0.29, reward fails: p = 0.086). m-p) Under partial water access conditions, female mice learn to associate the social choice port with social reward as demonstrated by a decrease in social reward latency over five consecutive sessions (**o**). Number of successful social trials (**m**), social choice latency (**n**) and number of social reward fails (**p**) remain stable across training. One-way ANOVA with post-hoc t tests (number of trials: p = 0.98, choice latency: p = 0.38, reward latency: p = 0.016, reward fails: p = 0.062). N = 12 female mice *p<0.05. Error bars indicate ± SEM.

**Supplementary Figure 3.**
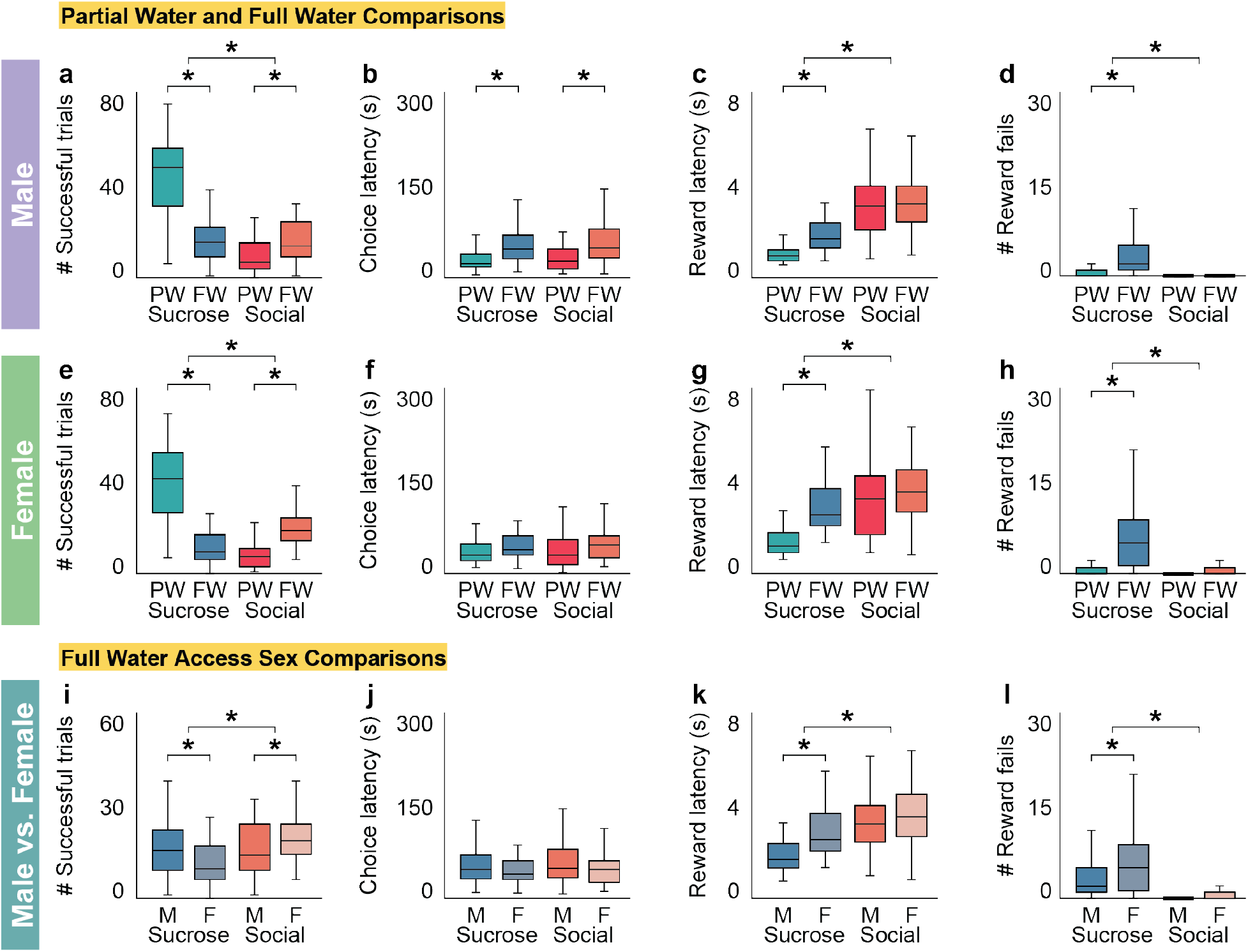
Fully trained male and female mice with full water access increase social and decrease sucrose reward-seeking behavior compared to training. **a,e**) Male (**a**) and female (**e**) mice complete significantly fewer successful sucrose trials and significantly more successful social trials during full water access conditions (FW) compared to training partial water access conditions (PW). Two-factor ANOVA with water condition (PW/FW) and trial type (sucrose/social) as factors (**a**, interaction: p = 2.63*10-26, water condition: p = 9.54*10-13, trial type: p = 1.22*10-27; **e**, interaction: p = 4.93*10-20, water condition: p = 5.76*10-6, trial type: p = 3.02*10-9) with post-hoc unpaired t tests comparing water condition within trial type (**a**, sucrose: p = 1.04*10-20, social: p = 2.32*10-5; **e**, sucrose: p = 4.49*10-13, social: p = 4.14*10-8). b,f) Male mice (**b**) complete social and sucrose trials with slower choice latency during FW compared to PW conditions, while female mice (**f**) complete social and sucrose trials with similar choice latency across both conditions. Two-factor ANOVA with water condition (PW/FW) and trial type (sucrose/social) as factors (**b**, interaction: p = 0.39, water condition: p = 6.64*10-9, trial type: p = 0.41; **f**, interaction: p = 0.91, water condition: p = 0.69, trial type: p = 0.15) with post-hoc unpaired t tests comparing water condition within trial type (**b**, sucrose: p = 6.34*10-5, social: p = 2.59*10-5). c,d,g,h) Male (**c**,**d**) and female (**g**,**h**) mice show increased sucrose reward latency and number of sucrose reward fails during FW compared to PW conditions, but maintain similar social reward latency and number of social reward fails across water access conditions. Two-factor ANOVA with water condition (PW/FW) and trial type (sucrose/social) as factors (**c,** interaction: p = 0.0036, water condition: p = 0.0003, trial type: p = 4.61*10-29; **d**, interaction: p = 1.58*10-6, water condition: p = 9.19*10-8, trial type: p = 1.42*10-11; **g**, interaction: p = 0.0025, water condition: p = 0.0002, trial type: p = 9.84*10-8; **h**, interaction: p = 1.21*10-6, water condition: p = 2.05*10-7, trial type: p = 3.23*10-8) with post-hoc unpaired t tests comparing water condition within trial type (**c**, sucrose: p = 1.31*10-12, social: p = 0.69; **d**, sucrose: p = 5.72*10-7, social: p = 0.061; **g**, sucrose: p = 9.10*10-9, social: p = 0.64; **h**, sucrose: p = 1.23*10-6, social: p = 0.12). i-l) Under full water access conditions, male mice complete a greater number of successful sucrose and a fewer number of successful social trials than female mice (**i**) at comparable choice latencies (**j**). Male mice show decreased sucrose reward latency compared to female mice (**k**). Both male and female mice make more sucrose reward fails than social reward fails and female mice make more sucrose reward fails than male mice (**l**). Two-factor ANOVA with sex (male/female) and trial type (sucrose/social) as factors (**i**, interaction: p = 0.0008, sex: p = 0.78, trial type: p = 0.0035; **j**, interaction: p = 0.57, sex: p = 0.013, trial type: p = 0.42; **k**, interaction: p = 0.032, sex: p = 0.0004, trial type: p = 1.64*10-7; **l**, interaction: p = 0.010, sex: p = 0.0048, trial type: p = 2.79*10-15) with post-hoc unpaired t tests comparing sex within trial type (**i**, sucrose: p = 0.011, social: p = 0.029; **j**, sucrose: p = 0.1154, social: p = 0.056; **k**, sucrose: p = 1.29*10-5, social: p = 0.35; **l**, sucrose: p = 0.0073, social: p = 0.28). N = 21 male mice, 12 female mice, 3 sessions per mouse. *p<0.05. Box plots: center line denotes median, box edges indicate the 25th and 75th percentiles and whiskers extend to ± 2.7σ.

**Supplementary Figure 4.**
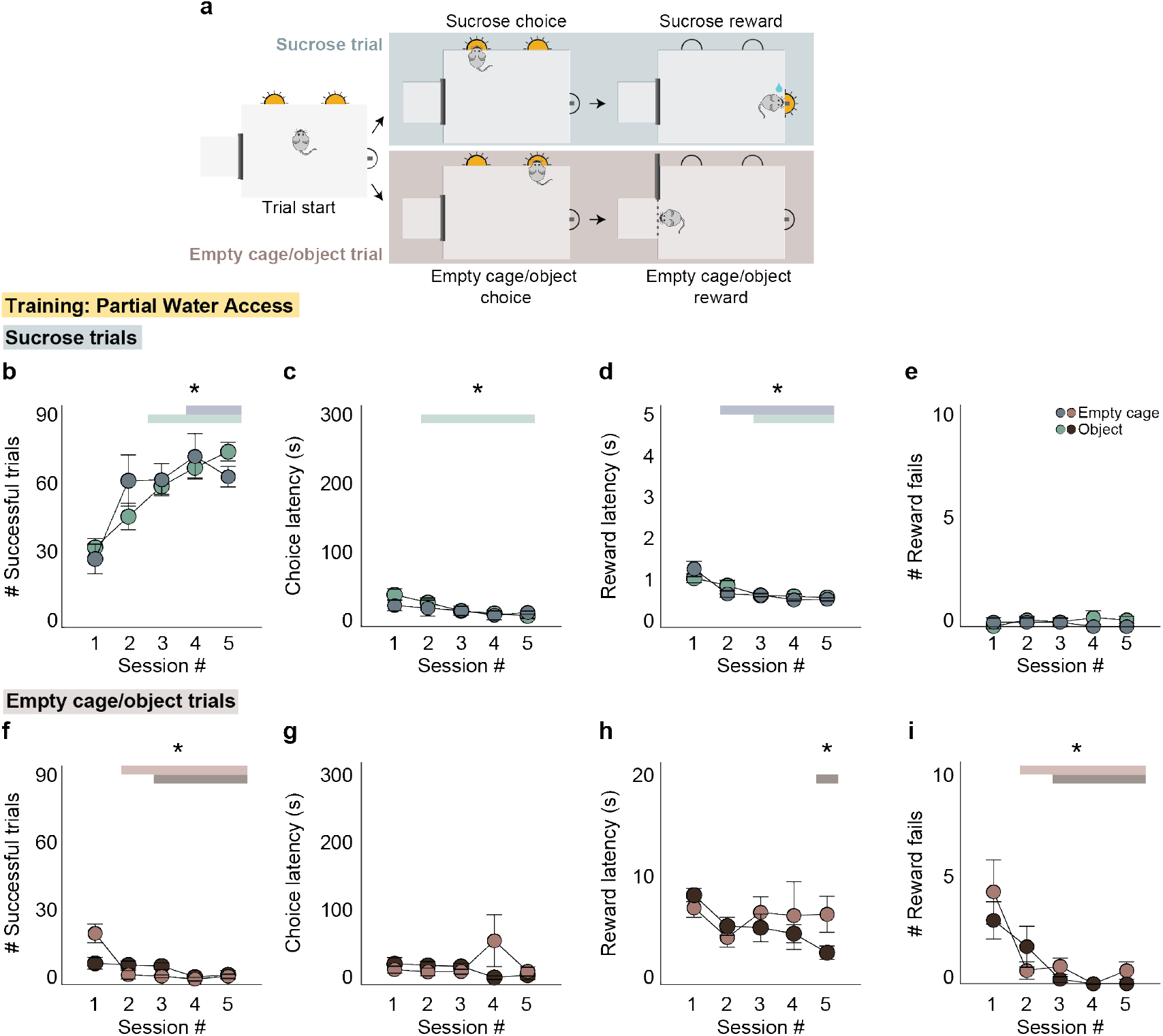
Mice do not find access to an empty cage or object positively reinforcing during training on the two choice operant assay. **a**) Schematic of the experimental setup of the two choice operant assay in which the social target has been removed and replaced with either an empty cage or a novel object. b-e) Under partial water access conditions, mice trained on the two choice operant assay with an empty social target cage or an object completed an increasing number of successful sucrose trials (**b**) and showed decreasing sucrose reward latency (**d**) over five consecutive sessions. Additionally, mice trained with an empty cage showed consistent sucrose choice latency, while mice trained with an object showed decreased sucrose choice latency (**c**). Mice in both conditions made very few reward fails (**e**) over the course of training. One-way ANOVA with post-hoc t tests (number of sucrose trials - ECG: p = 0.01, OG: p = 1.25*10-7; sucrose choice latency - ECG: p = 0.72, OG: p = 0.0015; sucrose reward latency - ECG: p = 5.03*10-5, OG: p = 0.0012; sucrose reward fails - ECG: p = 0.74, OG: p = 0.64). ECG: n = 5 mice, OG: n = 10 mice. f-i) Under partial water access conditions, mice trained on the two choice operant assay with an empty cage or an object completed fewer successful empty cage/object trials (**f**) over five consecutive sessions. Mice trained with an empty cage did not show a change in empty cage choice latency (**g**) or empty cage reward latency (**h**) with training. Mice trained with an object also did not show a change in object choice latency (**g**) and only showed a decrease in object reward latency on the final training session (**h**). Mice in both conditions made very few empty cage/object reward fails (**i**) after the first session of training. One-way ANOVA with post-hoc t tests (number of empty cage/object trials - ECG: p = 5.51*10-6, OG: p = 3.0*10-4; empty cage/object choice latency - ECG: p = 0.29, OG: p = 0.12; empty cage/object reward latency - ECG: p = 0.84, OG: p = 0.0036; empty cage/object reward fails - ECG: p = 0.0039, OG: p = 0.0015). ECG: n = 5 mice, OG: n = 10 mice. *p<0.05. Error bars indicate ± SEM.

**Supplementary Figure 5.**
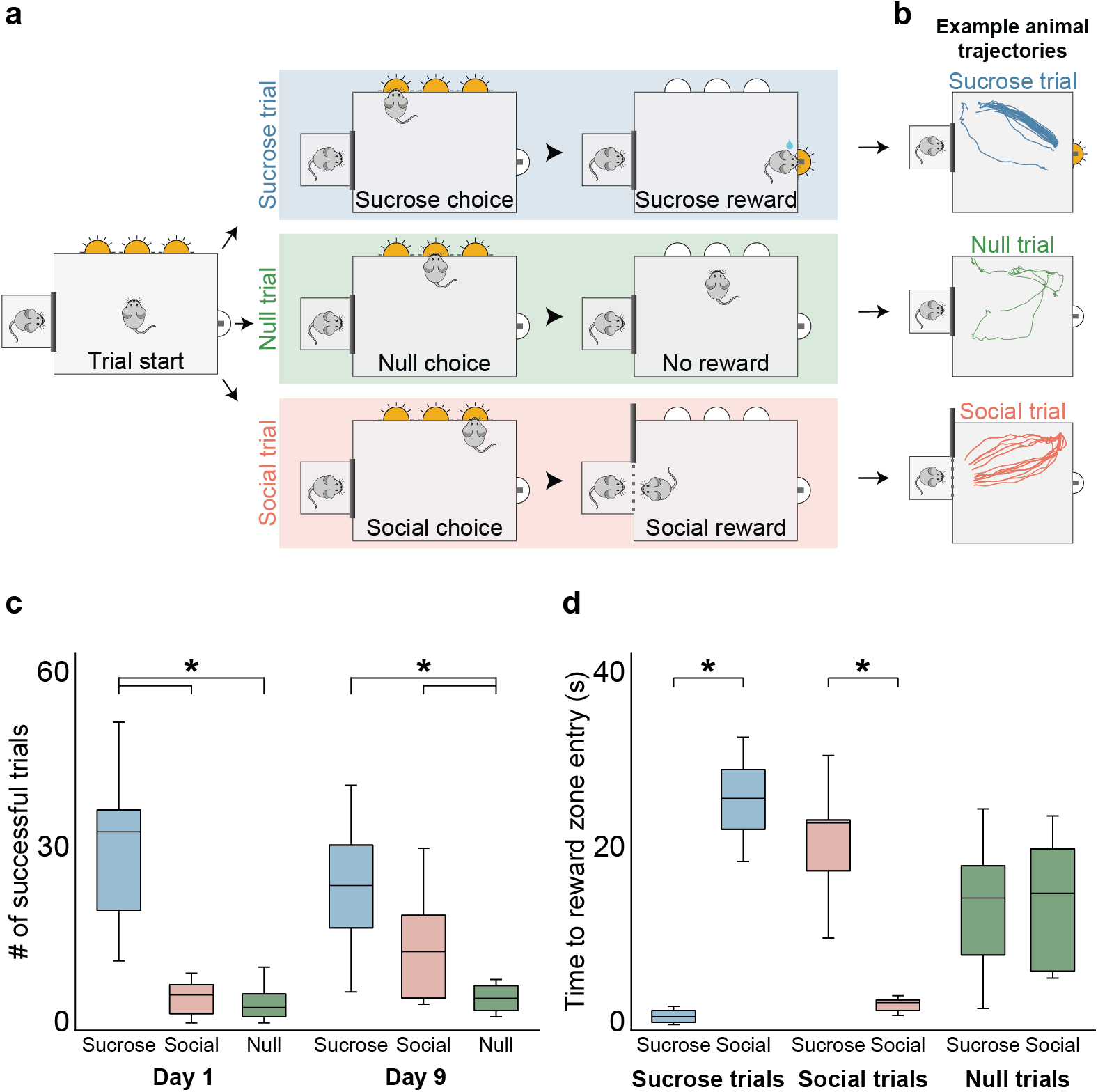
Mice preferentially increase the number of social reward trials but not null trials on a multi-choice operant assay. **a**) Schematic of the experimental setup of the multi-choice operant assay in which a third choice port (null port) has been added that results in neither a sucrose nor social reward (null trial). b) Example trajectories of mice during the reward period of the multi-choice assay on sucrose (top panel), null (middle panel) and social (bottom panel) trials (Day 9). c) On the first day of the multi-choice assay, mice complete more successful sucrose than social or null trials and an equivalent number of null and social trials. However, by the ninth day of the multi-choice assay, mice complete an equivalent number of successful sucrose and social trials and more social and sucrose trials than null trials. Two-factor ANOVA with training day (day 1/day 9) and trial type (sucrose/social/null) as factors (interaction: p = 0.0059, training day: p = 0.68, trial type: p = 5.02*10-13) with post-hoc unpaired t tests comparing trial type within training day (day 1 - suc v soc: p = 1.02*10-4, suc v null: p = 1.45*10-4, soc v null: p = 0.48; day 9 - suc v soc: p = 0.078, suc v null: p = 5.21*10-4, soc v null: p = 0.013). d) On the ninth day of the multi-choice assay, mice are significantly faster to access the reward zone associated with their choice than the opposite reward zone on sucrose or social reward trials, but not on null trials during which mice enter the sucrose and social reward zones with equivalent reward latencies. Two-factor ANOVA with choice type (sucrose/social/null) and reward zone (sucrose/social) as factors (interaction: p = 2.44*10-13, choice type: p = 0.41, reward zone: p = 0.10) with post-hoc unpaired t tests comparing reward zone within choice type (sucrose: p = 2.19*10-8, social: p = 1.82*10-5, null: p = 0.41). N = 10 mice. *p<0.05. Box plots: center line denotes median, box edges indicate the 25th and 75th percentiles and whiskers extend to ± 2.7σ.

**Supplementary Figure 6.**
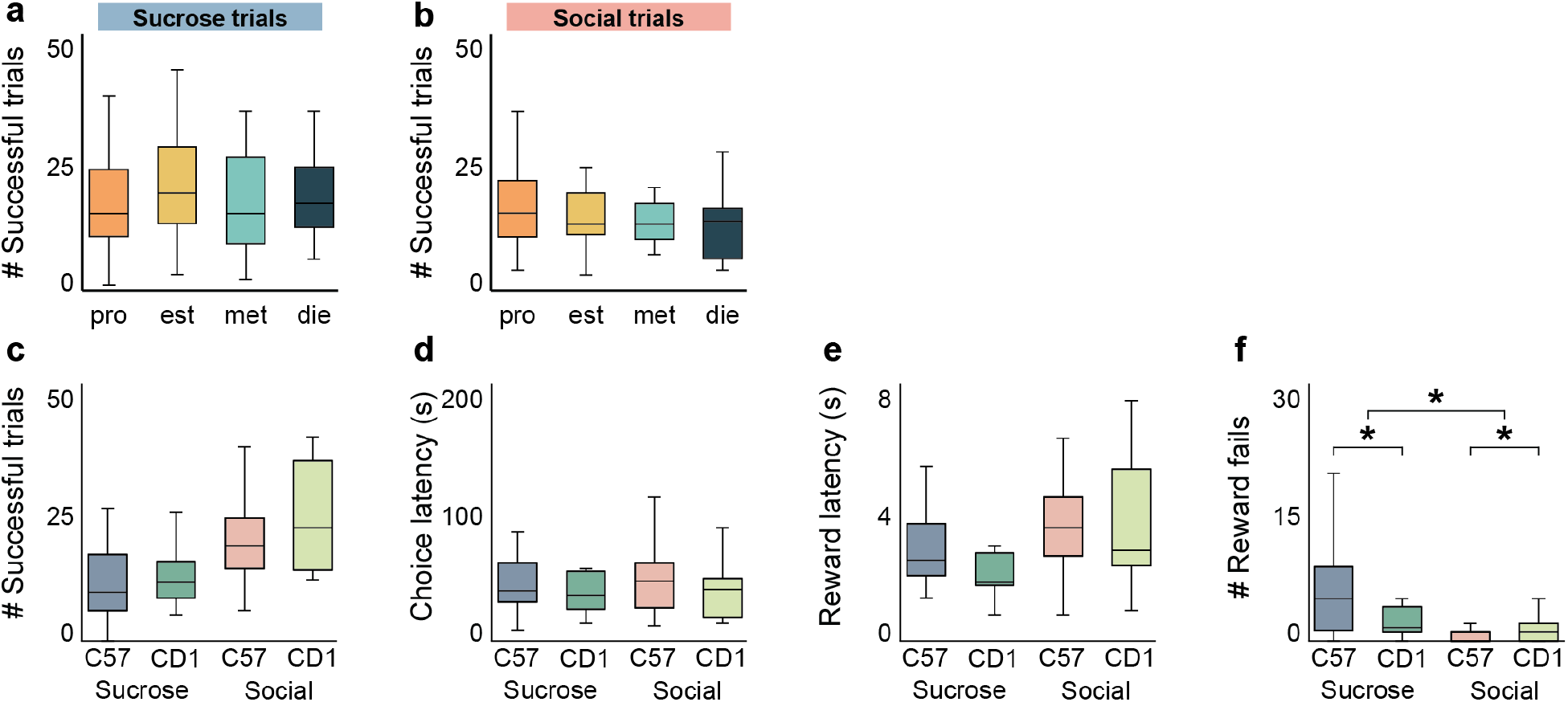
Reward-seeking behavior in female mice is not affected by estrous cycle or strain. **a,b**) The number of successful sucrose (**a**) and social (**b**) trials completed by female mice on the two choice operant assay does not differ across different stages of the estrous cycle. One-way ANOVA (**a**, p = 0.46; **b**, p = 0.52). N = 20 female mice. c-f) Female CD1 mice complete a similar number of successful sucrose and social trials (**c**) at similar choice (**d**) and reward (**e**) latencies with a similar number of reward fails (**f**) as female C57 mice on the two choice operant assay. Two-factor ANOVA with strain (C57/CD1) and trial type (sucrose/social) as factors (**c**, interaction: p = 0.31, strain: p = 0.14, trial type: p = 9.32*10-7; **d**, interaction: p = 0.38, strain: p = 0.69, trial type: p = 0.51; **e**, interaction: p = 0.23, strain: p = 0.18, trial type: p = 0.0039; **f**, interaction: p = 0.0094, strain: p = 0.086, trial type: p = 5.71*10-4) with post-hoc unpaired t tests comparing strain within trial type (sucrose: p = 3.12*10-2, social: p = 5.70*10-3). C57: n = 12 mice, CD1: n = 4 mice, 3 sessions per mouse. *p<0.05. Box plots: center line denotes median, box edges indicate the 25th and 75th percentiles and whiskers extend to ± 2.7σ.

**Supplementary Figure 7.**
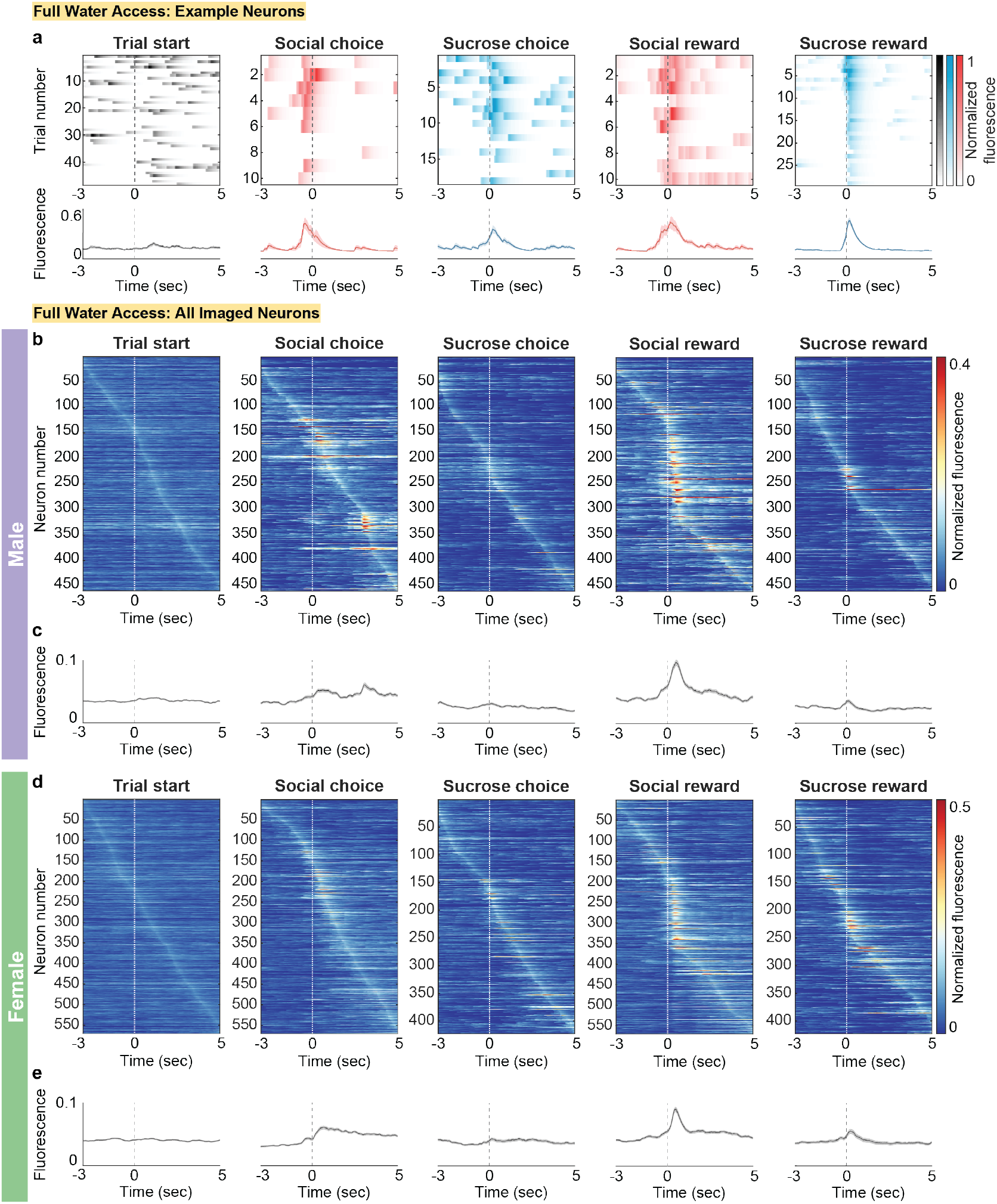
mPFC neurons in male and female mice are responsive to various task events. **a**) Responses of example mPFC neurons time-locked to each task event (from left to right: trial start, social choice, sucrose choice, social reward, and sucrose reward). The top panel shows a heatmap of the normalized fluorescence across all trials over the course of a single behavioral session and the bottom panel shows the average normalized fluorescence trace ± SEM. Dashed line at zero indicates task event onset. b,d) Heatmaps of average normalized fluorescence of all mPFC neurons recorded from male (**b**) and female (**d**) mice during the two choice operant assay aligned to each task event (from left to right: trial start, social choice, sucrose choice, social reward, and sucrose reward). Rows are sorted by maximum fluorescence in each heatmap. c,e) Average normalized fluorescence traces ± SEM of all recorded mPFC neurons from male (**c**) and female (**e**) mice aligned to each task event. Dashed line at zero indicates task event onset.

**Supplementary Figure 8.**
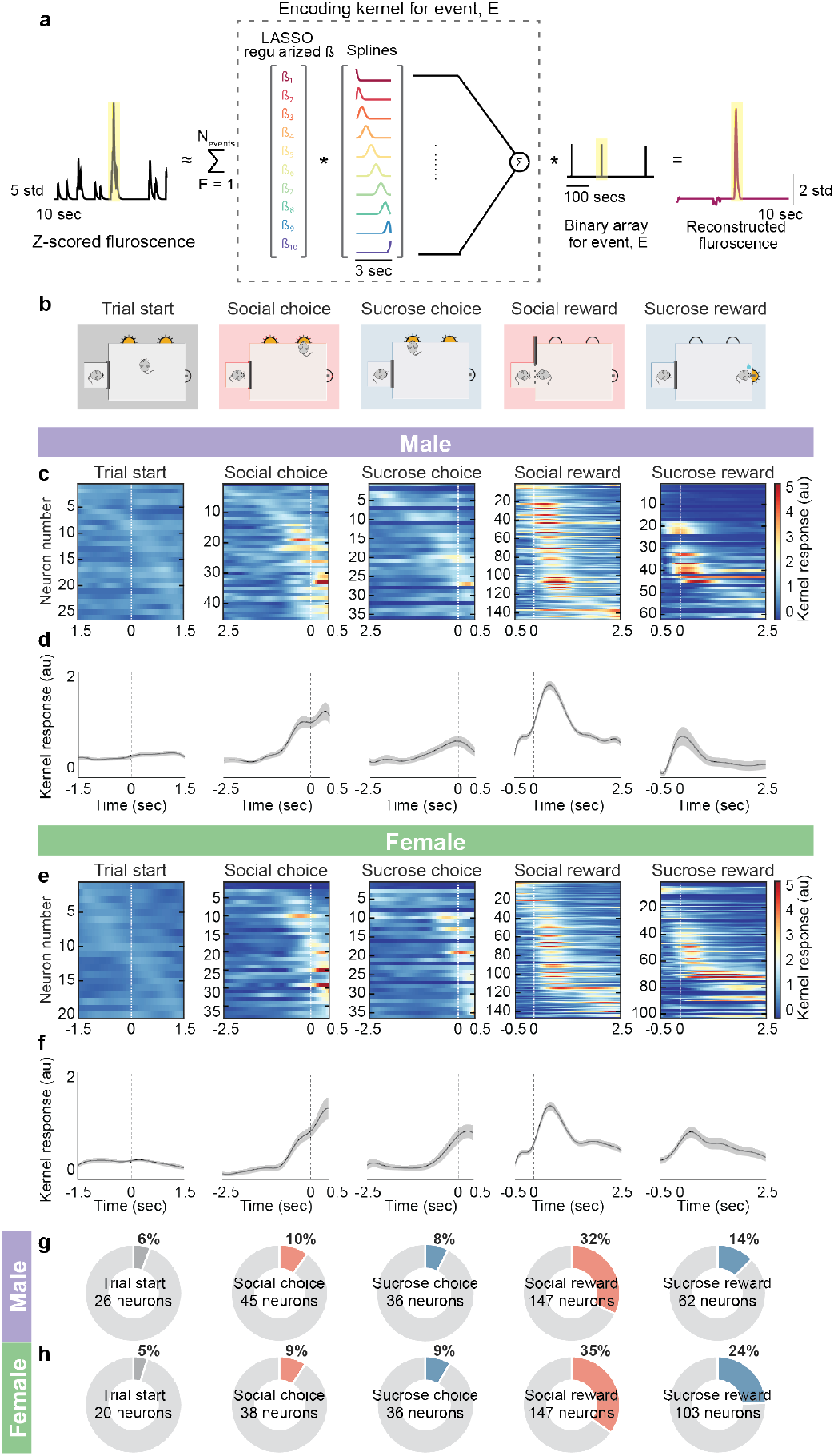
Encoding model identifies similar proportions of significantly task-modulated mPFC neurons. **a**) Schematic of the encoding model used to create a response kernel for every neuron and each operant task event. Fluorescence trace of a neuron (left panel) was estimated as the sum of all events’ response kernels (dotted box, middle panel), convolved with each event’s binary array (middle panel, right). These encoding kernels were calculated as the product between the lasso regularized beta coefficients (dotted box, middle panel, left) and a 10-spline 3 second duration basis set (dotted box, middle panel, right). Reconstructed fluorescence for one example neuron and one trial of the event is shown in the right panel. b) Schematic of the two choice operant task events. c,e) Heatmaps of average kernel responses (a.u.) of all mPFC neurons from male (**c**) and female (**e**) mice significantly modulated by each task event over a 3s window. Each row is a single neuron. Neurons are sorted by the time of maximum response across each task event. d,f) Average response traces of all neurons from the corresponding heatmap significantly modulated by each task event in male (**d**) and female mice (**f**) during a 3s window. Dashed line at zero indicates task event onset. Shaded error regions indicate ± SEM. g,h) Proportions of mPFC neurons that are significantly modulated by each task event in male (**g**) and female (**h**) mice as determined using an encoding model. Proportions are determined using the total number of recorded neurons (male: n = 459 neurons, 9 mice; female: n = 423 neurons, 5 mice).

**Supplementary Figure 9.**
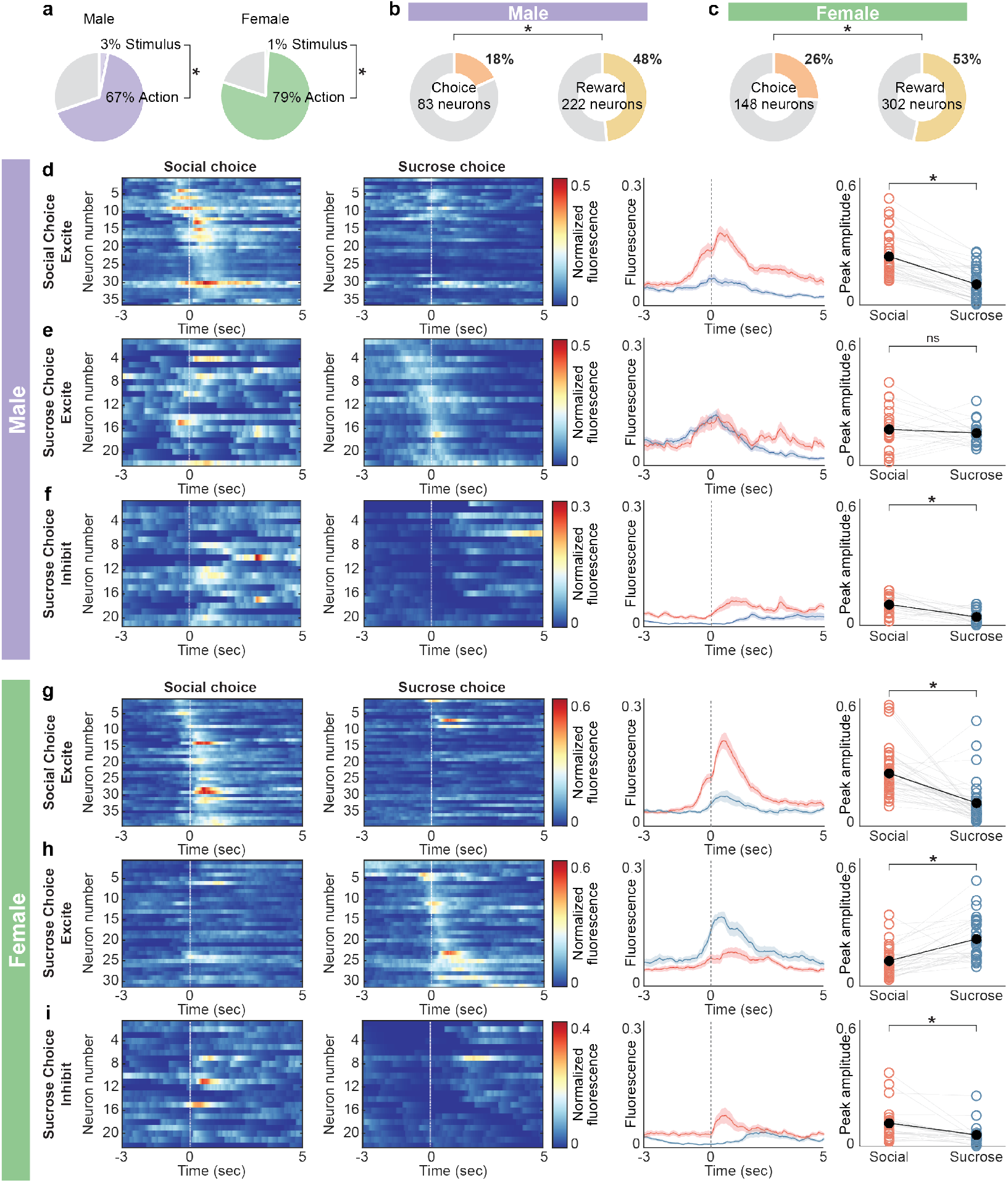
mPFC neurons represent social and nonsocial choice differently in male and female mice. **a**) Pie charts showing the distribution of responses in male (left) and female (right) to stimulus (male: n = 3.27%,15/459; female: n = 1.05%, 6/570) and action events (male: n = 66.5%, 305/459; female: n = 79.0%, 450/570) on the two choice operant assay. b,c) Proportions of total recorded neurons that are modulated by choice (left, male: n = 18.1%, 83/459; female: n = 26.0%, 148/570) and reward (left, male: n = 48.4%, 222/459; female: n = 53.0%, 302/570) across both trial types in male (**b**) and female (**c**) mice. d,g) Heatmaps of the average fluorescence of mPFC neurons in male (**d**, n = 36 neurons) and female (**g**, n = 39 neurons) mice that are significantly excited by social choice aligned to social choice (first panel) and sucrose choice (second panel). Neurons are sorted by the time of peak response to social choice. A row on both heatmaps corresponds to the same neuron. Average fluorescence traces of these social excite neuron responses to social (red) and sucrose (blue) choice (third panel). Comparison of the peak fluorescence of social and sucrose choice responses shows that these neurons on average have an excitatory response to social but not sucrose choice (fourth panel). Paired t test (male: p = 1.47*10-11; female: p = 3.23*10-6). e,h) Heatmaps of the average fluorescence of mPFC neurons in male (**e**, n = 22 neurons) and female (**h**, n = 31 neurons) mice that are significantly excited by sucrose choice aligned to social choice (first panel) and sucrose choice (second panel). Neurons are sorted by the time of peak response to sucrose choice. A row on both heatmaps corresponds to the same neuron. Average fluorescence traces of these social excite neuron responses to social (red) and sucrose (blue) choice (third panel). Comparison of the peak fluorescence of social and sucrose choice responses shows that in male mice these neurons on average have an excitatory response to both sucrose and social choice (**e**, fourth panel). Paired t test (p = 0.45). In contrast, in female mice (**h**, fourth panel) these neurons on average have an excitatory response to sucrose but not social choice. Paired t test (p = 3.61*10-6). f,i) Heatmaps of the average fluorescence of mPFC neurons in male (**f**, n = 21 neurons) and female (**i**, n = 22 neurons) mice that are significantly inhibited by sucrose choice aligned to social choice (first panel) and sucrose choice (second panel). Neurons are sorted by the time of minimum response to sucrose choice. A row on both heatmaps corresponds to the same neuron. Average fluorescence traces of these sucrose inhibit neuron responses to social (red) and sucrose (blue) choice (third panel). Comparison of the peak fluorescence of social and sucrose choice responses shows that these neurons on average have an excitatory response to social choice and an inhibitory response to sucrose choice (fourth panel). Paired t test (male: p = 3.25*10-4; female: p = 0.0042). Shaded error regions indicate ± SEM. Dashed line at zero indicates choice onset.

**Supplementary Figure 10.**
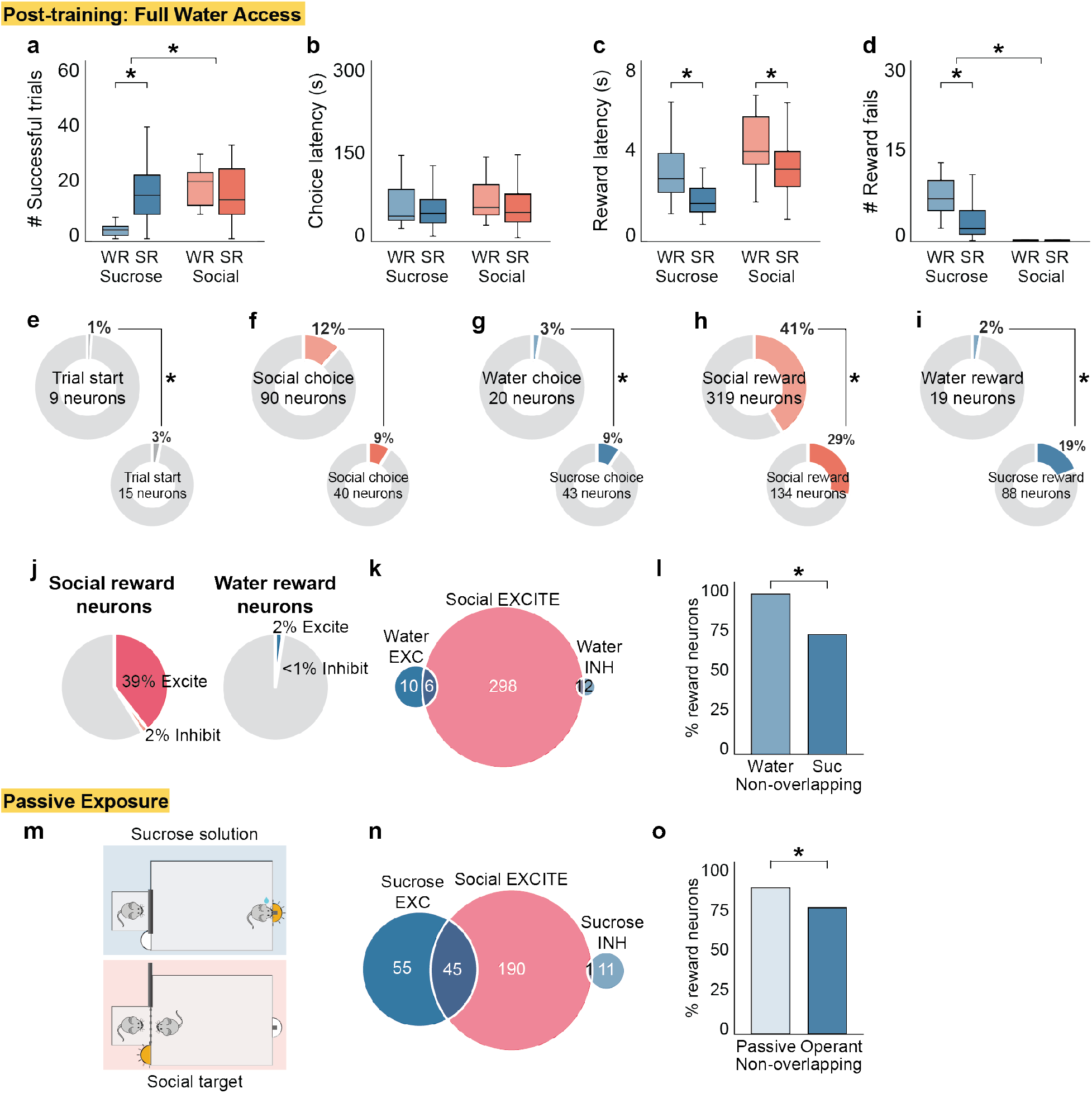
mPFC neural representations of social and nonsocial reward remain non-overlapping when the nonsocial reward is no longer sucrose and with passive exposure to rewarding stimuli. **a**) Under full water access conditions, mice run with water (WR) instead of sucrose on the two choice operant assay complete significantly fewer successful nonsocial reward trials and an equivalent number of successful social reward trials compared to mice run with sucrose (SR). Two-factor ANOVA with reward type (WR/SR) and trial type (sucrose/social) as factors (interaction: p = 1.17*10-5, reward type: p = 0.0029, trial type: p = 4.57*10-5) with post-hoc unpaired t tests comparing reward type within trial type (sucrose: p = 1.13*10-6, social: p = 0.29). b-d) Mice run with WR have similar choice latencies (**b**), are slower to engage with both types of reward (**c**) and make significantly more water reward fails (**d**) than mice run with sucrose (SR). Two-factor ANOVA with reward type (WR/SR) and trial type (sucrose/social) as factors (**b**, interaction: p = 0.64, reward type: p = 0.40, trial type: p = 0.70; **c**, interaction: p = 0.54, reward type: p = 8.17*10-5, trial type: p = 1.60*10-6; **d**, interaction: p = 0.0019, reward type: p = 0.0014, trial type: p = 2.23*10-14) with post-hoc unpaired t tests comparing reward type within trial type (**c**, sucrose: p = 1.43*10-4, social: p = 0.038; **d**, sucrose: p = 0.0017, social: p = 0.74). WR: n = 5 mice, SR: n = 21 mice, 3 sessions per mouse. e-i) Proportions of total recorded neurons that are modulated by the various task events in mice run with water instead of sucrose (WR) compared to mice run with sucrose (SR). Proportion z test with correction for multiple comparisons (**e**, p = 9.59*10-3; **f**, p = 0.11; **g**, p = 1.64*10-7; **h**, p = 2.49*10-5; **i**, p<0.00001). j) Pie charts showing the distribution of social reward responses (left, excite: n = 39.4%, 305/775; inhibit: n = 1.81%, 14/775) and nonsocial reward responses (right, excite: 2.07%, 16/775; inhibit: 0.39%, 3/775) in mice run with water instead of sucrose. k) A venn diagram showing that mPFC neurons are largely non-overlapping in their responsiveness to social and nonsocial reward in mice run with water. l) Neural representations of social and nonsocial reward in mice run with water are even more non-overlapping (n = 97.8%, 310/317 neurons) than those in mice run with sucrose reward (n = 73.1%, 125/171). Proportion z test (p<0.00001). WR: n = 775 neurons, 5 mice; SR: n = 459 neurons, 9 mice. m) Schematic of the passive exposure paradigm in which the gate opens to allow access to a social target or the sucrose reward port activates to allow access to sucrose solution at random intervals over a one hour session. n) A venn diagram showing that mPFC neurons are largely non-overlapping in their responsiveness to social and nonsocial stimuli in mice that are passively exposed to either a social target or sucrose solution. o) Passive exposure to social and nonsocial stimuli results in significantly more non-overlapping social and nonsocial reward representations (n = 84.8%, 256/302 neurons) than those in mice run on the two choice operant assay (n = 73.1%, 125/171). Proportion z test (p = 0.0021). Passive exposure: n = 780 neurons, 5 mice; SR: n = 459 neurons, 9 mice. *p<0.05. Box plots: center line denotes median, box edges indicate the 25th and 75th percentiles and whiskers extend to ± 2.7σ.

**Supplementary Figure 11.**
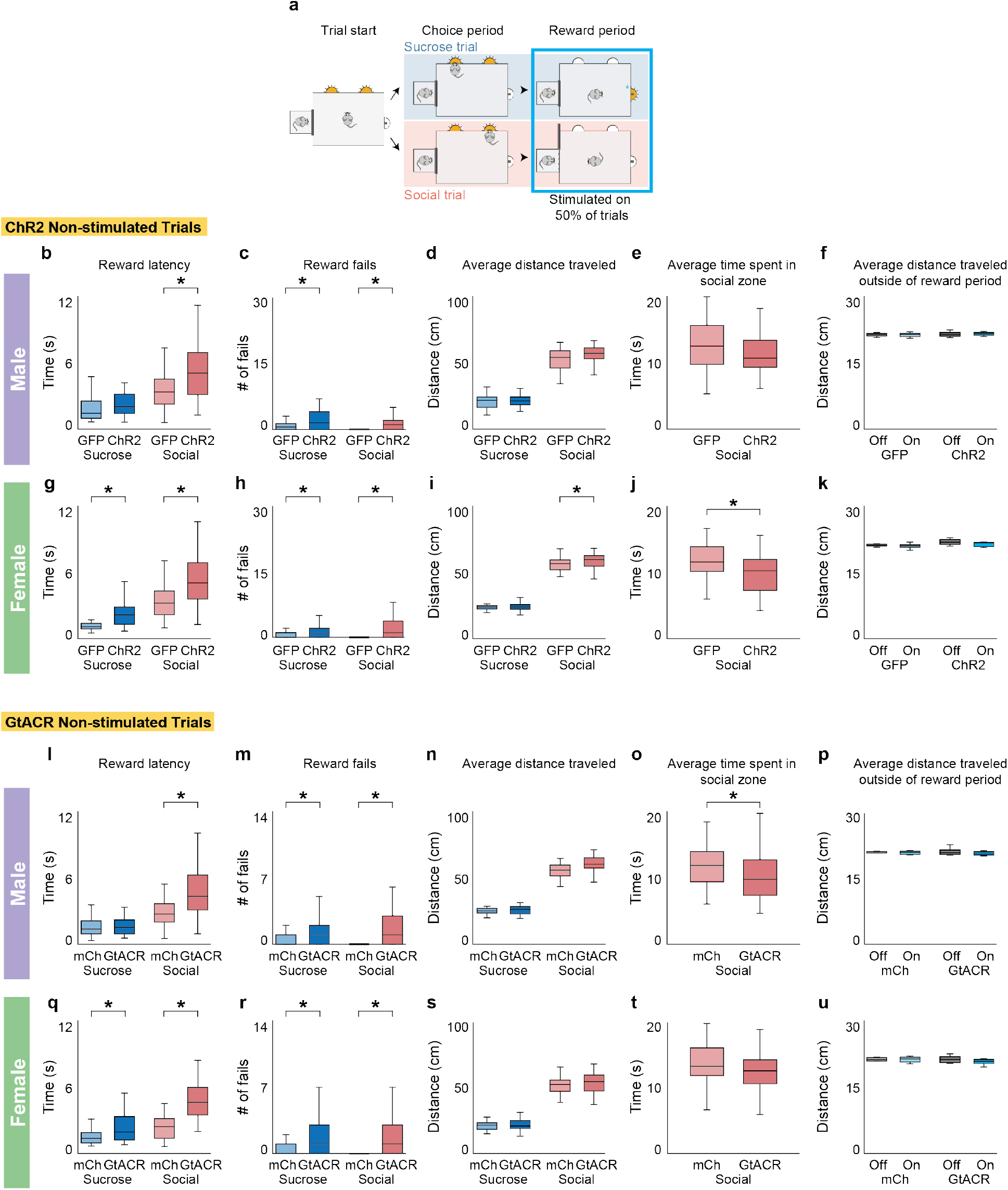
Optogenetic activation and inhibition of mPFC neurons disrupts reward-seeking behavior on trials without laser stimulation. **a**) Schematic of experimental strategy for optogenetic activation and inhibition experiments. Stimulation (ChR2/GFP experiments: 20 Hz, 1-3 mW, 5 ms pulse duration; GtACR/mCherry experiments: 10 mW, 8s pulse duration) during the reward period begins once mice make a choice and lasts for 8s on a random 50% of trials. b,c,g,h) Male ChR2 mice show increased reward latency (**b**) and reward fails (**c**) on social, but not sucrose trials without laser stimulation compared to GFP controls, while female ChR2 mice show increased reward latency (**g**) and reward fails (**h**) on both social and sucrose trials without laser stimulation compared to GFP controls. Two-factor ANOVA with virus (ChR2/GFP) and trial type (sucrose/social) as factors (**b**, interaction: p = 0.016, virus: p = 8.00*10-5, trial type: p = 4.62*10-16; **c**, interaction: p = 0.96, virus: p = 6.82*10-5, trial type: p = 0.0015, **g**, interaction: p = 0.11, virus: p = 2.61*10-9, trial type: p = 5.12*10-23; **h**, interaction: p = 0.024, virus: p = 2.23*10-6, trial type: p = 0.83) with post-hoc unpaired t tests comparing virus within trial type (**b**, sucrose: p = 0.074, social: p = 5.31*10-4; **c**, sucrose: p = 0.025, social: p = 2.50*10-5; **g**, sucrose: p = 1.61*10-8, social: p = 6.04*10-5; **h**, sucrose: p = 0.044, social: p = 1.07*10-5). d,i) Male ChR2 and GFP mice travel a comparable distance on sucrose and social trials without laser stimulation (**d**), while female ChR2 mice travel a further distance on social but not sucrose trials without laser stimulation compared to GFP controls (**i**). Two-factor ANOVA with virus (ChR2/GFP) and trial type (sucrose/social) as factors (**d**, interaction: p = 0.15, virus: p = 0.037, trial type: p = 4.71*10-89; **i**, interaction: p = 0.084, virus: p = 0.0052, trial type: 5.49*10-120) with post-hoc unpaired t tests comparing virus within trial type (**d**, sucrose: p = 0.97, social: p = 0.059; **i**, sucrose: p = 0.86, social: p = 0.0076). e,j) Male ChR2 mice and GFP control mice spend a comparable amount of time in the social zone during the reward period on trials without laser stimulation (**e**), but female ChR2 mice spend less time in the social zone during the reward period on trials without laser stimulation compared to GFP controls (**j**). Unpaired t test (male: p = 0.29; female: p = 8.02*10-4). f,k) Male and female ChR2 mice traveled the same distance compared to GFP controls when stimulated outside of the reward period on the two choice assay. Two-factor ANOVA with virus (ChR2/GFP) and stimulation (on/off) as factors (**f**, interaction: p = 0.55, virus: p = 0.28, stimulation: p = 0.74; **k**, interaction: p = 0.36, virus: p = 7.46*10-4, stimulation: p = 0.11). l,q) Male GtACR mice show increased reward latency (**l**) on social, but not sucrose trials without laser stimulation compared to mCherry controls, while female GtACR mice show increased reward latency (**q**) on both social and sucrose trials without laser stimulation compared to mCherry controls. Two-factor ANOVA with virus (GtACR/mCherry) and trial type (sucrose/social) as factors (**l**, interaction: p = 0.0004, virus: p = 0.0001, trial type: p<0.00001; **q**, interaction: p = 1.78*10-12, virus: p = 2.51*10-4, trial type: p = 2.24*10-14) with post-hoc unpaired t tests comparing virus within trial type (**l**, sucrose: p = 0.60, social: q = 2.14*10-5; **q**, sucrose: p = 1.94*10-4, social: p = 4.25*10-9). m,n,r,s) Both male and female GtACR mice make significantly more social and sucrose reward fails (**m**,**r**) on trials without laser stimulation compared to mCherry controls, but show no difference in the average distance traveled (**n**,**s**). Two-factor ANOVA with virus (GtACR/mCherry) and trial type (sucrose/social) as factors (**m**, interaction: p = 0.22, virus: p<0.00001, trial type: p = 0.38; **n**, interaction: p = 0.69, virus: p = 0.57, trial type: p = 2.00*10-67; **r**, interaction: p = 0.74, virus: p<0.00001, trial type: p = 0.31; **s**, interaction: p = 0.67, virus: p = 0.20, trial type: p = 2.94*10-84) with post-hoc unpaired t tests comparing virus within trial type (**m**, sucrose: p = 0.027, social: p = 3.04*10-5; **r**, sucrose: p = 0.0076, social: p = 3.12*10-5). o,t) Female GtACR and mCherry control mice spend a comparable amount of time in the social zone during the reward period on trials without laser stimulation (**t**), but male GtACR mice spend less time in the social zone during the reward period on trials without laser stimulation compared to mCherry controls (**o**). Unpaired t test (male: p = 0.020; female: p = 0.12). p,u) Male and female GtACR mice traveled the same distance compared to mCherry controls when stimulated outside of the reward period on the two choice assay. Two-factor ANOVA with virus (GtACR/mCherry) and stimulation (on/off) as factors (**p**, interaction: p = 0.38, virus: p = 0.85, stimulation: p = 0.25; **u**, interaction: p = 0.22, virus: p = 0.45, stimulation: p = 0.22). *p<0.05. Box plots: center line denotes median, box edges indicate the 25th and 75th percentiles and whiskers extend to ± 2.7σ.

**Supplementary Figure 12.**
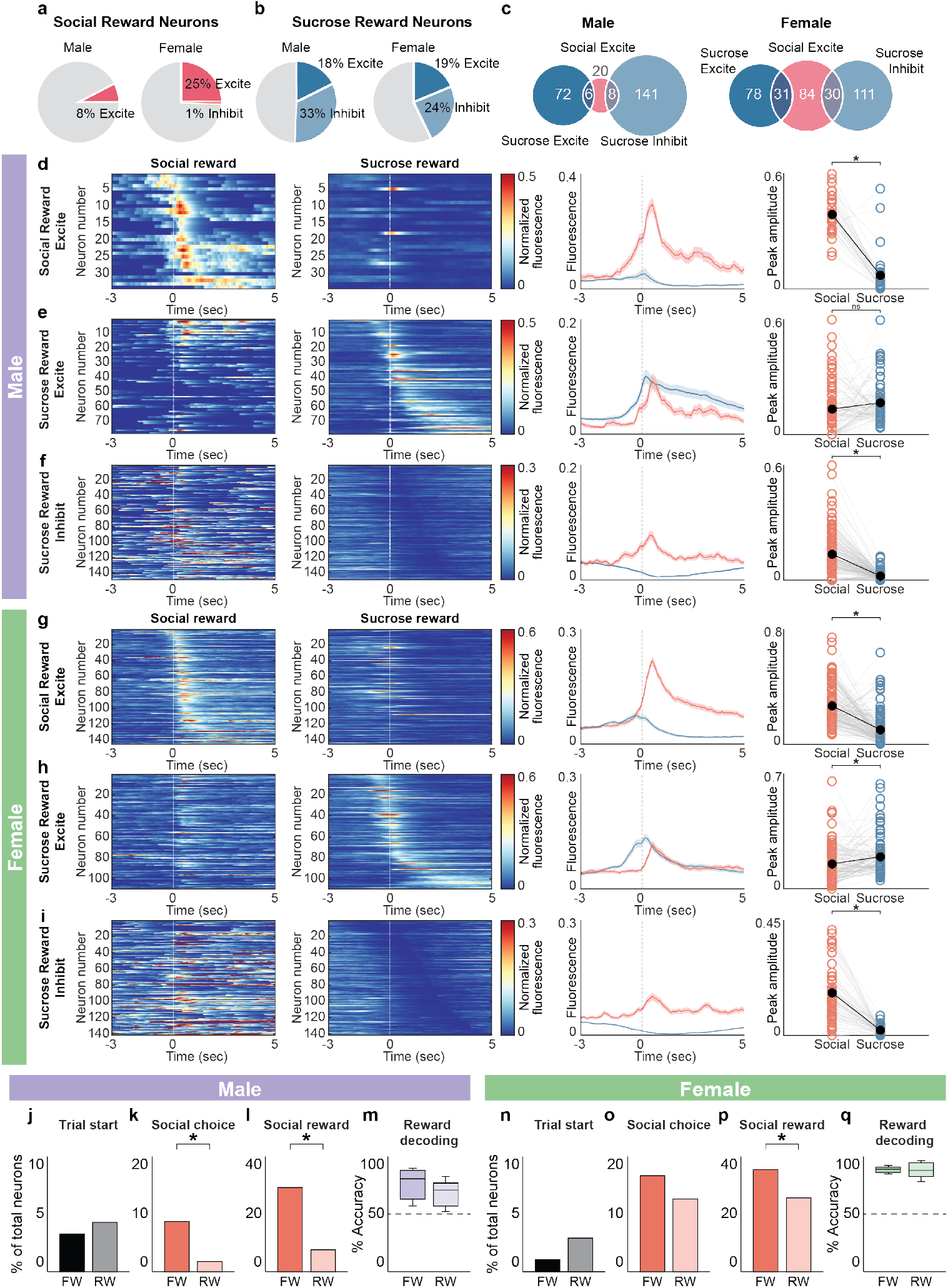
mPFC neurons maintain sex-dependent selectivity of social and nonsocial reward responses with restricted water access. **a**) Pie charts showing the distribution of social reward responses in male (left) and female (right) mice on restricted water access (RW). b) Pie charts showing the distribution of sucrose reward responses in male (left) and female (right) mice on RW. c) Venn diagrams showing that largely non-overlapping populations of mPFC reward neurons respond to social and sucrose reward in male (left) and female (right) mice on RW. d,g) Heatmaps of the average fluorescence of mPFC neurons in male (**d**, n = 34 neurons) and female (**g**, n = 145 neurons) mice that are significantly excited by social reward on RW aligned to social reward (first panel) and sucrose reward (second panel). Neurons are sorted by the time of peak response to social reward. A row on both heatmaps corresponds to the same neuron. Average fluorescence traces of these social excite neuron responses to social (red) and sucrose (blue) reward (third panel). Comparison of the peak fluorescence of social and sucrose reward responses shows that these neurons on average have a robust excitatory response to social but not sucrose reward in male and female mice. Paired t test (male: p = 5.48*10-14; female: p = 3.48*10-20). e,h) Heatmaps of the average fluorescence of mPFC neurons in male (**e**, n = 78 neurons) and female (**h**, n = 109 neurons) mice that are significantly excited by sucrose reward on RW aligned to social reward (first panel) and sucrose reward (second panel). Neurons are sorted by the time of peak response to sucrose reward. A row on both heatmaps corresponds to the same neuron. Average fluorescence traces of these sucrose excite neuron responses to social (red) and sucrose (blue) reward (third panel). Comparison of the peak fluorescence of social and sucrose reward responses shows that these neurons respond similarly to social and sucrose rewards in male mice. In contrast, these neurons respond more robustly to sucrose reward than social reward in female mice. Paired t test (male: p = 0.12; female: p = 4.28*10-3). f,i) Heatmaps of the average fluorescence of mPFC neurons in male (**f**, n = 149 neurons) and female (**i**, n = 141 neurons) mice that are significantly inhibited by sucrose reward on RW aligned to social reward (first panel) and sucrose reward (second panel). Neurons are sorted by the time of minimum response to sucrose reward. A row on both heatmaps corresponds to the same neuron. Average fluorescence traces of these sucrose inhibit neuron responses to social (red) and sucrose (blue) reward (third panel). Comparison of the peak fluorescence of social and sucrose reward responses that these neurons on average have an excitatory response to social reward and an inhibitory response to sucrose reward in both males and females. Paired t test (male: p = 5.12*10-26; female: p = 3.53*10-25). j–l) No significant difference was observed in the proportion of trial start neurons (**j**, 4.3%, n = 19/444 RW vs 3.3% FW) across both conditions in male mice. Proportion z test with correction for multiple comparisons (trial start: p = 0.43). During restricted water access conditions (RW), a significantly smaller proportion of mPFC neurons responded to social choice (**k**, 1.8%, n = 8/444) and social reward (**l**, 7.7%, n = 34/444) when compared to full water access conditions in male mice (social choice: 8.7%, social reward = 29.2%, FW). Proportion z test with correction for multiple comparisons (social choice: p = 3.67*10-6, social reward: p<0.00001). m) Decoders trained on male mPFC neural reward responses during FW and RW access conditions decoded the reward type with equivalent accuracy. Paired t test (p = 0.21). Decoding accuracy was calculated for each animal using neurons tracked across FW and RW access conditions, with a trial-matched number of sucrose and social trials (n = 178 neurons, 8 mice). All decoding was significantly greater than shuffled data, indicated by a dashed line at 50%. n-p) During restricted water access conditions (RW), a similar proportion of mPFC neurons respond to trial start (**n**, 2.9%, n = 17/585) and social choice (**o**, 12.7%, n =74/585) when compared to full water access conditions in female mice (trial start: 1,1%, social choice: 16.7%, FW). Proportion z test with correction for multiple comparisons (trial start: p = 0.024, social choice: p = 0.054). A significantly smaller proportion of mPFC neurons responded to social reward in RW (**p**, 25.6%, n = 150/585) compared to FW (35.4%). Proportion z test with correction for multiple comparisons (p = 2.99*10-4). q) Decoders trained on female mPFC neural reward responses during FW and RW access conditions decoded the reward type with equivalent accuracy. Paired t test (p = 0.91). Decoding accuracy was calculated for each animal using neurons tracked across FW and RW access conditions, with a trial-matched number of sucrose and social trials (n = 226 neurons, 5 mice). All decoding was significantly greater than shuffled data, indicated by a dashed line at 50%. *p<0.05. Shaded error regions indicate ± SEM. Dashed line at zero indicates reward onset. Box plots: center line denotes median, box edges indicate the 25th and 75th percentiles and whiskers extend to ± 2.7σ.

**Supplementary Figure 13.**
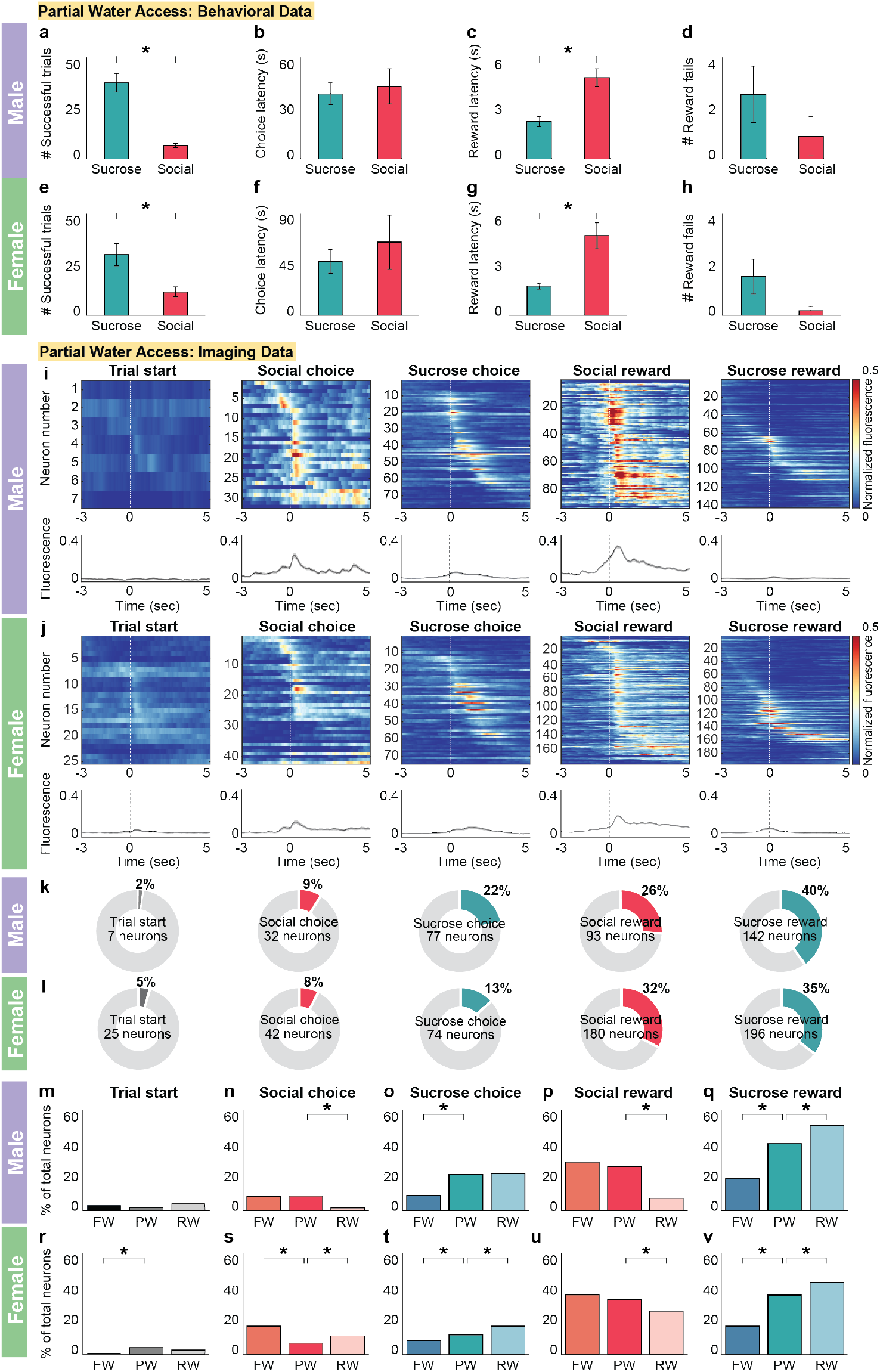
Reward-seeking behavior and neural representations of reward display an intermediate phenotype during partial water access conditions. **a,e**) Under partial water access conditions, male (**a**) and female (**e**) mice complete a greater number of successful sucrose trials compared to social trials. Paired t test (male: p = 5.77*10-5; female: p = 0.044). b,f) Male (**b**) and female (**f**) mice have comparable choice latencies across both trial types. Paired t test (male: p = 0.54; female: p = 0.52). c,g) Under partial water access conditions, male (**c**) and female (**g**) mice show decreased sucrose reward latency compared to social reward latency. Paired t test (male: p = 0.0016; female: p = 0.014). d,h) Male (**d**) and female (**h**) mice make a similar number of sucrose and social reward fails. Paired t test (male: p = 0.29; female: p = 0.30). i,j) Top row: Heatmaps of average normalized fluorescence of all mPFC neurons that are significantly modulated (excited or inhibited) by each task event in male (**i**) and female (**j**) mice during partial water access conditions. Neurons are sorted by the time of maximum fluorescence across each task event. Bottom row: Average normalized fluorescence traces of the neurons from the corresponding heatmap that are significantly modulated by each task event. Shaded error regions indicate ± SEM. k,l) Proportions of mPFC neurons that are modulated by the various task events in male (**k**) and female (**l**) mice under partial water access conditions. Proportions are determined using the total number of recorded neurons (male: n = 355 neurons, 9 mice; female: n = 556 neurons, 6 mice). m-q) Comparison of proportion of mPFC neurons that are significantly modulated by each task event in male mice across three water access conditions. Under partial water access conditions (PW), male mice have a similar proportion of mPFC neurons significantly modulated by social choice (**n**) and social reward (**p**) compared to full water access (FW) but not restricted water access conditions (RW), whereas male mice have a similar proportion of mPFC neurons significantly modulated by sucrose choice (**o**) across RW but not FW. Proportion z test (social choice - FW vs PW: p = 0.88, PW vs RW: p = 3.40*10-6; sucrose choice - FW vs PW: p = 8.77*10-7, PW vs RW: p = 0.84; social reward - FW vs PW: p = 0.34, PW vs RW: p = 1.07*10-12). The proportion of mPFC neurons modulated by sucrose reward (**q**) during PW is significantly greater than FW but less than RW. Proportion z test (FW vs PW: p = 5.95*10-11, PW vs RW: p = 0.0026). There is no change in proportion of mPFC neurons modulated by trial start (**m**) across all three water access conditions. Proportion z test (FW vs PW: p = 0.26, PW vs RW: p = 0.068). r-v) Comparison of proportion of mPFC neurons that are significantly modulated by each task event in female mice across three water access conditions. Under partial water access conditions (PW), female mice have a different proportion of mPFC neurons significantly modulated by social choice (**s**), sucrose choice (**t**) and sucrose reward (**v**) compared to both full water (FW) and restricted water (RW) access conditions. Proportion z test (social choice - FW vs PW: p = 2.92*10-6, PW vs RW: p = 0.0044; sucrose choice - FW vs PW: p = 0.033, PW vs RW: p = 6.28*10-9; sucrose reward - FW vs PW: p = 1.49*10-11, PW vs RW: p = 0.0096). The proportion of mPFC neurons modulated by social reward (**u**) was significantly different between PW and RW but not FW. Proportion z test (FW vs PW: p = 0.28, PW vs RW: p = 0.012). The proportion of mPFC neurons modulated by trial start (**r**) was significantly different between PW and FW but not RW. Proportion z test (FW vs PW: p = 0.00041, PW vs RW: p = 0.15). Box plots: center line denotes median, box edges indicate the 25th and 75th percentiles and whiskers extend to ± 2.7σ.

**Supplementary Figure 14.**
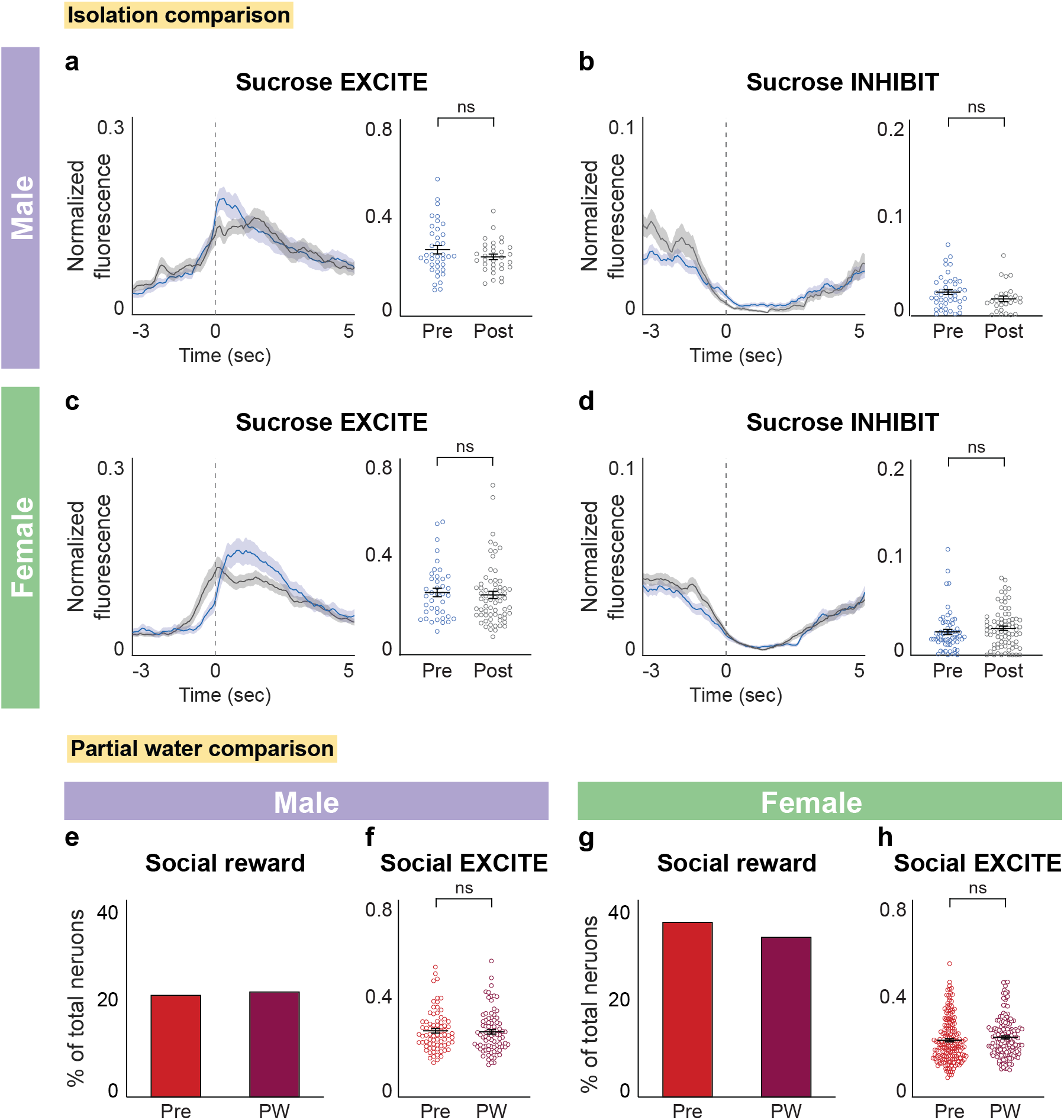
Changes in reward representations in the mPFC following social isolation are specific to social reward in male and female mice. **a,b**) Average normalized fluorescence traces of mPFC neurons significantly excited (**a**) and inhibited (**b**) by sucrose reward before (blue) and after (gray) social isolation in male mice time-locked to sucrose reward (left panel). Comparison of peak fluorescence before (sucrose excite: n = 38 neurons; sucrose inhibit: n = 44 neurons) and after (sucrose excite: n = 33 neurons; sucrose inhibit: n = 28 neurons) social isolation shows that neural responses to sucrose reward remain stable with social isolation in male mice (right panel). Unpaired t test (sucrose excite: p = 0.17; sucrose inhibit: p = 0.077). c,d) Average normalized fluorescence traces of mPFC neurons significantly excited (**c**) and inhibited (**d**) by sucrose reward before (blue) and after (gray) social isolation in female mice time-locked to sucrose (left panel). Comparison of peak fluorescence before (sucrose excite: n = 40 neurons; sucrose inhibit: n = 60 neurons) and after (sucrose excite: n = 67 neurons; sucrose inhibit: n = 79 neurons) social isolation shows that neural responses to sucrose reward remain stable with social isolation in male mice (right panel). Unpaired t test (sucrose excite: p = 0.66; sucrose inhibit: p = 0.30). e-h) Similarly to social isolation, the proportion of total neurons that are significantly modulated by social reward does not change between partial water and full water access in male (**e**) and female (**g**) mice. Proportion z test (male: p = 0.81; female: p = 0.28). However, unlike following social isolation, the peak amplitude of social excite neurons remains stable across partial (male: n = 84 neurons; female: n = 164 neurons) and full water access conditions (male: n = 76 neurons; female: n = 177 neurons) in male (**f**) and female (**h**) mice. Unpaired t test (male: p = 0.74; female: p = 0.21). Shaded error regions indicate ± SEM.

## Methods

### Experimental Subjects

For all experiments, C57BL/6J male and female mice aged 6-10 weeks from Jackson laboratories were used. Mice were maintained on a reverse light dark cycle (12-hour light off-12-hour light on schedule). All experimental procedures were approved by the Emory Institutional Animal Care and Use Committee.

### Viruses

The viruses used in this study were purchased from Addgene: AAV5-Syn-GCaMP6f-WPRE-SV40 (100835-AAV5), AAV5-hSyn-hChR2(H134R)-EYFP (26973-AAV5), AAV5-CAG-GFP (37825-AAV5), AAV1-CKIIa-stGtACR2-FusionRed (105669-AAV1), and AAV5-CKIIa-mCherry (114469-AAV5).

### Stereotaxic Surgeries

For imaging surgeries, mice aged ∼6-8 weeks (n = 19 male mice, 6 female mice) were anesthetized with 1-2% isoflurane and placed in stereotactic setup (Kopf). A microsyringe (Nanoject) was used to inject 0.75 µL of AAV5-Syn-GCaMP6f-WPRE-SV40 (Addgene, injected titer of 1.3×10^13 parts/ml) unilaterally into the mPFC (1.6 mm anterior, 0.5 mm lateral and 2.4 mm in depth). Mice were then implanted with a 0.6 mm diameter, 7.3 mm length GRIN lens (Product ID:1050-004597, Inscopix) over the mPFC (1.8 mm anterior, 0.5 mm lateral and 2.1 mm in depth). A metal cap was cemented over the lens to protect it. Mice were group housed following lens implant. 3-4 weeks following viral injection and lens implant, a baseplate (Product ID: 1050-004638, Inscopix) attached to the miniature microscope (nVista, Inscopix) was positioned ∼0.45 mm above the lens such that blood vessels and neurons were in focus. The baseplate was then cemented in place using dental cement (Metabond) and a baseplate cover (Product ID: 1050-004639, Inscopix) was secured over the baseplate to protect the lens.

For optogenetic activation surgeries, mice aged ∼6-8 weeks (n = 14 male mice, 14 female mice) were anesthetized with 1-2% isoflurane and placed in stereotactic setup (Kopf). A microsyringe (Nanoject) was used to inject 0.35 µL of AAV5-hSyn-hChR2(H134R)-EYFP (experimental, Addgene, injected titer of 2.4×10^13 parts/ml, n = 8 male mice, 8 female mice) or AAV5-CAG-GFP (control, Addgene, injected titer of 1.3×10^13 parts/ml, n = 6 male mice, 6 female mice) bilaterally into the mPFC (1.6 mm anterior, 0.5 mm lateral and 2.4 mm in depth). Mice were then implanted with custom-made optic ferrules consisting of a 2.5 mm diameter metal ferrule sleeve (Product ID: MM-FER2003SS-3300, Precision Fiber Products) and a 0.3 mm diameter optic core (Product ID: FT300UMT, ThorLabs) bilaterally at a 10 degree angle into the mPFC above the viral injection (1.8 mm anterior, 0.7 mm lateral and 2.15 mm in depth).

For optogenetic inhibition surgeries, mice aged ∼6-8 weeks (n = 13 male mice, 13 female mice) were anesthetized with 1-2% isoflurane and placed in stereotactic setup (Kopf). A microsyringe (Nanoject) was used to inject 0.35 µL of AAV1-CKIIa-stGtACR2-FusionRed (experimental, Addgene, injected titer of 1.3×10^13 parts/ml, n = 7 male mice, 7 female mice) or AAV5-CKIIa-mCherry (control, Addgene, injected titer of 2.3×10^13 parts/ml, n = 6 male mice, 6 female mice) bilaterally into the mPFC (1.6 mm anterior, 0.5 mm lateral and 2.4 mm in depth). Mice were then implanted with custom-made optic ferrules consisting of a 2.5 mm diameter metal ferrule sleeve (Product ID: MM-FER2003SS-3300, Precision Fiber Products) and a 0.3 mm diameter optic core (Product ID: FT300UMT, ThorLabs) bilaterally at a 10 degree angle into the mPFC above the viral injection (1.8 mm anterior, 0.7 mm lateral and 2.15 mm in depth).

### Behavioral Assays

#### Automated Operant Assay

We developed an automated, closed loop operant assay to quantitatively assess social and nonsocial reward behaviors. The assay consists of an acrylic chamber (12×12×12 inches) with social and nonsocial (sucrose) reward access on opposing sides and two nose-poke ports (choice ports) on the wall adjacent to both reward access sites (Figure 1a). The two choice ports (Product ID: 1009, Sanworks BPOD) are located 3 inches from the corners and 1 inch from the base of the chamber. The sucrose reward port (Product ID: 1009, Sanworks BPOD) is centered on one adjacent wall and contains a solenoid valve connected to a liquid reservoir containing 10% sucrose in water. On the wall opposite from the sucrose reward port, a 2×2 inch square cutout provides access to the social target chamber. The cutout is covered on the exterior side of the chamber by a flat sheet of aluminum fixed to a conveyor belt with a stepper motor which acts as a movable gate, controlled by an Arduino Uno. Behind this cutout, is a smaller acrylic chamber (3×3×3 inch) with an exposed side blocked by 6 aluminum rods (⅕ cm in diameter and 0.5 cm apart) that holds the target animal. Social target animals were non-cagemate, age- and sex-matched conspecifics. Nose pokes are detected by infrared beam break-sensors. A trial begins when both choice ports illuminate and is followed by nose-poking in either choice port to initiate social or nonsocial reward delivery. Nose-poking of the choice port furthest from the sucrose reward port illuminates the reward port and makes 10 µL of sucrose available upon entry into the reward port for up to 8s. Alternatively, nose-poking of the choice port furthest from the social target chamber, opens the gate for 20s, which allows the experimental animal to interact with the social target through the aluminum bars without permitting entry into each other’s chamber. Following sucrose or social reward consumption, the trial ends and after a 30 s intertrial interval (ITI), a new trial begins with the illumination of both choice ports. Each behavioral session is one hour in duration. Our social operant assay consists of several phases: training (4 stages, ∼2 weeks) and post-training (up to 3 weeks).

#### Training Paradigms

Mice were co-housed and placed on a restricted water schedule prior to the start of training. At the end of each day mice were provided 1.5 mL of water in addition to their consumption during training.

##### Training stage 1: Operant conditioning

In the first stage (operant conditioning), mice were trained to associate poking the reward port with a sucrose reward. Nose port entry triggered the port LED light on and resulted in the delivery of a sucrose reward of 10 µL of 10% sucrose in water. Each session lasted an hour and animals were run through the assay once a day. Mice had to reach a minimum of 50 pokes in a single session to progress to the next training stage. If the animal did not engage with the reward port for more than two minutes after the start of the trial, then the reward port delivered 30 µL liquid without engaging the LED. This automatic reward delivery also occurred at the beginning of the first trial in each session to prime the animal to engage with the port.

##### Training stage 2: Single port one choice

In the second stage (single port one choice), mice were conditionally rewarded with 10 µL of sucrose for poking the reward port within 8s of the reward port LED turning on. Trials were separated by a 30s ITI, and the animal had to refrain from entering the port for 3s for the next trial to start. A trial was considered a sucrose reward fail if the mouse failed to enter the sucrose reward port after it had been illuminated for 8s. Mice had to reach a minimum of 40 successful trials during one session to proceed to the next training stage.

##### Training stage 3: Opposing port one choice

In the third stage (opposing port one choice), the choice port LED furthest from the sucrose reward port (sucrose choice port) turned on to indicate trial start. This assay is self-paced, so mice had an indefinite amount of time to nose poke the illuminated sucrose choice port. A successful choice port poke resulted in the LED flashing on and off successively 3 times in 0.1s intervals, after which the reward port LED turned on for up to 8s. Mice had to nose poke the reward port within this window to receive a 10 µL sucrose reward. Failure to do so resulted in a reward fail and initiation of a new trial. Trials were separated by a 15s ITI. In addition, the current trial ended if the animal engaged with the other inactive port instead of the sucrose choice port during the choice period. Mice had to refrain from entering the sucrose choice port for 3s for the next trial to start. Mice had to perform 40 successful sucrose reward trials to proceed to the final stage of training.

##### Training stage 4: Two choice (social-sucrose) operant assay

In the final stage (two choice operant), mice on the restricted water schedule were given 500ul of water prior to the start of each session (partial water access condition). As with the previous training stage, trial start was indicated by the choice port LED turning on. However, unlike the previous training stage, a second choice port was activated so that the animal could choose between social and nonsocial rewards. Nose-poking the illuminated choice port furthest from the sucrose reward port (sucrose choice port) resulted in the LED flashing on and off successively 3 times in 0.1s intervals, after which the sucrose reward port LED turned on for 8s. Mice had to enter the sucrose reward port within this window to receive a 10 µL sucrose reward. Failure to do so resulted in a sucrose reward fail. Nose-poking the illuminated choice port furthest from the social target chamber (social choice port) resulted in the LED flashing on and off successively 3 times in 0.1s intervals, after which an automated gate opened for 20s to provide the mouse access to a social target (non-cagemate, sex- and age-matched conspecific) in a smaller chamber. The social target chamber contains an open side covered with aluminum bars that are 0.5 cm apart. This allows mice to freely interact, smell and investigate each other without permitting either animal to enter the other chamber. The gate remained open for 20s before closing. Failure to investigate the social target in this time resulted in a social reward fail. Trials were separated by a 30s ITI. Mice were randomly assigned to one of two configurations of the operant chamber in which the social and sucrose rewards were on different sides in order to account for baseline side preferences that may arise.

##### Post-training: Two choice (social-sucrose) operant assay

Once fully trained, mice were removed from the limited water schedule and had ad libitum access to water outside of behavioral sessions. They were maintained on this full water access (FW) schedule and were run daily on the two choice operant assay for a week. For restricted water access (RW) experiments, animals were placed on the limited water schedule (1.5 mL of water/day) without access to 500 µL of water before sessions and run daily on the two choice operant assay for a week. For social isolation experiments, mice were fully trained and run daily for one week on the full water schedule. Mice were then single housed for one week and not run on the assay. After one week, mice were run again for one session on the two choice operant assay.

##### Empty cage/object control

After mice had been run through the first three training stages, a subset of the mice were run on the final stage of training with either an empty social target cage (empty cage group) or a novel object (object group) (Supplementary Figure 4). Mice were then removed from the limited water schedule, given ad libitum water access, and run daily for several days on the full water access schedule without a social target and with either an empty cage or a novel object. One mouse from the empty cage group was excluded from full water access analysis due to failure to engage with the task.

##### Multi-choice operant assay

A third choice port (null port) was added to the two choice operant assay (Supplementary Figure 5a). Nose-poking the null port resulted in neither a social nor a sucrose reward (null trial). A subset of mice were run on this assay through similar training stages as the two choice operant assay. Mice were then removed from the limited water schedule, given ad libitum water access, and run daily for nine days on the full water access schedule.

##### Estrous cycle monitoring

A subset of the female mice underwent vaginal lavage after each behavioral session. The cytological evidence from each lavage was used to determine the daily estrous phase of these female mice (Supplementary Figure 6a,b).

##### Water reward control

After mice had been run through the first three training stages, a subset of the mice were run on the final stage of training in which the sucrose reward had been replaced with water (Supplementary Figure 10a-l). When the mouse nose-poked the nonsocial reward port, 10 µL of water were dispensed instead of 10 µL of sucrose solution. Mice were then removed from the limited water schedule, given ad libitum water access, and run daily for several days on the full water access schedule while receiving a water reward when performing a nonsocial trial.

##### Passive exposure

Mice were placed in a behavioral arena and passively exposed to either a non-cagemate, sex- and age-matched social target through the opening of an automated gate or a 10 µL droplet of a sucrose solution from a sucrose port (Supplementary Figure 10m-o). Passive exposure type was randomized and there were at least 30s between exposures.

### Definition of task events and parameters

***Trial start*** - the time point when both choice ports illuminate.

***Social choice*** - the time point when the experimental mouse enters (nose-pokes) the social choice port after a trial starts.

***Sucrose choice*** - the time point when the experimental mouse enters (nose-pokes) the sucrose choice port after a trial starts.

***Social choice latency*** - the time between the trial start and social choice port entry. Entry into port (nose-poke) is detected by an infrared beam break.

***Sucrose choice latency*** - the time between the trial start and sucrose choice port entry. Entry into port (nose-poke) is detected by an infrared beam break.

***Social reward zone*** - a 1×3 inch rectangular region within the operant arena adjacent to and centered around the social target access point where the gate opens.

***Social reward consumption*** - the time point when the experimental mouse first enters the social reward zone following a social choice. Entry into the social reward zone is detected from the behavioral video using Bonsai software^83^. A mouse is considered to have “consumed” a social reward if it enters the social reward zone within 20s after social choice port entry.

***Sucrose reward consumption*** - the time point when the experimental mouse first enters the sucrose reward port following a sucrose choice. Entry into the sucrose reward port is detected by an infrared beam break. A mouse is considered to have “consumed” a sucrose reward if it enters the sucrose reward port within 8s after sucrose choice port entry.

***Social reward latency*** - the time between social choice port entry and the first entry into the social reward zone.

***Sucrose reward latency*** - the time between sucrose choice port entry and the first entry into the sucrose reward port.

***Social reward fail*** - a trial in which the experimental mouse fails to enter the social reward zone within 20s (duration for which the social gate is open) following social choice.

***Sucrose reward fail*** - a trial in which the experimental mouse fails to enter the sucrose reward port within 8s (duration of sucrose reward port activation) following sucrose choice.

***Successful trial -*** a trial in which the experimental mouse entered the respective reward zone following choice port entry while the reward was available (20s for social trials, 8s for sucrose trials). On the multi-choice assay (Supplementary Figure 5a), all null trials were considered successful.

### Behavioral Analysis

All behavioral sessions were recorded at 40 Hz using Pylon software. Following behavioral sessions, videos were analyzed and behavior was quantified. Several metrics were used to quantify mouse behavior for a given behavioral session, including number of pokes per reward type, choice latency and reward latency. Poke number was defined as the total number of times that an animal nose-poked a choice port, independent of reward consumption. Number of successful trials was determined as the total number of times that an animal nose-poked a choice port and entered the appropriate reward zone when the reward was available for consumption.

Choice latency was defined as the time between when the choice port LEDs turned on (trial start) and the animal nose-poked either choice port. Following each choice port nose-poke, the chosen port would flash and the reward would be made available for consumption. Reward latency was defined as the time from the start of the presentation of the reward to the consumption of the reward. Sucrose reward consumption start was measured as the first entry of the mouse into the reward port after it was activated by a sucrose choice port nose-poke. Social reward consumption start was measured as the first entry of the mouse into a 1×3 inch area (social zone) in front of the opened social gate following a social choice port nose-poke. Social zone entries were tracked through behavioral video analysis using Bonsai software^83^. Failure to consume a reward within the allotted time following reward presentation was considered a reward fail.

To quantify the time to investigate a social target after the reward was presented (social latency), behavioral video data from each session were analyzed using Bonsai software^83^. We used the software to track start and end of trials by detecting changes in light levels of the choice port LEDs. Additionally, we tracked the body position (centroid) of the mouse in the chamber and the mouse’s entry into either reward area (1×3 inches) around the social target and sucrose reward port. Reward latency was quantified as the time elapsed between turning off of the choice port LED and the animal’s first entrance into the corresponding reward zone. This analysis was combined with existing session data from Bpod to quantify the social and nonsocial reward behavior of each mouse on all behavioral (training and post-training) sessions.

### Endoscopic Calcium Imaging Experiments and Analysis

∼1 week post injection, GRIN lens (0.6 mm diameter, 7.3 mm length) implanted mice were trained sequentially on each behavioral assay. Once the mice had been trained for ∼2 weeks, they were implanted with the baseplate and habituated with a tethered dummy microscope (Product ID: 1050-003762, Inscopix) during behavioral sessions for at least 3 days prior to the actual imaging session. On the day of imaging, mice were allowed to habituate to the scope for 10 minutes prior to the start of the session. During the behavioral imaging session, the LED power was set to 0.7 and the analog gain on the image sensor was set to 1.6-1.8. Images were acquired at 20 Hz using Inscopix nVista software. Following imaging acquisition, data were spatially downsampled by a factor of 4 and motion corrected using Inscopix software. A CNMFe algorithm^46^ was then used to identify individual neurons and their fluorescence traces. The fluorescence traces were subsequently aligned to the behavioral data and used for task modulation analysis. One animal was excluded from calcium imaging analysis due to failure to express GCaMP6f. Custom MATLAB software was used for all imaging data analysis.

### Optogenetic Activation Experiments

∼1 week post injection and ferrule implants, ChR2- and GFP-expressing mice were trained sequentially on each behavioral assay. Before each behavioral session, optic ferrules on the animal’s head were coupled to patch cords (Product ID: SBP, Doric) attached to a commutator (Product ID: FRJ_1×1_FC, Doric) using ceramic split mating sleeves (Product ID: ADAF, ThorLabs). The commutator was connected to the 473 nm laser (Product ID: BL473T8-200FC, SLOC Lasers) via a patch cable (Product ID: M72L05, ThorLabs). Mice were tethered during behavioral training to habituate them to the patch cables. Mice were run on the two choice operant assay for five days on the partial water access schedule followed by five days on the full water access schedule. Optogenetic activation experiments were then performed over seven consecutive behavioral sessions. During each one-hour behavioral session, mice were stimulated for 8s during the reward period on a random 50% of trials (473 nm, 20 Hz, 5 ms pulse duration, 1-3 mW at fiber tip). BPOD Sanworks and a custom MATLAB program were used to control laser pulses. Behavioral analysis was performed using Bonsai and custom MATLAB software to compare behavioral task parameters, such as reward latency, between (non-stim versus stim trials) and across (GFP control versus ChR2 experimental) mice.

### Optogenetic Inhibition Experiments

∼1 week post injection and ferrule implants, GtACR- and mCherry-expressing mice were trained sequentially on each behavioral assay. Before each behavioral session, optic ferrules on the animal’s head were coupled to patch cords (Product ID: SBP, Doric) attached to a commutator (Product ID: FRJ_1×1_FC, Doric) using ceramic split mating sleeves (Product ID: ADAF, ThorLabs). The commutator was connected to the 473 nm laser (Product ID: BL473T8-200FC, SLOC Lasers) via a patch cable (Product ID: M72L05, ThorLabs). Mice were tethered during behavioral training to habituate them to the patch cables. Mice were run on the two choice operant assay for five days on the partial water access schedule followed by five days on the full water access schedule. Optogenetic activation experiments were then performed over seven consecutive behavioral sessions. During each one-hour behavioral session, mice were stimulated for 8s during the reward period on a random 50% of trials (473 nm, 8s pulse duration, 10 mW at fiber tip). BPOD Sanworks and a custom MATLAB program were used to control laser pulses. Behavioral analysis was performed using Bonsai and custom MATLAB software to compare behavioral task parameters, such as reward latency, between (non-stim versus stim trials) and across (mCherry control versus GtACR experimental) mice.

### Analysis of Optogenetic Experiments

We tracked each mouse’s position from the time of choice port entry to the end of the trial (social trials: from nose-poke to 20s after; sucrose trials: from nose-poke to 8s after) using its centroid coordinates obtained through Bonsai analysis. With these coordinates, we created a 2-D distribution (256 x 256 pixels) of each mouse’s exploration of the chamber. To quantify the distance traveled in centimeters during the reward period per trial (Figure 5f,j,p,t and Supplementary Figure 11d,i,n,s), we identified the number of non-zero pixels from the time of choice port entry to the end of the trial and scaled the size of each pixel to its size in centimeters. To calculate the time spent in the social zone per social trial (Figure 5g,k,q,u and Supplementary Figure 11e,j,o,t), we used Bonsai tracking analysis to identify the mouse’s entry into the social reward zone from the time of choice port entry to the end of the trial (20 secs after social choice port entry). To control for locomotor changes with mPFC activation, mice were stimulated for up to 60s in the operant arena outside of the reward period. The distance traveled (cm) outside of the reward period was quantified in 5s bins for the duration of the stimulation as described above (Supplementary Figure 11f,k,p,u).

### Histology and Microscopy

After all imaging and optogenetic experiments were completed, mice were sacrificed and perfused with 0.5% PBS followed by 4% paraformaldehyde. The brains were dissected out and fixed in 4% PFA overnight at 4°F. They were then transferred to 30% sucrose solution and allowed to fix for a minimum of 24 hours. To visualize viral expression and GRIN lens/ferrule placement, brains were sliced at 50µm slices using an Epredia HM430 microtome and the slices containing the target brain regions were mounted in DAPI solution (Product ID: 0100-20, Southern Biotech). Slices were imaged using fluorescence microscopy (Keyence BZ-X800). Images were registered using WholeBrain software^84^ to determine GRIN lens and ferrule placement.

### Data Analysis and Statistics

#### Task modulation analysis

We classified neurons on whether they were significantly modulated by various task events (trial start, social/sucrose choice, social/sucrose reward). To do this, we examined the normalized change in fluorescence in each peri-behavioral task window. Given the temporal proximity of task events (especially choice and reward delivery), we designated a 3s window around each task event and calculated the mean fluorescence within that window. Peri-behavioral task windows were assigned as follows to eliminate overlap between task event responses: trial start: 1.5s before and after; choice: 2.5s before 0.5s after; reward: 0.5s before and 2.5s after. For excitatory responses, we compared the maximum value of the mean fluorescence to the maximum value expected by a shuffled distribution (>99.5 percentile). For inhibitory responses, which require a sustained suppression of fluorescence, we compared the mean value of fluorescence across the time window to that expected by a shuffled distribution (<0.5 percentile). To determine if the percent of non-overlapping reward-responsive neurons observed in male and female mice (Figure 4e) was greater than chance, we compared the proportion of non-overlapping reward responses found in male and female mice to a shuffled distribution (generated by shuffling labels 100 times). All statistical tests were performed using custom MATLAB scripts.

#### Tracking neurons across imaging sessions

We imaged mPFC neurons on the two choice operant assay during three water access conditions (two full water access and one restricted water access conditions) and tracked these neurons across imaging sessions as described in Sheintuch et al, 2017^50^. The average time between imaging sessions in male mice was as follows: FWvRW: 8.13 ± 0.72 days, FWvFW2: 21.29 ± 1.13 days and FW2vRW: 12.14 ± 0.86 days. The average time between imaging sessions in female mice was as follows: FWvRW: 10.80 ± 0.92 days, FWvFW2: 4.0 ± 0.84 days and FW2vRW: 7.0 ± 0.32 days. We then classified neurons based on their reward responses (social reward excite, sucrose reward excite, sucrose reward inhibit or reward unresponsive) and compared their reward response profiles across similar (full water access and full water access 2) and dissimilar (full water access/full water access 2 and restricted water access) water access conditions.

#### Encoding model and task modulation analysis

To isolate the contribution of each behavioral event (trial start, social choice, sucrose choice, social reward, sucrose reward) to the variation in the fluorescence activity, we used a linear encoding model as previously described to estimate neural response kernels for every neuron^21,22^ (Supplementary Figure 8). We modeled the z-scored fluorescence activity of each neuron (chosen as the *dependent variable*) as a consequence of a behavioral event (chosen as the *independent variable*), which was designed to be a set of binary arrays of the behavioral event time (1 when the event occurs, 0 otherwise), convolved with an spline basis set (with 10 splines) spanning 3 seconds. Thus, the independent variables of the multiple-linear regression model capture time-delayed relationships between the behavioral event and the corresponding fluorescence. Mathematically, the regression model is set up as below –

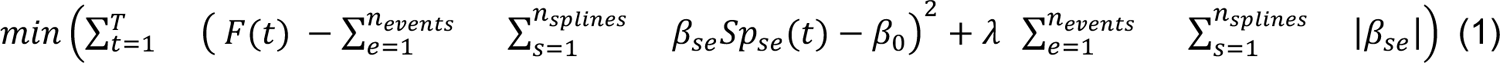

where, *T* is the total duration of the recording, *F(t)* is the z-scored fluorescence of the neuron at time *t*, *n_events_* is the number of behavioral events being modeled, and *n_splines_* is the number of degrees of freedom used for the spline basis set (*n_splines_* = 10). The regression parameters β_se_ and β_0_ are the coefficients for the s^th^ spline for event e and intercepts respectively. Lasso regularization parameter λ is identified through minimizing the squared distance between the fluorescence and the modeled data, using five fold cross-validation. *Sp*_se_(*t*) is the independent variable of the regression, and is calculated as follows –

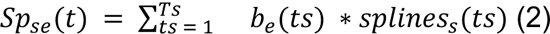

where, *ts* samples from 1 to *Ts* = 3 seconds, *b*_e_(*ts*) is the binary array for the event e, and *splines*_s_ is the s^th^ bspline basis set, created in MATLAB using the *create_bspline_basis* function of the MATLAB package fdasrvf (https://github.com/jdtuck/fdasrvf_MATLAB).

Considering the temporal proximity of the events, we chose three second windows for the choice events (‘Sucrose choice’/’Social choice’) as 2.5 seconds before, and 0.5 seconds after the event; reward events (‘Sucrose reward’/’Social reward’) as 0.5 seconds before, and 2.5 seconds after; and ‘Trial start’ to be 1.5 seconds before and after the event.

Finally, using the regression parameters β_se_ and β_0_, we calculated the kernel for each behavioral event and neuron as

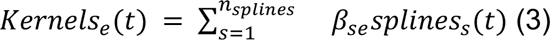

We used this encoding model to identify neurons that significantly encode a behavioral event. To identify significant neurons, we compared the change in the goodness of fit of our encoding model (Eq. 1) to the fluorescence data calculated with all events (full model, with n_events_ = N) to a reduced model where one of the events is removed (n_events_ = N-1). We used F-statistic as a measure of goodness of fit for the models and iteratively calculated this statistic by removing predictors of each event from the model. We then compared this value with 500 instances of shuffled data, obtained by shuffling the fluorescence data in 1-second bins. We obtained a p-value by comparing the F-statistic from the real data to the shuffled data. If the calculated p-value for the response of a neuron to the event is less than 0.01 (account for multiple comparisons), the neuron is considered to encode the event.

#### Neural trajectories for choice and reward

To determine the population activity trajectories for sucrose and social trials, we considered all recorded neurons across trials and reduced them to a three-dimensional neural subspace using Principal Component Analysis (PCA)^85^. For visualizing the male and female choice (Figures 3k,m) and reward (Figures 4s,t) trajectories, we calculated trial-averaged z-scored fluorescence chosen around choice or reward time windows respectively and concatenated across animals to form a three-dimensional array as [*number of neurons x trial type x time window*]. We fit a PCA (coded in MATLAB using *pca*) to the array after collapsing across the last two dimensions and projected the resulting dimensionality-reduced array independently onto social and sucrose trials (*score* output from *pca*). To quantify the difference between the neural subspace occupied by these trials (Figures 3l,n; 4s,t) we repeated the procedure as above for each animal and calculated pairwise Euclidean distance both within and between the first three PC projected vectors of social and sucrose trials^31^. Average Euclidean distances (Figures 3l,n; 4s,t) are reported across 9 male mice and 6 female mice. Additionally, in Figure 4s,t, we identified the contribution of social and sucrose reward neurons to the variability explained by PC1 as compared to the rest of the mPFC population. Average PC1 coefficients (absolute values for the loading of PC1, *coeff* output from *pca*) are reported across all neurons identified as significantly encoding social (134 neurons in male, 202 neurons in female), sucrose reward (88 neurons in male, 107 neurons in female) or neither (283 neurons in male, 298 neurons in female). In Figure 6n,o, we determined if neural subspace occupied by social and sucrose trials varied with changes in the internal state of the animal. We calculated trial-averaged z-scored fluorescence chosen around reward from neurons tracked across both the FW and RW access conditions (see section <u>Tracking neurons across</u> <u>imaging sessions)</u> that were concatenated, dimensionality-reduced, and projected independently onto social and sucrose, FW and RW trials as described previously. We also quantified the euclidean distances both between and within (Figure 6n,o) the PC projected vectors of the above four trial types and reported the average across 8 male mice and 5 female mice.

#### Decoding choice and reward using neural data

To decode the choice made by the mouse (Figure 3h) in the current trial or the identity of the reward consumed during the trial (Figure 4p), we used logistic regression (coded in MATLAB using *fitclinear*) for the neural data from the corresponding choice or reward time window. For each mouse, we used trial-matched numbers of sucrose and social trials to construct a three-dimensional array of [*number of neurons x number of trials x time window of z-scored fluorescence data*]. We then collapsed across the last two dimensions to use as an input to the regression. This was done to maximize the number of samples that could be used to train our decoder^86^. We used a 70-30, train-test split in our dataset to train the decoder and reported the test decoding accuracy as an average of 500 iterative splits of the dataset. We also calculated the decoding accuracy of a shuffled dataset, obtained by shuffling the z-scored data in 1 second bins. The neuron was considered to significantly decode the choice made or reward being consumed in the current trial if averaged test data accuracy was significantly higher than when compared with the accuracy of a shuffled dataset using a two-tailed t test. Average accuracy is reported across 9 male mice and 5 female mice (1 female mouse was excluded due to insufficient number of sucrose trials for splitting into training and test datasets).

#### Time-course of choice decoding

To determine the earliest time point from which neural activity can successfully decode the choice made on the current trial (social or sucrose), we modified the logistic regression model previously described as follows. The input to the decoder was the z-scored fluorescence with a time-window of 6 seconds prior to the nose poke, and 2 seconds after. For trials in which choice latency was <6 seconds, we time matched by linearly interpolating the neural data from trial start to nose poke. This data was then binned into non-overlapping 500ms bins and decoding accuracy was independently calculated for each time bin. This process was repeated for shuffled data, and significant choice decoding was identified using a two-tailed t test when the decoding accuracy in the time bin of the original data was significantly different from the shuffled data. To compare the time-course of choice decoding between males and females (Figure 3g), we accounted for the differences in the trial numbers and the number of animals in each group. We chose an equal number of animals from each sex and matched the number of sucrose and social trials both between and within the groups.

#### Sex decoding

To determine if the neural activity around the choice and reward periods decode the sex identity of the mice, we applied the logistic regression model as previously described as follows. For each decoder, we input the corresponding z-scored fluorescence around either a social or sucrose trial from a pair of male and female mice, matched for the number of trials and neurons recorded from. Across the 8 male and 5 female mice (1 male and 1 female mouse were excluded due to insufficient number of trials for comparison), we calculated pairwise decoding accuracies across both unshuffled and shuffled data and reported the average accuracy across the 40 comparisons made either during social and sucrose choice (Figure 3i) or reward (Figure 4o).

#### Mahalanobis distance

To identify if neural responses to social and sucrose rewards are uniquely represented at a population level, we calculated the Mahalanobis distance between the reward responses of all neurons to their baseline. In line with previous studies^27,31,87^, we chose the Mahalanobis distance to quantify the difference introduced in the population response due to our two rewarding conditions, while accounting for the underlying baseline variability in the neurons^88^. For each trial type of social or sucrose, we calculated the Mahalanobis distance as:

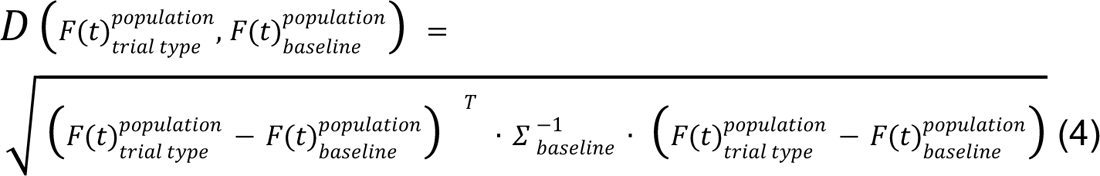

where at time t, *F*(*t*)_trial type_^population^ is the population response vector (number of neurons x 1), *F*(*t*)_baseline_^population^ is the average population response vector at baseline, and Σ_baseline_^-1^ is the covariance matrix calculated over all baseline timepoints. We calculated baseline responses for each mouse as the fluorescence average over the inter-trial-interval period of 30 secs and time-matched to the duration of the reward. Mahalanobis distances are calculated as averages over all social and sucrose trials across male (Figure 4q, 101 social and 103 sucrose trials, 9 mice) and female mice (Figure 4r, 104 social and 83 sucrose, 5 female mice) from 3 seconds before to 5 seconds after social/sucrose reward and baseline. Boxplots on the right panel are average distances for a 3 second window around the reward.

